# TIPP3 and TIPP3-fast: Improved Abundance Profiling in Metagenomics

**DOI:** 10.1101/2024.10.28.620576

**Authors:** Chengze Shen, Eleanor Wedell, Mihai Pop, Tandy Warnow

## Abstract

We present TIPP3 and TIPP3-fast, new tools for abundance profiling in metagenomic datasets. Like its predecessor, TIPP2, the TIPP3 pipeline uses a maximum likelihood approach to place reads into labeled taxonomies using marker genes, but it achieves superior accuracy to TIPP2 by enabling the use of much larger taxonomies through improved algorithmic techniques. We show that TIPP3 outperforms leading methods for abundance profiling in two important contexts: when reads come from genomes not already in a public database (i.e., novel genomes) and when reads contain sequencing errors. We also show that TIPP3-fast has slightly lower accuracy than TIPP3, but is still more accurate than other leading methods and uses a small fraction of TIPP3’s runtime. Additionally, we highlight the potential benefits of restricting abundance profiling methods to those reads that map to marker genes (i.e., using a filtered marker-gene based analysis), which we show typically improves accuracy. TIPP3 is freely available at https://github.com/c5shen/TIPP3.

**Author summary:** TIPP3 is a new marker gene-based abundance profiling tool that builds on TIPP and TIPP2 with significant enhancements. TIPP3 supports larger reference packages (∼ 55,000 sequences per marker gene) and achieves higher accuracy in abundance profiling, especially with challenging input reads containing sequencing errors or novel genomes. TIPP3 outperforms TIPP2 and other leading methods in profiling accuracy, and its fast version TIPP3-fast is competitive in runtime with the competing methods while being more accurate under challenging conditions. TIPP3 is open-source and available at https://github.com/c5shen/TIPP3.

## Introduction

Understanding the complex interactions between microorganisms in their communities has become critical to human and environmental health studies [1–4]. Typically, the first step in understanding these interactions (i.e., microbiome analysis) is taxonomic profiling, which identifies and quantifies the relative abundance of species in a microbial community.

Some studies estimate abundance profiles using amplified 16S ribosomal RNA present in prokaryotic species [5], which is cost-effective but can have quantification errors introduced by copy number variation [6]. As sequencing costs decrease continuously, newer methods for taxonomic profiling increasingly use metagenomic data, which consists of DNA reads sequenced directly from the target microbial environment, capturing millions of sequences from all genomes in the community.

Current methods for taxonomic profiling vary in their approach to using metagenomic data. Methods such as Kraken [7] and Kraken2 [8] query and map k-mers of all input reads for classification in its database, which consists of sequenced microbial genomes. Other methods, including MetaPhyler [9], MetaPhlAn [10, 11], mOTUs [12], TIPP [13], and TIPP2 [14], use marker genes to assign reads to microbial clades for classification and abundance profiling, since marker genes are single-copy and universal in Bacteria and Archaea species. The reads that are classified by these methods are only those that have been assigned to a particular marker gene. Thus, the resulting estimated abundance does not need to be adjusted for genome size or copy number variation. Many methods use a reference database for read identification, but they may fail to identify reads of under-represented species. Some methods, such as TIPP [13] and TIPP2 [14], use maximum likelihood for placing reads into reference taxonomic trees of marker genes for classification. Phylogenetic placement enables the detection of distant homologs to reference sequences, allowing characterizations of highly diverse metagenomic reads [15]. TIPP2 differs from TIPP mainly by having denser taxon sampling for each marker gene, which results in improved accuracy. However, the way TIPP2 uses pplacer [16], the phylogenetic placement method, restricts its usage to trees with at most 10,000 sequences, and thus TIPP and TIPP2 are not scalable to large taxonomic trees [17].

In just the last few years, new approaches have enabled phylogenetic placement methods to place into much larger reference trees [17–21]. Given the improvement in accuracy obtained by TIPP2 over TIPP as a result of using a slightly larger taxonomic tree for each marker gene, we hypothesize that these improved phylogenetic placement methods could potentially lead to further improvements in abundance profiling accuracy.

In this study, we present TIPP3, an updated version of TIPP2. TIPP3 builds on TIPP2 and has a more extensively built reference package with 38 marker genes and more than 50,000 sequences per gene. TIPP3 also leverages the recent developments in more accurate multiple sequence alignment methods and scalable phylogenetic placement methods. We show empirically that TIPP3 is more accurate than TIPP2 for abundance profiling, particularly for lower taxonomic levels such as species, genus, and family. Compared to other leading profiling methods, TIPP3 is the most accurate under most conditions, especially for long reads with higher sequencing error (e.g., PacBio) and for reads from novel genomes.

We also introduce TIPP3-fast, a slightly less accurate but much faster version of TIPP3, that is competitive in runtime with the other methods while being more accurate under challenging conditions.

In addition, we demonstrate that filtering input reads to only those that map to marker genes improves the profiling accuracy of Kraken2 and Bracken [22], two other leading abundance profiling methods, under most conditions, but that TIPP3 maintains an accuracy advantage over these methods for challenging conditions. Overall, we demonstrate that TIPP3 and TIPP3-fast are two valuable new additions to abundance profiling tools.

## Materials and Methods

Here we describe the materials and methods used in our study; for additional details, see Supplementary Materials, Sections S1–S4.

### The TIPP pipeline

TIPP3 and its fast version, TIPP3-fast, both use the same basic pipeline structure as TIPP2 [14], but differ in how the specific steps are performed in order to obtain improved accuracy. We begin with a high-level description of the common pipeline structure (see Fig 1), and then describe how TIPP3 and TIPP3-fast differ from TIPP2.

**Fig 1.**
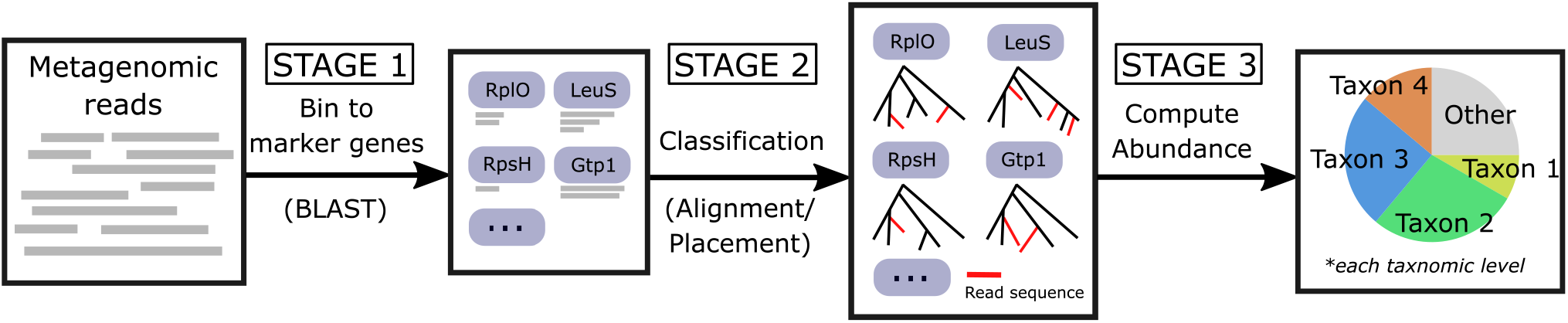
Overview of the TIPP pipeline. TIPP3 follows the same pipeline structure as TIPP and TIPP2, but differs in how some steps are performed in order to achieve higher accuracy and scalability. The common pipeline structure has three stages. Stage 1: Metagenomic reads are first binned to marker genes with BLAST. Stage 2: The query reads are added to the selected marker gene’s multiple sequence alignment and a phylogenetic placement method is used to place reads into corresponding taxonomic trees using these alignments. Stage 3: Taxonomic labels are inferred from the placements and aggregated for the final abundance profile computation.

Prior to running the method, a reference package consisting of a large set of marker genes with both alignments and taxonomic trees is constructed. The sequences from the marker genes are aggregated together to create a BLAST database for binning reads. Input reads are binned to marker gene sequences in the reference package, with a threshold of at least 50bp coverage. Then, binned reads are added to their corresponding marker gene multiple sequence alignments (MSAs) and placed into marker gene taxonomic trees. The placement within the taxonomic tree specifies some (perhaps all) of the taxonomic labels for the read, but only taxonomic levels with placement support above the user-selected threshold are considered. For example, if a read has 80% support at the species level and 98% support at the genus level and we use a support threshold of 95%, the read will be classified only at the genus level and higher. Then, the classification results are aggregated to form the final abundance profile.

We now briefly explain each stage in TIPP3 and how it differs from TIPP2.

#### Stage 1: Read binning

BLAST [23] is used to bin input reads to their corresponding marker gene sequences (≥ 50bp coverage). If a read does not map to any marker gene sequence, then it is discarded from further analysis.

#### Stage 2: Read classification

This stage can be broken into two sub-stages. We first use WITCH [24], a new method for adding sequences into MSAs, to add reads to the marker gene MSAs that they map to. Then, the query sequences are added into the relevant taxonomic tree using an improved technique for running pplacer, where it is run with the `taxtastic` package [25] (pplacer-taxtastic), which allows it to place reads into large taxonomic trees (up to 100,000 leaves). When speed is an objective, we use instead BATCH-SCAMPP [21], another maximum likelihood phylogenetic placement method that uses a divide-and-conquer approach method and takes advantage of the sublinear scaling of the phylogenetic placement method EPA-ng [15]. We use a support value of 90% for pplacer-taxtastic and 95% for BATCH-SCAMPP, and assign taxonomic labels only at those levels that achieve at least the corresponding support values.

#### Stage 3: Abundance profile computation

After reads are placed and classified from Stage 2, an abundance profile can be computed by pooling all read classifications. The relative abundance is computed as the total number of reads classified within a taxon divided by the total number of reads classified.

### TIPP3 vs. TIPP2

TIPP3 uses the same high-level algorithmic structure and the same techniques for Stage 1, but differs in the reference package construction (which is a precursor to the pipeline) and in Stage 2 (taxonomic classification of those reads that map to marker genes). Here we describe the differences.

### Reference package construction

TIPP3 uses a slightly different set of marker genes than TIPP2 (see Supplementary Materials Section S5.3 for details). TIPP3 utilizes an updated NCBI taxonomy [26] and a much larger reference package than TIPP2, increasing the number of sequences per marker gene from ∼ 4300 for TIPP2 to more than 50,000 for TIPP3. TIPP2 used PASTA [27] to compute the marker gene alignments, but TIPP3 uses MAGUS [28], which is a more accurate multiple sequence alignment method. MAGUS and PASTA both use a divide-and-conquer approach for aligning subsets of sequences, but MAGUS uses a more sophisticated technique for merging subset alignments compared to PASTA and produces more accurate multiple sequence alignments. TIPP2 used RAxML [29] to compute the taxonomic trees for their marker gene alignments and TIPP3 uses RAxML-ng [30], but both use the NCBI taxonomy as the constraint tree.

### Aligning and placing reads to marker genes

While both TIPP2 and TIPP3 use BLAST to bin reads to marker genes, they use different techniques to add the reads to the marker gene MSAs and taxonomic trees. TIPP2 uses UPP [31] to add reads to the MSAs and TIPP3 uses WITCH [24], which is more accurate. WITCH and UPP are two methods for adding sequences into a multiple sequence alignment; they are similar in their initial algorithmic design, in that they both represent the marker gene MSA using an ensemble of hidden Markov Models (HMMs), but then they differ in how the ensemble is used to add each query sequence. In UPP, each query sequence picks a single HMM in the ensemble based on the bit score, and the alignment of the query sequence to the single HMM then determines how the query sequence is added to the marker gene alignment. WITCH, in contrast, lets each query sequence pick the top few HMMs in the ensemble based on a corrected bit score, and then combines the resultant MSAs into a single MSA using a weighted ensemble approach. As shown in [24], the WITCH approach produces more accurate alignments than the UPP approach. When speed is important, TIPP3-fast, the fast version of TIPP3, uses BLAST to add reads to the marker gene alignment.

TIPP2 used pplacer for phylogenetic placement with RAxML numeric parameters, and TIPP3 uses new phylogenetic placement methods that can scale to larger reference trees. Specifically, TIPP3 uses pplacer-taxtastic [17], which uses the Python package `taxtastic` [25] for the numeric parameters; interestingly, this allows pplacer to place into trees with ∼100,000 leaves [17]. When speed is important, TIPP3-fast, the fast version of TIPP3, uses BATCH-SCAMPP (BSCAMPP), which is much faster than pplacer-taxtastic, and can scale to even larger trees.

### Abundance profiling methods

We compared TIPP3 to Bracken [22], Kraken2 [8], and mOTUsv3 [32] for abundance profiling accuracy on our testing data. Kraken2 is designed for sequence classification, and Bracken is intended to build abundance profiles based on Kraken2 outputs. Kraken2 is kmer-based and uses a large pre-built database to map reads to their lowest common ancestor taxon [7]. Bracken uses the output from Kraken2 classification and information about genomes in the database to estimate abundance at the species level and above. mOTUsv3 is a marker gene-based abundance profiling method and maps metagenomic reads to their corresponding marker gene cluster units in its database [12]. mOTUsv3 is designed for short reads, thus needing data pre-processing to deal with PacBio long reads. As recommended in the mOTUs github page, we used the “long read” option provided in mOTUs (starting with version 2) to break each long read into multiple short read segments and used the generated mock short reads for abundance profiling. We did not include a comparison to MetaPhlAn [11] since it was reported to have a high error in the TIPP2 study [14]. MetaPhyler [9] was excluded because it is no longer actively developed or maintained.

Additionally, we compared TIPP3 to alternative abundance profiling pipelines by substituting the reads phylogenetic placement method used by TIPP3 with APPLES-2 [19] and App-SpaM [33]. These two methods are not designed for abundance profiling of metagenomic data but can perform read placement into marker gene taxonomic trees in Stage 2 of TIPP3. APPLES-2 is a distance-based method that can place sequences into very large trees (up to 200,000 leaves) [19]. App-SpaM is an alignment-free placement method designed for placing short sequences into an existing tree, based on their phylogenetic distances to sequences in the tree [33].

### Datasets and read generation

For the training experiment, we used two TIPP2 datasets for read generation, one with 51 genomes and the other with 33 genomes. For the testing experiment, we used genomes from all three TIPP2 datasets and additional genomes queried from NCBI to create three new datasets for read generation. The three testing datasets are denoted as “known”, “mixed”, and “novel” based on whether the genomes included are present in the TIPP3 reference package. The “known” dataset has 50 known genomes, the “mixed” dataset has 53 known and 47 novel genomes, and the “novel” dataset has 50 novel genomes. Genomes in the three testing datasets are disjoint from each other. For a fair comparison to Bracken and Kraken2, we made sure that known genomes are also present in the Bracken/Kraken2 database. When a genome is novel to TIPP3, it is also not present in the Bracken/Kraken2 database. We used the “PlusPF” Kraken2 database published in June 2023 [34], which has the closest date to the NCBI taxonomy used for TIPP3. The composition of all datasets and their included genomes can be found in the Supplementary Materials, Section S3.

For each set of genomes, we simulated two types of reads that resemble Illumina (150bp) or PacBio (∼ 3000bp) reads. We used the ART sequence simulator [35] for Illumina reads and PBSIM [36] for PacBio reads generation. We show the properties of the simulated reads for each dataset in Table 1, and more details for read generation can be found in Supplementary Materials, Section S3.

**Table 1.**
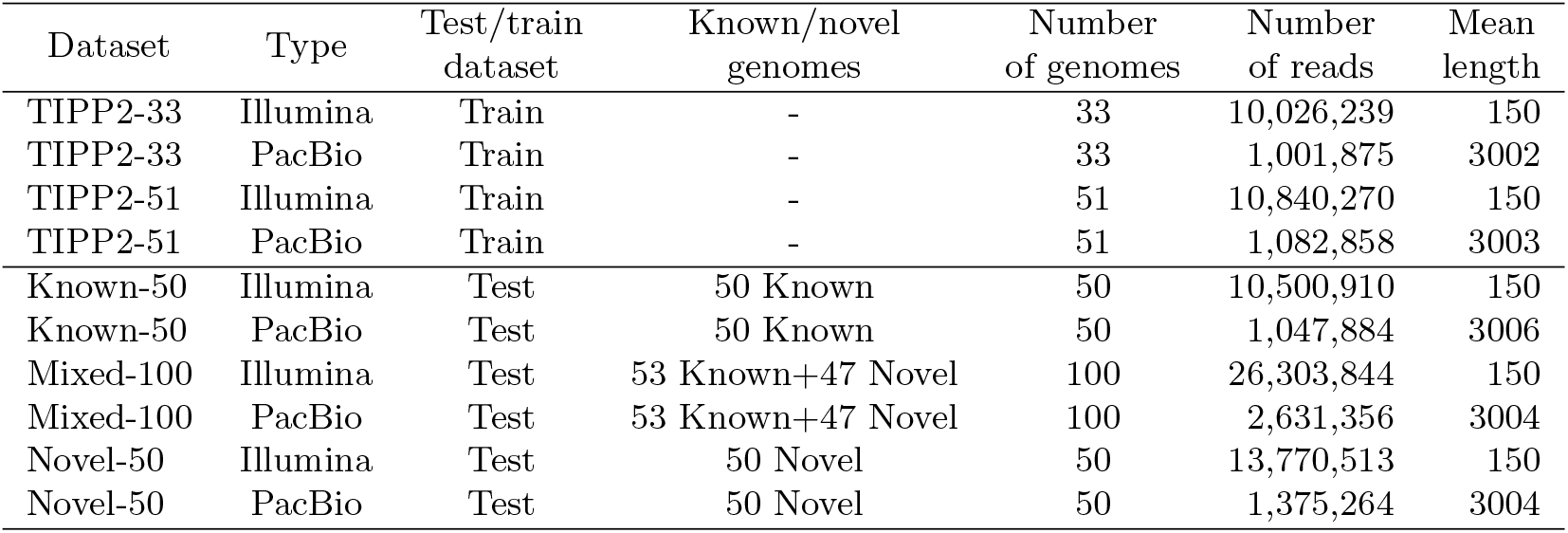
Properties of simulated reads.

| Dataset | Type | Test/train dataset | Known/novel genomes | Number of genomes | Number of reads | Mean length |
| --- | --- | --- | --- | --- | --- | --- |
| TIPP2-33 | Illumina | Train | - | 33 | 10,026,239 | 150 |
| TIPP2-33 | PacBio | Train | - | 33 | 1,001,875 | 3002 |
| TIPP2-51 | Illumina | Train | - | 51 | 10,840,270 | 150 |
| TIPP2-51 | PacBio | Train | - | 51 | 1,082,858 | 3003 |
| Known-50 | Illumina | Test | 50 Known | 50 | 10,500,910 | 150 |
| Known-50 | PacBio | Test | 50 Known | 50 | 1,047,884 | 3006 |
| Mixed-100 | Illumina | Test | 53 Known+47 Novel | 100 | 26,303,844 | 150 |
| Mixed-100 | PacBio | Test | 53 Known+47 Novel | 100 | 2,631,356 | 3004 |
| Novel-50 | Illumina | Test | 50 Novel | 50 | 13,770,513 | 150 |
| Novel-50 | PacBio | Test | 50 Novel | 50 | 1,375,264 | 3004 |

### Evaluation criteria

#### Normalized Hellinger distance

TIPP and TIPP2 used the Hellinger distance [37] to measure the abundance profiling error of methods. Given a set of reads, the Hellinger distance of an estimated abundance profile to the true abundance profile on a taxonomic level (e.g., at the species level) is given by:

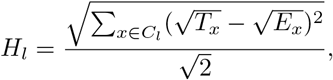

where *T*_*x*_ is the true abundance and *E*_*x*_ the estimated abundance of a clade *x*, for each *x* in the set of clades *C*_*l*_ on a taxonomic level *l*. Reads that are unclassified at a certain level are not counted for the Hellinger distance calculation.

However, in certain cases, *H*_*l*_ does not correctly reflect the actual profiling error of a method. Here, we present a new measurement, **Normalized Hellinger distance**, 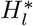,that provides unbiased measurements of estimated profiles in all cases. New variables included in the modified equation are *n*, the total number of reads classified, and *n*_*l*_, the number of reads classified at taxonomic level *l*. See Section S4 for the full derivation of the normalized Hellinger distance and an example of when Hellinger distance is unsuited.

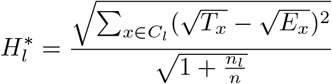

#### Additional evaluation metrics

In the training experiment, we also evaluate TIPP3 for its taxonomic identification accuracy at each taxonomic level, which is presented in terms of (1) the fraction of correctly identified reads, (2) the fraction of incorrectly identified reads, and (3) the fraction of unclassified reads. For each taxonomic level, the three values sum to 1.

We also measure the wall-clock running time and maximum memory usage. Each method is run on the University of Illinois Campus Cluster given 16 CPU cores and up to 256 GB of memory.

## Results

### Overview

We include three experiments in this study. In Experiment 1, we use the training data to set the algorithmic parameters for TIPP3. The algorithmic parameters include the alignment and phylogenetic placement methods for our binned query reads, and the set of marker genes that are used to filter reads for abundance profiling. In Experiment 2, we run Bracken and Kraken2 with two different sets of input reads: the set of all reads without any filtering, and the set of filtered reads that are binned to our marker genes in Stage 1 of TIPP3, demonstrating that filtering can improve abundance profiling accuracy. In Experiment 3, we validate the improvement of TIPP3 over TIPP2 using our testing data, and we compare TIPP3 to other leading profiling methods: Bracken, Kraken2, and mOTUsv3. We show that TIPP3 is often the most accurate abundance profiling method, especially for challenging datasets that are hard to correctly profile.

### Experiment 1: Designing TIPP3

#### TIPP3 algorithmic parameters

In this experiment, we explored different ways to run TIPP3 and decided on the most suitable TIPP3 pipeline optimizing for profiling accuracy and runtime. We explored (1) different ways to add reads to marker gene MSAs, (2) different ways to place reads into the marker gene taxonomic tree, and (3) different selections of marker genes for the aggregated abundance profile. We used Illumina and PacBio reads from two TIPP2 datasets for all training experiments and three marker genes (RplO, RpsK, and RpsL) for the abundance profile, except when exploring marker gene selection for TIPP3, where we examined all 40 marker genes. We provide a summary of the results that determined the parameters for TIPP3. Detailed experimental results can be found in the Supplementary Materials, Section S5.

#### TIPP3

Based on the training results and optimizing for the abundance profile accuracy, we selected WITCH [24] for adding reads to marker gene MSAs. For phylogenetic placement, we selected pplacer with the `taxtastic` package [25] (pplacer-taxtastic) and with a support value of 90%. Finally, we chose 38 marker genes (excluding FtsY and RpoB because of poor individual profiling results) for the aggregated abundance profile. See Supplementary Materials, Figs S1–S14.

#### TIPP3-fast

When optimizing for accuracy on our training data, we observed two bottlenecks for runtime. The first was the time used to add reads to marker gene alignments using the most accurate method tested, WITCH. The second was the read placement time required by pplacer-taxtastic, the most accurate method tested for placing reads into taxonomic trees of marker genes. To optimize speed, we design “TIPP3-fast”, which differs from TIPP3 in two ways. First, instead of using WITCH to add reads into marker gene MSAs, it uses BLAST to compute pairwise alignments of binned reads to marker genes. Second, instead of using pplacer-taxtastic, it uses BSCAMPP [21], which is a faster phylogenetic placement method that is also based on maximum likelihood; for this placement technique, it uses a support value of 95%. Although BSCAMPP has slightly lower accuracy than pplacer-taxtastic, our study shows that it is much faster and only reduces accuracy by a small amount. TIPP3-fast is also included in Experiment 3 for comparison to other abundance profiling methods on testing data.

### Experiment 2: Restricting Kraken2 and Bracken to filtered reads

Bracken and Kraken2 estimate abundances based on all input query reads. In this experiment, we examined the impact of restricting these methods to just those reads that map to the marker genes (i.e., filtering the input reads). We refer to these two different ways of running each method by appending either “(all)” or “(filtered)” to the method’s name.

Fig 2a shows the impact on profiling accuracy for filtering both methods. Filtering consistently improves accuracy for both Kraken2 and Bracken when working with Illumina reads. For PacBio reads, filtering continues to enhance Kraken2’s accuracy, but this isn’t always true for Bracken. While Bracken(filtered) outperforms Bracken(all) at the species and genus levels for PacBio reads, in some cases, it increases profiling errors compared to the unfiltered version. This issue is particularly noticeable at the order, class, and phylum levels when profiling reads from novel genomes, where Bracken(filtered) is less accurate than Bracken(all).

**Fig 2.**
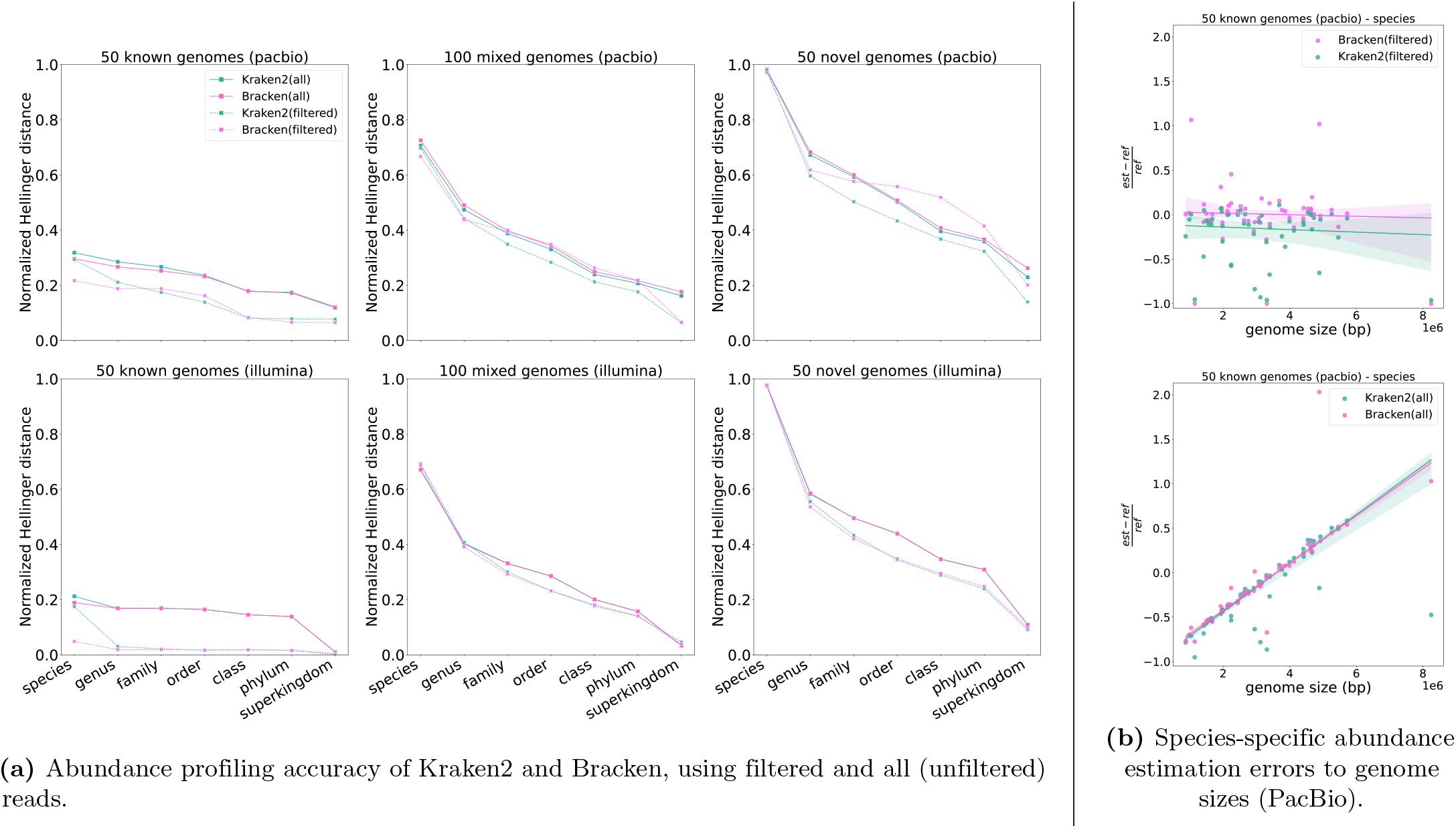
The impact of filtering on Bracken and Kraken2 for abundance profiling. Left: Abundance profiling accuracy by normalized Hellinger distance (lower means more accurate) of two ways of running Kraken2 and Bracken, for PacBio (top) and Illumina (bottom) simulated reads from 50 known, 100 mixed, and 50 novel genomes. Dashed lines correspond to using filtered reads, and solid lines correspond to using all (unfiltered) reads. Right: Scatter plot of species-specific abundance estimation errors (PacBio reads) to corresponding genome sizes for 50 known genomes of Bracken/Kraken2, using filtered (top) and all reads (bottom) as inputs. The estimation error for each taxon is calculated as the fractional difference between its estimated abundance and the reference abundance (y-axis). A Robust Linear Model with Huber Loss [38] was used to fit a regression line for each method. The shaded area around each fitted line represents a 95% confidence interval of the corresponding method.

To better understand why filtering improves accuracy for Kraken2 and Bracken, we investigated the impact of genome size on abundance profiling error on PacBio reads, when using either all the reads or only those that map to the marker gene, taken from known genomes. We plotted the fractional estimation errors for individual species against their corresponding genome sizes, and computed a Robust Linear Model with Huber Loss [38] to fit a regression line for each method, with a 95% confidence interval displayed. The results (Fig 2b) show that when using filtered reads, Bracken and Kraken2 exhibit estimation errors that are independent of genome sizes; however, when using all input reads there is a strong linear increase in error as the genome size increases.

In summary, we do see some conditions (specifically, PacBio reads from novel genomes) where filtering does not improve Bracken2 and can even reduce accuracy, but in general, filtering improves or maintains accuracy for both methods – and consistently so at the species through family levels. Since the primary focus of abundance profiling is typically on lower taxonomic levels (species and genus) and filtering improves accuracy at these levels, we present results only for the filtered versions of Bracken and Kraken2 in the remaining figures.

### Experiment 3: Evaluation of TIPP3 for abundance profiling Comparing TIPP3 to TIPP2

Using our testing datasets we now demonstrate the impact of using a larger reference package within the TIPP3 pipeline by comparing TIPP3 to “TIPP2.5”, a version of TIPP3 restricted to using the smaller reference package size of TIPP2.

We generated a new reference package for TIPP2.5 by sub-sampling the TIPP3 marker gene taxonomic trees and alignments, selecting 1-3 genomes per genus. This allows the taxonomy for each marker gene to contain ∼ 5505 sequences (∼ 4300 sequences in TIPP2).

Fig 3 compares TIPP3 to TIPP2.5 for abundance profiling accuracy using normalized Hellinger distance. For all six testing datasets, TIPP3 is more accurate than TIPP2.5. The difference between the two methods is more noticeable on lower taxonomic levels such as species, genus, and family, particularly for reads from known genomes. As we include more novel genomes in our dataset the error of both methods increases and the difference in accuracy between the two methods decreases, especially on the lower taxonomic levels such as species and genus. These trends are consistent across both Illumina- and PacBio-style reads, showing that TIPP3 improves upon TIPP2 through its more densely sampled reference package.

**Fig 3.**
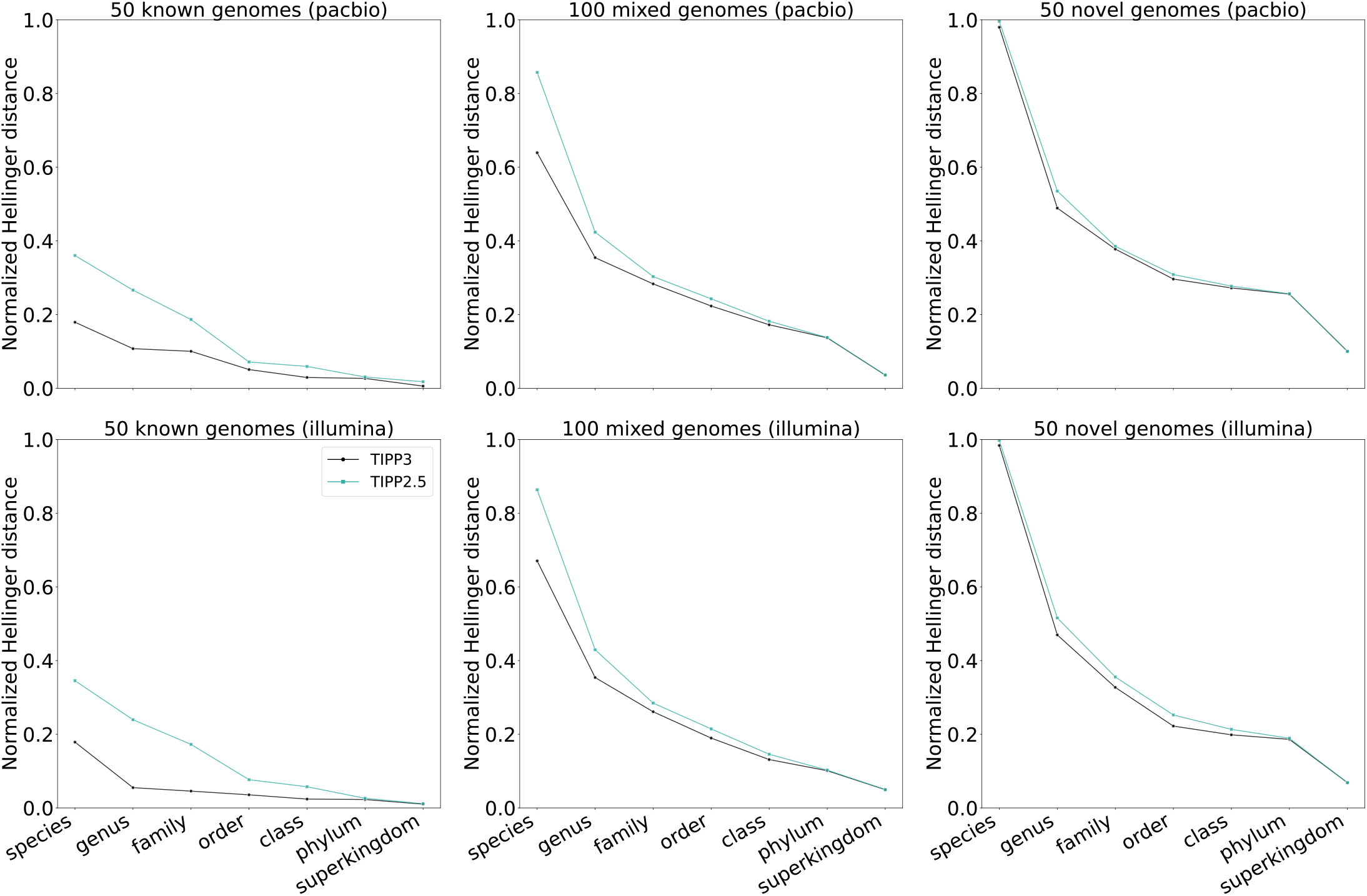
Abundance profiling accuracy by normalized Hellinger distance (lower means more accurate) of TIPP3 and TIPP2.5 for PacBio and Illumina reads from 50 known, 100 mixed, or 50 novel genomes. The abundance profile is computed using 38 marker genes. TIPP3 uses pplacer with the `taxtastic` package for placement and a support value of 90%. TIPP2.5 uses pplacer for query placement and a support value of 95%, the same setup in TIPP2 [14]. Both TIPP3 and TIPP2.5 use WITCH to add query reads to marker gene MSAs.

#### Comparing TIPP3 to other methods

We explored the impact of substituting the maximum likelihood-based phylogenetic placement methods (pplacer-taxtastic for TIPP3 and BSCAMPP for TIPP3-fast) by either a distance-based method (APPLES-2 [19]) or an alignment-free method (AppSpaM [33]). These experiments, shown in Fig S15, establish that changing the phylogenetic placement method to either APPLES2 or App-SpaM reduces accuracy. Hence, here we only show results for TIPP3 (which uses pplacer-taxtastic) and TIPP3-fast (which uses BSCAMPP).

We now compare the accuracy and runtime/memory usage of TIPP3 and TIPP3-fast to Kraken2(filtered) [8], Bracken(filtered) [22], and mOTUsv3 [32]. These experiments, shown in Fig 4, show that mOTUsv3 did not output any classification for any PacBio reads, even using the pre-processing step recommended by the authors of mOTUsv3 to deal with long reads (a strategy that is also used in [39]). Hence, results for mOTUsv3 are not shown for the three PacBio datasets.

**Fig 4.**
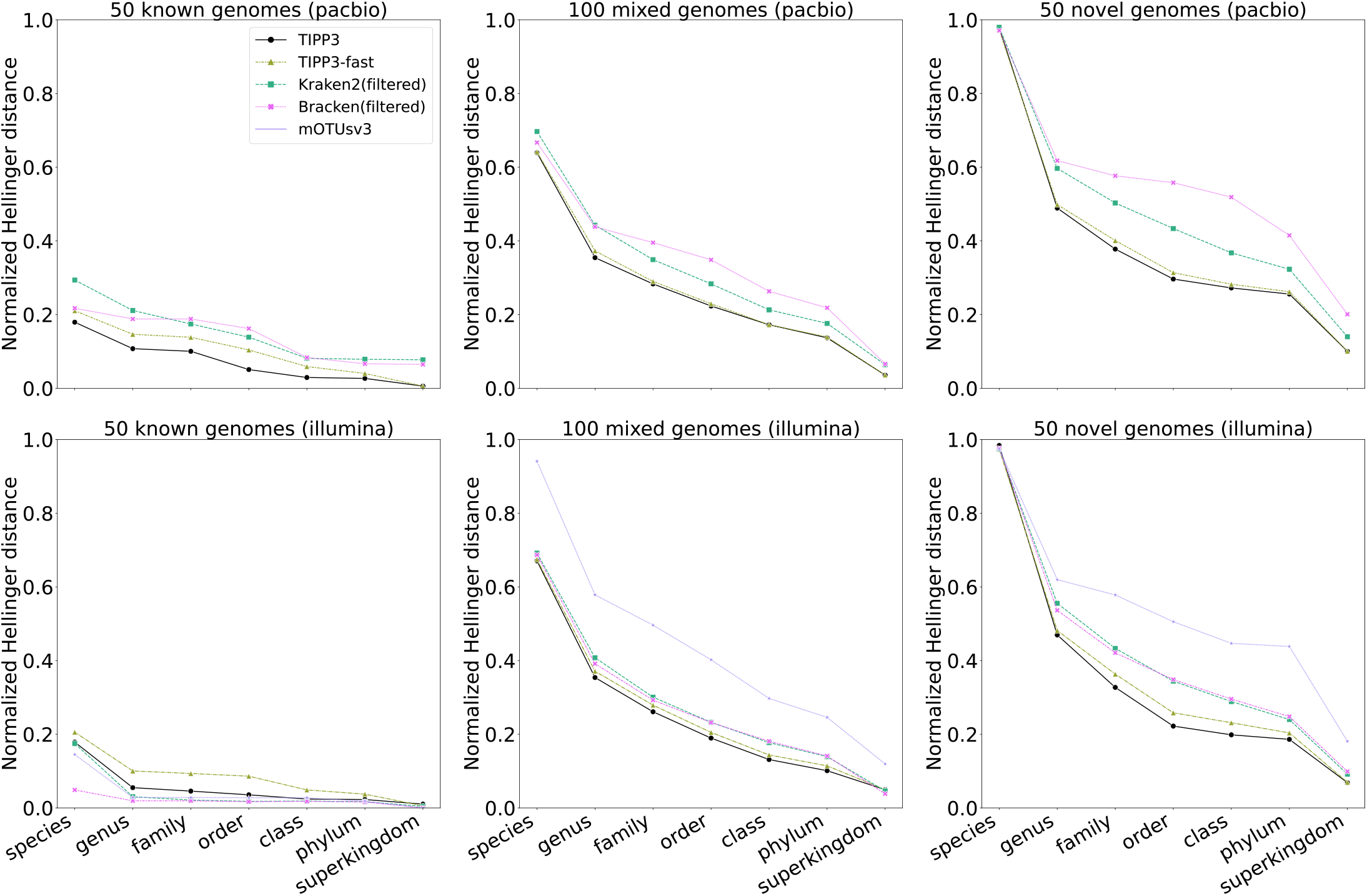
Abundance profiling accuracy by normalized Hellinger distance (lower means more accurate) of top-performing methods/pipelines for PacBio and Illumina simulated reads from 50 known, 100 mixed, and 50 novel genomes. Abundance profiles are computed using 38 marker genes for the methods except mOTUsv3 which did not utilize TIPP3 marker genes. For PacBio read datasets, mOTUsv3 did not produce any classification or profile and thus is not shown.

#### Known genomes

When analyzing PacBio long reads from 50 known genomes (Fig 4, top left), TIPP3 is the most accurate method, trailed by only TIPP3-fast. Bracken(filtered) was able to improve accuracy from Kraken2(filtered) at the species and genus levels, but its overall profiling accuracy is lower than TIPP3 and TIPP3-fast. In the case of profiling Illumina short reads from 50 known genomes (Fig 4, bottom left), Kraken2(filtered), Bracken(filtered), and mOTUsv3 are the most accurate methods, with Bracken(filtered) being much more accurate than the remaining methods at the species level estimation. On Illumina-style reads TIPP3 has lower accuracy than the three methods above at lower taxonomic levels (species, genus, family, and order) and ties at higher levels.

#### Mixed genomes

For reads from 100 mixed genomes (53 known and 47 novel genomes), the relative profiling accuracy between methods changes considerably. Most notably, all methods have much lower accuracy than when classifying reads from known genomes, particularly on the lower taxonomic levels. Furthermore, for both Illumina and PacBio reads, TIPP3 and TIPP3-fast are the most accurate methods. For PacBio reads (Fig 4, top middle) TIPP3 and TIPP3-fast have a larger accuracy advantage over Kraken2(filtered) and Bracken(filtered) than they do for the Illumina reads (Fig 4, bottom middle). Interestingly, Bracken(filtered) is less accurate than Kraken2(filtered) on PacBio reads except at the species level, opposite to the observation with Illumina reads. mOTUsv3 shows a much higher error than the other four methods on Illumina reads.

#### Novel genomes

When all reads are from novel genomes, the absolute error in terms of normalized Hellinger distance of all methods increases even further (Fig 4, top right and bottom right). All methods have errors close to 1 at the species level, which is expected since all genomes are novel. Except for the species level, TIPP3 is the most accurate method for both Illumina and PacBio reads. TIPP3-fast has a slightly higher error than TIPP3 but is more accurate than Kraken2(filtered), Bracken(filtered), and mOTUsv3. For PacBio reads, we again observe that Bracken(filtered) has lower accuracy than Kraken2(filtered), this time with a large differentiation between the two methods at higher taxonomic levels without the improvement at the species and genus levels that is seen on the known and mixed genome datasets.

#### Detailed evaluation on species abundance

We now examine abundance profiling error on a per-species basis, where the estimation error is given by 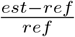 where *est* is the estimated abundance and *ref* is the true (reference) abundance of a species. We use the subset of the species for which at least one of the studied methods (TIPP3, TIPP3-fast, Kraken2(filtered), Bracken(filtered), and mOTUsv3) has abundance profiling error*±*10%. Full results for species and genus abundances can be found in Supplementary Materials, Section S6 (Figs S16–S19).

#### Illumina reads of 50 known genomes

When inspecting individual species for Illumina reads, Bracken(filtered) has nearly zero estimation error for all species abundances (Fig 5a). mOTUsv3 is the second most accurate, with only three species abundances exhibiting noticeable errors. TIPP3, TIPP3-fast, and Kraken(filtered) display a similar error composition, generally having more underestimation errors compared to Bracken(filtered) and mOTUsv3, while having few overestimation errors. These results are consistent with the relative performance shown in Fig 4.

**Fig 5.**
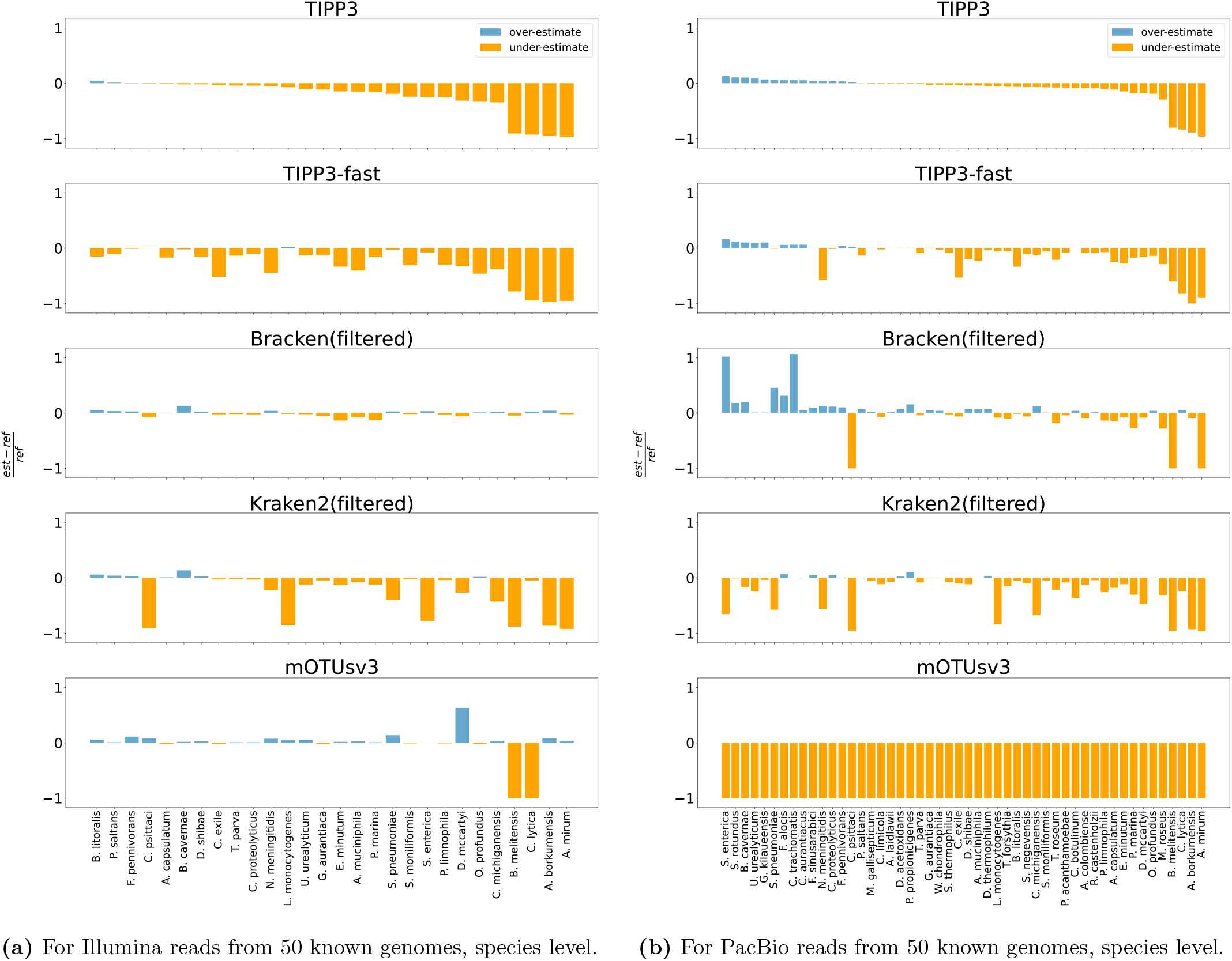
Species-specific abundance estimation error for Illumina (left) and PacBio (right) reads of 50 known genomes of methods from Fig 4. The estimation error of a species is computed as the fractional difference between its estimated abundance and the reference abundance, shown on the y-axis. For each comparison, a taxon is shown if and only if it is present in the reference and at least one method has an estimation error *±*10%. Taxa are sorted left-to-right by TIPP3’s error, from overestimation to underestimation. Full results for all datasets at species and genus levels can be found in Supplementary Section S6, including all taxon names.

#### PacBio reads of 50 known genomes

For PacBio reads (Fig 5b), TIPP3 demonstrates the highest accuracy, followed closely by TIPP3-fast, with the primary source of error for both methods being the underestimation of species abundances. Similarly, Kraken2(filtered) tends to underestimate species abundances, consistent with the observations in Fig 5a. The comparison between Bracken(filtered) and Kraken2(filtered) is more complicated; Bracken(filtered) has more overestimation errors than Kraken2(filtered), but has much fewer underestimation errors. Interestingly, according to Fig 4, Bracken(filtered) has an overall lower abundance profiling error, using the normalized Hellinger distance, than Kraken2(filtered). Finally, mOTUsv3 did not produce any profiles, and we recorded this as 100% underestimation error for all species.

#### Runtime and memory

All methods were given 16 cores of CPU and up to 256 GB of memory and allowed to run to completion. We used the University of Illinois, Urbana-Champaign Campus Cluster, which is a heterogeneous runtime environment with a mixture of old and new generations of CPUs, making the runtime comparison somewhat unreliable. Given this caveat, we report runtime and memory usage until an output abundance profile was computed, including the runtime for filtering reads for some methods (TIPP3, TIPP3-fast, Bracken(filtered), and Kraken(filtered)).

Fig 6 shows the runtime usage of TIPP3, TIPP3-fast, Kraken2(filtered), Bracken(filtered), and mOTUsv3 on the six datasets. Full results for TIPP3 using APPLES2 and App-SpaM for read placement are shown in Fig S20. All methods were able to complete each dataset without any runtime error. On all datasets, TIPP3 used the longest time (101–312 hours) to complete, with a large portion of the runtime dedicated to running WITCH to add query reads to marker gene MSAs. The other methods were able to complete each of the given testing datasets in *<*10 hours. The fastest methods are Kraken2(filtered), Bracken(filtered), and mOTUsv3, taking 0.1-3.7 hours to complete any dataset. TIPP3-fast is slightly slower at 1.0-5.2 hours.

**Fig 6.**
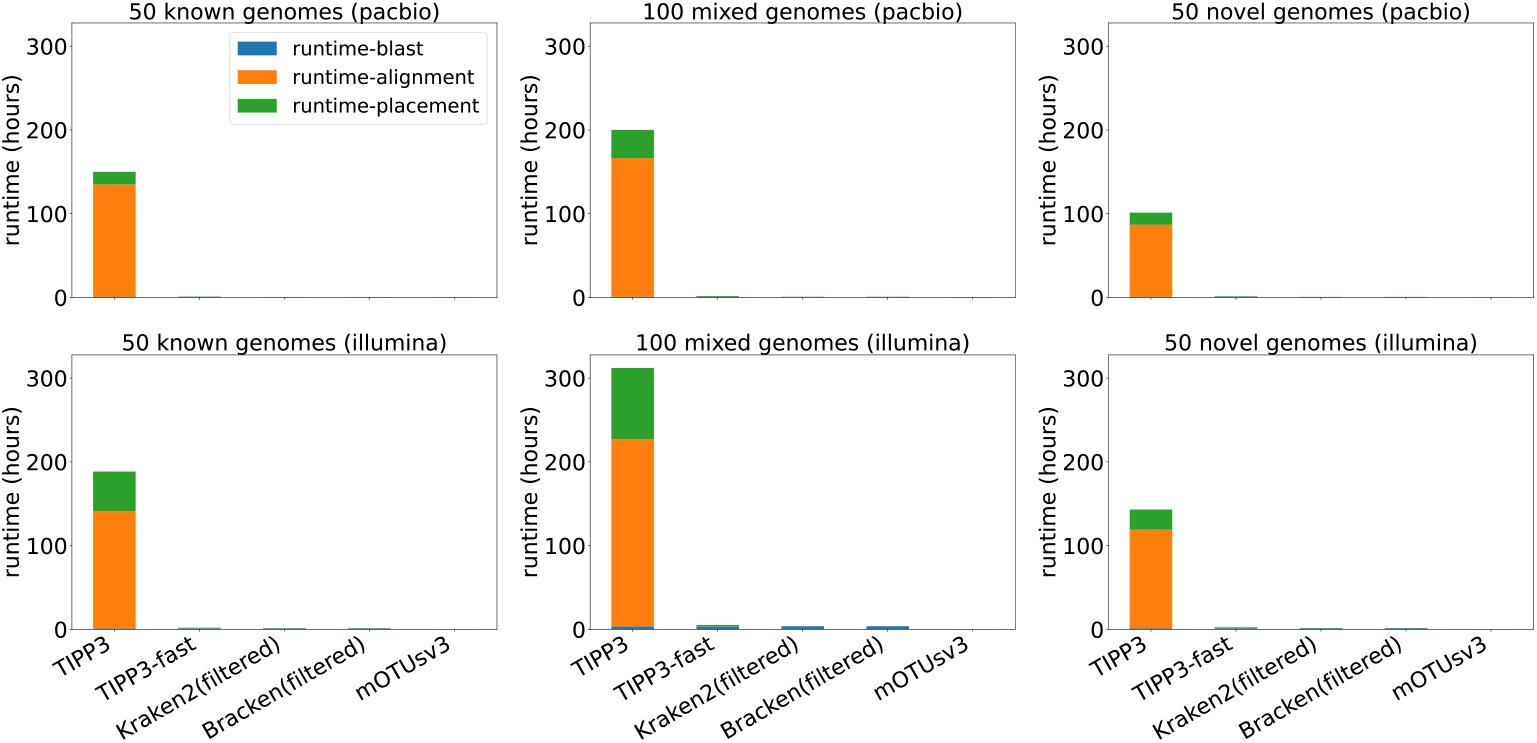
Runtime in hours of TIPP3, TIPP3-fast, Kraken2(filtered), Bracken(filtered), and mOTUsv3 for PacBio and Illumina reads from 50 known, 100 mixed, and 50 novel genomes. “Alignment” refers to the technique used to add reads into the marker gene alignment and “placement” refers to the runtime for phylogenetic placement (both are relevant only to TIPP3 and TIPP3-fast). The time shown does not include the creation of the reference package for the relevant method. Results for TIPP3 using APPLES2 and App-SpaM can be found in Fig S20.

Memory usage of methods is also shown in Fig S20, and all methods tested were able to complete within the 256 GB limit. Peak memory usage for TIPP3 was generally higher than all other methods. TIPP3-fast required less memory on all datasets than TIPP3 and for PacBio reads required almost as little as mOTUsv3, the most memory-efficient method tested. For Kraken2 and Bracken, the high memory usage for all datasets was likely due to loading the database to memory but could be greatly alleviated with the `--memory-mapping` option as suggested in Kraken2 [8]. We did not use this option for this study as we optimized the comparison for high accuracy rather than performance.

## Discussion

Our experiment revealed several consistent trends. For example, abundance profiling error is higher for PacBio reads than for Illumina reads, and higher as well for novel genomes than for known genomes. Another consistently observed trend is that error goes down as the taxonomic level increases (e.g., error is lower at the family level than at the species level). These trends have been observed before [14] and are expected.

Our results generally showed that TIPP3 had superior accuracy compared to the other methods, except for the easiest condition (Illumina reads from known genomes) where other methods were more accurate at the lower taxonomic levels. However, on the more challenging datasets, where there was either sequencing error or the reads were partially or fully from novel genomes, TIPP3 had an advantage. We now explore the evidence to see what contributes to TIPP3’s advantage over the other methods, including over TIPP2, for these more challenging conditions.

We begin with the comparison between TIPP3 and TIPP2.5, which allows us to assess the impact of using larger taxonomic trees, without changing other aspects of TIPP3. Given the substantial improvement in accuracy seen in TIPP3, this shows the benefit of using more densely sampled taxonomic trees for abundance profiling. This observation is consistent with the improvement of TIPP2 relative to TIPP [14]. Furthermore, this is also consistent with prior studies that have shown that more accurate phylogenetic placement can be obtained through the use of larger more densely sampled reference trees [20, 21].

For the comparison between TIPP3, Kraken2 (filtered or not), Bracken (filtered or not), and mOTUsv3, note that only TIPP3 is based on phylogenetic placement of reads added to a multiple sequence alignment, rather than analyses based on k-mers or BWA [40] alignment to reference sequences. Furthermore, TIPP3 can be paired with any phylogenetic placement method, and our study explored the use of APPLES-2 [19], App-SpaM [33], and two maximum likelihood-based placement methods (pplacer-taxtastic and BSCAMPP). On the PacBio model conditions, using either APPLES2 or App-SpaM had substantially higher error than using pplacer-taxtastic or BSCAMPP (Fig S15), showing the importance of using maximum likelihood phylogenetic placement. However, pplacer-taxtastic provides somewhat more accurate results than using BSCAMPP (Fig S21); while there are several differences between the two placement methods, the divide-and-conquer strategy in BSCAMPP is known to reduce accuracy slightly, but provides a speed advantage, and this is likely the main contribution to the accuracy difference. Finally, how the reads are added to the marker gene alignments also impacts accuracy (Fig S21), with a small advantage to using WITCH compared to BLAST, emphasizing the value of multiple sequence alignment rather than pairwise alignment.

A comparison between TIPP3 and TIPP3-fast also provides insight into the impact of the choice of phylogenetic placement method (pplacer-taxtastic for TIPP3 and BSCAMPP for TIPP3-fast) as well as how reads are added into the multiple sequence alignment (WITCH for TIPP3 and BLAST for TIPP3-fast), as these are the only two differences between the methods. As shown in the supplementary materials Fig S21, both choices contribute to the improvement in accuracy for TIPP3 over TIPP3-fast, but the choice of phylogenetic placement method has a larger impact. The design of TIPP3-fast included these two changes even though they slightly reduced accuracy, because of the runtime advantage they gave, which together resulted in reducing the runtime substantially (Fig S22 and Table S2). Thus TIPP3-fast is competitive in runtime and still more accurate than Kraken2 (filtered or unfiltered), Bracken (filtered or unfiltered), and mOTUsv3, under challenging conditions.

Our study also explored the impact of reducing the number of marker genes in order to reduce the runtime and possibly improve accuracy, and it is worth noting that although we started with 40 marker genes, we selected only 38 of them for use in TIPP because this change improved accuracy. Our exploration of having a further reduction in the number of genes showed a reduction in accuracy, without a substantial improvement in runtime, and was discarded (Figs S14, S21, and S22). Overall, our study demonstrates the importance of choosing marker genes carefully.

Finally, our study showed the generally beneficial impact of modifying existing abundance profiling methods by restricting them to reads from marker genes, so that the “filtered” versions of Bracken and Kraken2 were generally more accurate than their original versions. We hypothesize that the main reason for the improvement is the ability to use only reads that come from single-copy universal genes (i.e., marker genes), and this hypothesis is consistent with our experiment on how error in Kraken2 and Bracken increases with genome size. Interestingly, we also saw that filtering reduced accuracy for Bracken when analyzing PacBio reads from novel genomes at higher taxonomic levels (order, class, and phylum). At this time, we do not have a hypothesis for why this is happening, but we do note that in general, this is the hardest model condition (PacBio reads from novel genomes) and that the number of reads that were generated was low compared to the number of reads generated for Illumina sequencing).

## Conclusions

In this study, we introduced a new method, TIPP3, for accurate abundance profiling. TIPP3 outperforms its predecessor TIPP2 in terms of profiling accuracy and also provides more accurate profiles than other taxonomic profiling tools—particularly when input reads have sequencing errors and come from genomes absent from reference databases used by these tools. TIPP3-fast is a much faster version of TIPP3, having comparable runtime to Kraken2, Bracken, and mOTUs and with only a small decrease in accuracy to TIPP3. Therefore, TIPP3-fast maintains TIPP3’s accuracy advantage over the other methods under hard conditions for abundance profiling. Given that microbial communities are abundant but mostly still under-explored and may include many currently unknown genomes, tools like TIPP3 and TIPP3-fast are valuable for the accurate characterization of these microbial communities.

Based on this study, we can make some recommendations for the choice of abundance profiling method. When working with Illumina reads from known genomes, then Bracken(filtered) is the most accurate method (and much more accurate than Bracken(all)). However, for other conditions, then TIPP3 is the most accurate, followed by TIPP3-fast. The choice between TIPP3 and TIPP3-fast essentially depends on how important runtime is compared to accuracy, as TIPP3 is much slower (50 to 150 times slower) than TIPP3-fast.

This study suggests several directions for further improvement of TIPP3. While TIPP3 achieves high profiling accuracy using the most accurate setting, it has a significantly slower runtime compared to other methods. The step in TIPP3 where reads are added into the marker gene alignment using WITCH is the biggest contribution to runtime, which suggests that developing new methods for this step that are substantially faster but not much less accurate than WITCH is a promising direction.

Another direction for future research is algorithm design to enable accuracy to continue to improve as the number of sequences in each marker gene increases. Although most of the algorithmic steps in TIPP3 are already known to work well on very large datasets (e.g., MAGUS for the marker gene alignment and WITCH for adding reads to marker gene alignments), pplacer-taxtastic is possibly restricted to about 100,000 sequences. If so, then either we would need to improve the scalability of pplacer-taxtastic, or rely on BSCAMPP and possibly other fast methods for phylogenetic placement.

## Supporting information

### Supporting materials

All supporting text, 22 supporting figures, and 2 supporting tables are included in the supplementary document.

## Supplementary Materials

### Supporting Materials

This document provides the supporting materials for TIPP3, including additional information for obtaining the reference package, software commands for benchmarking, information for input read generation and normalized Hellinger distance, and additional results.

#### S1 Additional Information on TIPP3 Reference Packages

We used the same NCBI RefSeq Bacteria and Archaea genomes as TIPP2 [1], downloaded in November 2019 from the NCBI RefSeq database. The TIPP3 reference package was then constructed using the following pipeline. We first extracted and cleaned the same set of 40 marker genes as TIPP2 from the RefSeq genomes. After filtering, we aligned each set of marker gene sequences with MAGUS [2] and built a taxonomy tree with RAxML [3]. For running pplacer [4] with the taxtastic package [5], we re-estimated tree branch lengths with FastTree-2 [6]. The latest TIPP3 reference package is available at https://doi.org/10.13012/B2IDB-4931852_V1. Scripts for data processing can be found at https://github.com/shahnidhi/TIPP_reference_package.

##### Pre-alignment data processing

1. Filter the set of Archaea and Bacteria genomes (a total of 173,240 genomes) for each marker gene by (1) removing sequences that do not have matched amino/nucleotide sequences, (2) removing sequences that are 3 standard deviations from the median length. (with `data/filterData.py`).
2. Update the taxids using the **NCBI taxonomy downloaded on July 5th, 2023** (with `taxit update_taxids`).
3. Create a new taxonomy table for each marker gene (with `taxit taxtable`).
4. Update species mapping (species name to taxid) (with `update_species_mapping.py`).
5. Build taxonomy for each marker gene (with `build_unrefined_tree.pl`).
6. The outputs, for each marker gene, are (1) a final filtered set of sequences (∼ 55, 000 sequences), and (2) an unrefined taxonomy of the sequences.

##### Alignment

1. Perform a non-recursive MAGUS alignment [2] on the filtered sequences for each marker gene.
2. The exact command of non-recursive MAGUS:

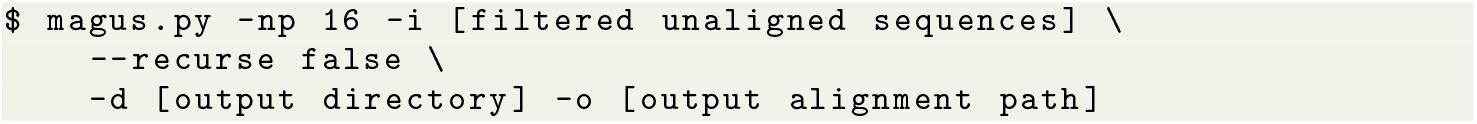
3. The output is an alignment for each set of marker gene sequences.

##### Build taxonomy

1. Back-translate the amino acid alignment to nucleotide alignment (with `backtranslate_refseq.py`).
2. Clean up the tree internal labels on the taxonomy before refining it (with `nw_topology -bI`).
3. Mask gappy sites (*>* 95% gaps) of the nucleotide alignment (with `ogcat mask -p 0.95`).
4. Resolve polytomies with RAxML [3] under the GTR+CAT model.

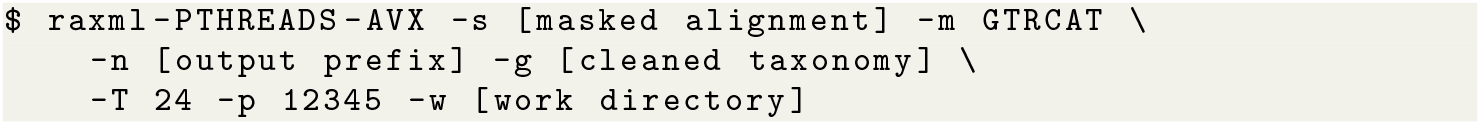
5. Add branch length information to the RAxML refined tree.

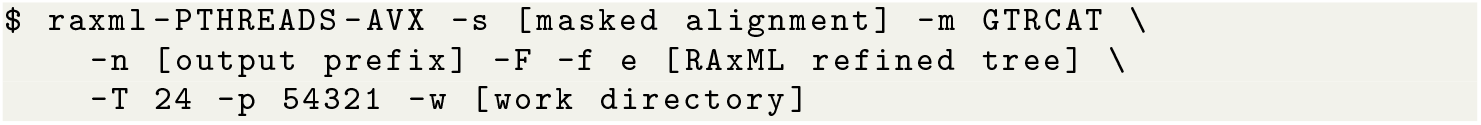
6. Re-estimate the branch lengths with RAxML-ng [7] under the GTR+GAMMA model.

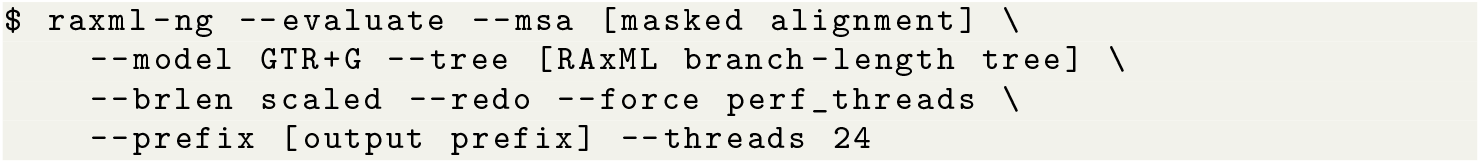
7. Add back the internal labels with taxonomic information (with `relabel-modified.py`).

##### Re-estimating numeric parameters

For `pplacer` [4] running with the `taxtastic` Python package [5], we re-estimated the taxonomy’s numeric parameters with FastTree-2 [6] using the following command:

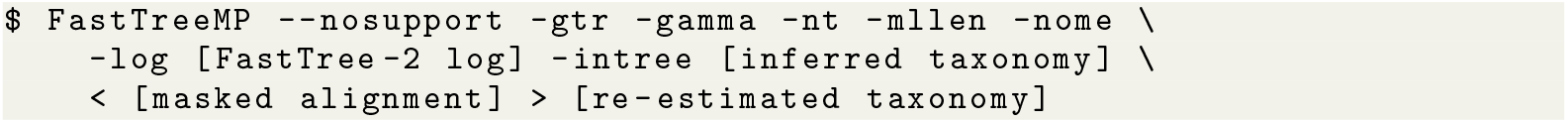

Then, a `taxtastic` package is created as:

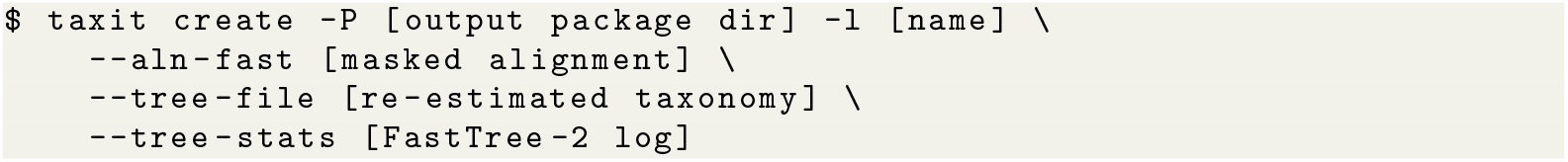

### S2 Software Commands

We ran all software with 16 CPU cores and up to 256 GB of memory, running each software with no time limit until completion.

1. We ran Kraken2 (v2.1.3) with the following command:

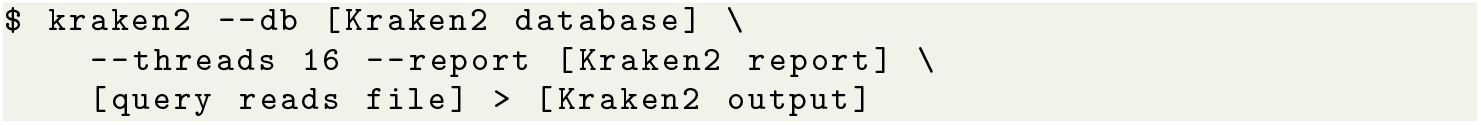
2. We ran Bracken (v2.9) with the following command (using Kraken2 output):

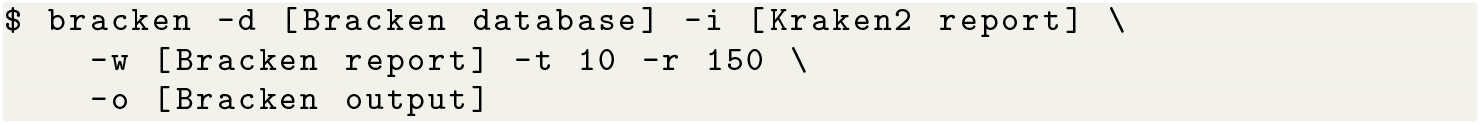
3. We ran mOTUsv3 (v3.1.0) with the following command:

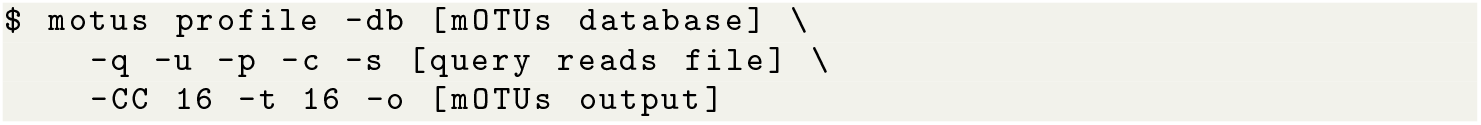
4. We ran APPLES-2 (v2.0.11) with the following command (for each marker gene and its assigned reads):

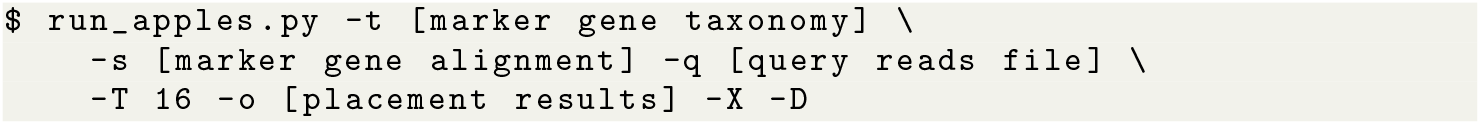
5. We ran App-SpaM (v1.03) with the following command (for each marker gene and its assigned reads):

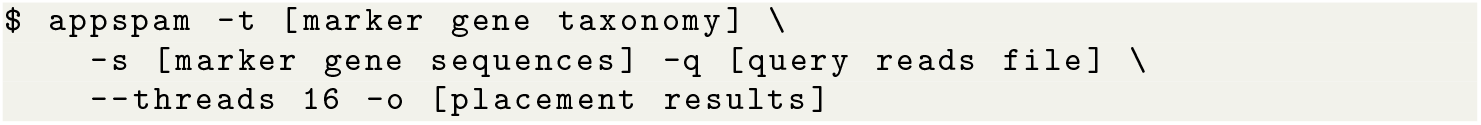
6. We ran SCAMPP (v2.0.1) with the following command (for each marker gene and its assigned reads):

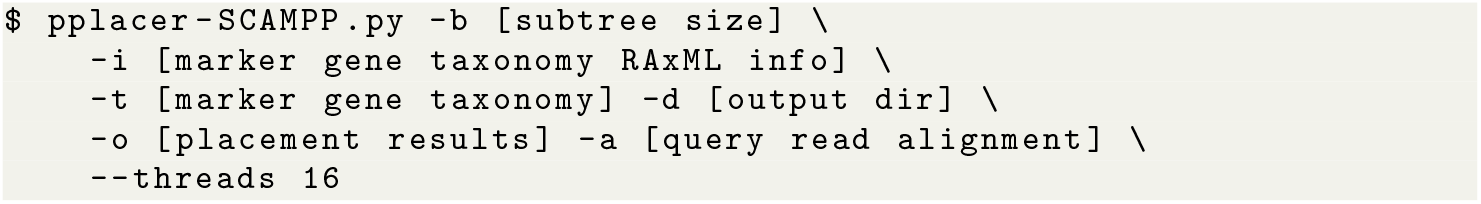
7. We ran BSCAMPP (v1.0.0) with the following command (for each marker gene and its assigned reads):

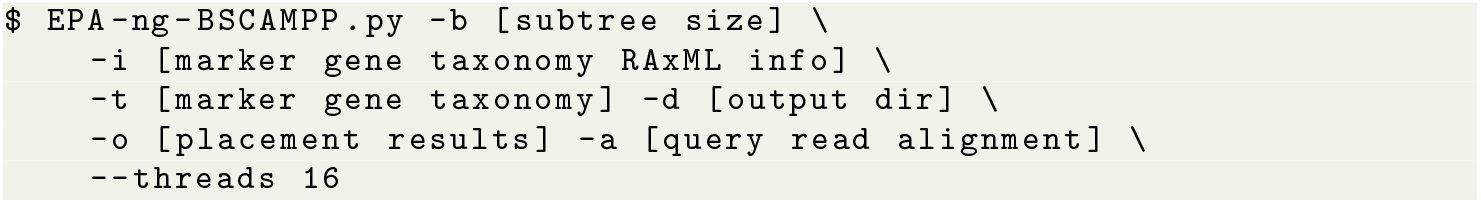
8. We ran pplacer (v1.1.alpha19-0-g807f6f3) with the `taxtastic` (v0.10.0) package with the following command (for each marker gene and its assigned reads):

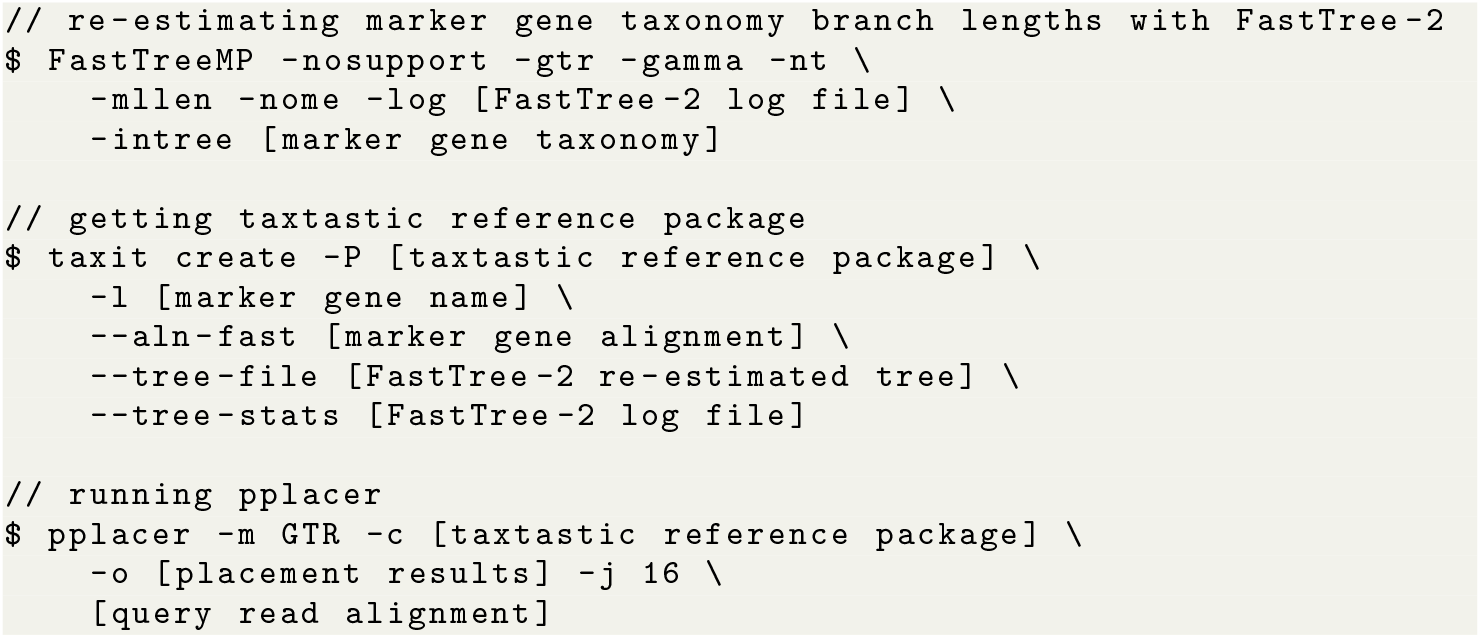

### S3 Additional Information for Read Generation

Reads are generated for each set of reference genomes using either the ART sequence simulator [8] or PBSIM [9]. For a given set of genome sequences, query reads are simulated in two ways: Illumina (150bp) or PacBio (∼ 3000bp). The commands used are shown below.

#### Illumina reads generation

Illumina reads are simulated with `art_illumina` v2.5.8 with the following command. The sequence length is 150 bp and the coverage is 20x.

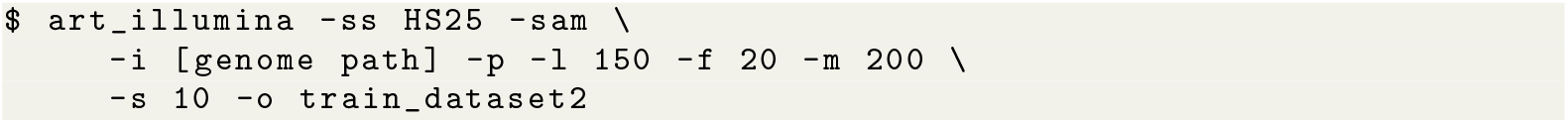

#### PacBio reads generation

PacBio reads are simulated with `pbsim` with the following command. The model used is CLR, and the average sequence length is set to 3000bp with a minimum length of 400bp. The average accuracy is set to 0.78 with a standard deviation of 0.07. According to the authors [9], the simulated data generated by `pbsim` with the CLR model and a 0.78 accuracy has 3.23% substitution rate, 10.53% insertion rate, and 3.98% deletion rate (obtained by aligning simulated reads to reference using LAST [10]). The coverage is also set to 20x.

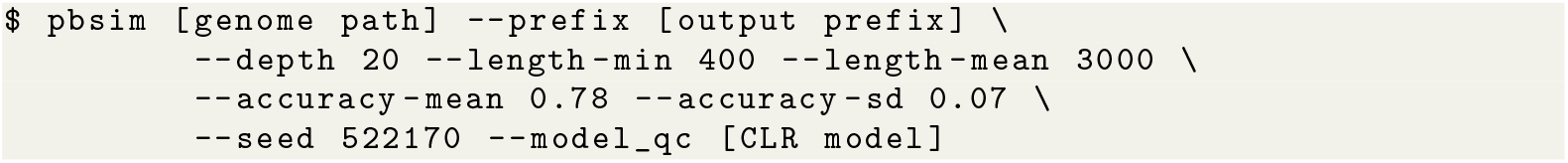

#### Genome accession numbers for reads generation

##### 1. TIPP2 dataset 1 (33 genomes)

GCA_002214465.1, GCA_002844335.1, GCA_002906575.1, GCA_002951815.1, GCA_003430825.1, GCA_006739055.1, GCA_006742785.1, GCA_011106835.1, GCA_011764545.1, GCA_012222825.1, GCA_013085545.1, GCA_013177635.1, GCA_013177655.1, GCA_013201685.1, GCA_013201895.1, GCA_013267355.1, GCA_013283835.1, GCA_013347265.1, GCA_013391845.1, GCA_013394065.1, GCA_013409125.2, GCA_013415885.1, GCA_013466785.1, GCA_013488225.1, GCA_014042035.1, GCA_014076455.1, GCA_014076495.1, GCA_014189535.1, GCA_014217485.1, GCA_014218275.1, GCA_900324035.1, GCA_900631605.1, GCA_902702935.1

##### 2. TIPP2 dataset 2 (51 genomes)

GCF_000020225.1, GCF_000265505.1, GCF_000238215.1, GCF_000183135.1, GCF_000190735.1, GCF_000190595.1, GCF_000020145.1, GCF_000069185.1, GCF_000092825.1, GCF_000023245.1, GCF_000023105.1, GCF_000063485.1, GCF_000020465.1, GCF_000218625.1, GCF_000484535.1, GCF_000020965.1, GCF_000021565.1, GCF_000183745.1, GCF_000191445.1, GCF_000192885.1, GCF_000018385.1, GCF_000018145.1, GCF_000009365.1, GCF_000020945.1, GCF_000022565.1, GCF_000266885.1, GCF_000195435.3, GCF_000017025.1, GCF_000163895.2, GCF_000147095.1, GCF_000024205.1, GCF_000025025.1, GCF_000018865.1, GCF_000017805.1, GCF_000021685.1, GCF_000024985.1, GCF_000092105.1, GCF_000024565.1, GCF_000025885.1, GCF_000279145.1, GCF_000010305.1, GCF_000441755.1, GCF_000253035.1, GCF_000298475.2, GCF_000237205.1, GCF_000092785.1, GCF_000286715.1, GCF_000018785.1, GCF_000021265.1, GCF_000284335.1, GCF_000235405.2

##### 3. Testing, 50 known genomes

GCF_000020225.1, GCF_000265505.1, GCF_000238215.1, GCF_000183135.1, GCF_000190735.1, GCF_000190595.1, GCF_000020145.1, GCF_000092825.1, GCF_000023245.1, GCF_000023105.1, GCF_000063485.1, GCF_000020465.1, GCF_000218625.1, GCF_000484535.1, GCF_000020965.1, GCF_000021565.1, GCF_000183745.1, GCF_000191445.1, GCF_000192885.1, GCF_000018385.1, GCF_000018145.1, GCF_000009365.1, GCF_000020945.1, GCF_000022565.1, GCF_000266885.1, GCF_000195435.3, GCF_000017025.1, GCF_000163895.2, GCF_000147095.1, GCF_000024205.1, GCF_000025025.1, GCF_000018865.1, GCF_000017805.1, GCF_000021685.1, GCF_000024985.1, GCF_000092105.1, GCF_000024565.1, GCF_000025885.1, GCF_000279145.1, GCF_000010305.1, GCF_000441755.1, GCF_000253035.1, GCF_000298475.2, GCF_000237205.1, GCF_000092785.1, GCF_000286715.1, GCF_000018785.1, GCF_000021265.1, GCF_000284335.1, GCF_000235405.2

##### 4. Testing, 100 mixed genomes

GCF_030758975.1, GCF_964019505.1, GCF_032268605.1, GCF_030490285.1, GCF_029625215.1, GCF_036670025.1, GCF_037948395.1, GCF_030285785.1, GCF_030408375.1, GCF_032594115.1, GCF_034554815.1, GCF_035930445.1, GCF_030296615.1, GCF_033344015.1, GCF_964020205.1, GCF_964020185.1, GCF_030719275.1, GCF_031885465.1, GCF_036352135.1, GCF_030296595.1, GCF_024346835.1, GCF_030419005.1, GCF_964019495.1, GCF_964019535.1, GCF_036204065.1, GCF_033126985.1, GCF_030389495.1, GCF_963556095.2, GCF_030540775.2, GCF_037076455.1, GCF_036250655.1, GCF_030687935.1, GCF_033095965.1, GCF_025232395.1, GCF_025736875.1, GCF_030296575.1, GCF_030409055.1, GCF_947241125.1, GCF_030177935.1, GCF_030296055.1, GCA_013409125.2, GCA_013200995.1, GCA_004295565.1, GCA_013201685.1, GCA_014217485.1, GCA_900000005.1, GCA_013177635.1, GCA_002813655.1, GCA_014236795.1, GCA_012225885.1, GCA_011455695.1, GCA_013391845.1, GCA_014042035.1, GCA_014069315.1, GCA_900631605.1, GCA_002156705.1, GCA_014048085.1, GCA_011462075.1, GCA_004799685.1, GCA_008326385.1, GCA_000956175.1, GCA_002214465.1, GCA_002214385.1, GCA_002214485.1, GCA_013347265.1, GCA_003201835.2, GCA_013170725.1, GCA_009884315.1, GCA_013170785.1, GCA_013154935.1, GCA_011106835.1, GCA_014076455.1, GCA_013283835.1, GCA_013201895.1, GCA_013201665.1, GCA_013201935.1, GCA_009789175.1, GCA_004768745.1, GCA_013693755.1, GCA_013201825.1, GCA_008931705.1, GCA_002499975.2, GCA_001971705.1, GCA_002214505.1, GCA_002355655.1, GCA_002844335.1, GCA_013201725.1, GCA_000875775.1, GCA_003430825.1, GCA_013347305.1, GCA_002906575.1, GCA_009729015.1, GCA_002214565.1, GCA_006739055.1, GCA_012923785.1, GCA_003290265.1, GCA_012427845.1, GCA_013085545.1, GCA_003935895.2, GCA_900324035.1

##### 5. Testing, 50 novel genomes

GCF_017357865.1, GCF_034262375.1, GCF_019880205.1, GCF_029101545.1, GCF_034258515.1, GCF_030316605.1, GCF_030252185.1, GCF_036492835.1, GCF_034479515.1, GCF_007833795.1, GCF_964019775.1, GCF_027563145.1, GCF_030438475.1, GCF_030407165.1, GCF_029023725.1, GCF_964019755.1, GCF_030294945.1, GCF_943169825.2, GCF_030408715.1, GCF_963675105.1, GCF_034479615.1, GCF_034371605.1, GCF_030295325.1, GCF_964019545.1, GCF_029101585.1, GCF_933509905.1, GCF_964019365.1, GCF_032397745.1, GCF_020865585.1, GCF_020524035.2, GCF_030622045.1, GCF_036320655.1, GCF_030296555.1, GCF_030408455.1, GCF_027923555.1, GCF_034359465.1, GCF_030294405.1, GCF_033472595.1, GCF_031312535.1, GCF_008726475.3, GCF_030406165.1, GCF_036178885.1, GCF_018128945.1, GCF_030845235.1, GCF_005517195.1, GCF_030758955.1, GCF_030550815.1, GCF_964020195.1, GCF_030408755.1, GCF_026250525.1

### S4 Additional Information for Normalized Hellinger Distance

#### S4.1 Normalized Hellinger distance

Remember that regular Hellinger distance is given by:

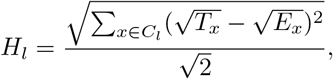

##### Theorem S4.1

(Theoretically Worst Hellinger Distance). *If an abundance profiling method classifies n reads in total, and n_l_ reads are assigned with taxonomic labels at the taxonomic level l, then the theoretically worst Hellinger distance that the method can have at taxonomic level l is* 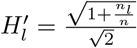

*Proof*. In the worst case, an estimated profile is disjoint from the true profile at taxonomic level *l*. More formally this means that for any clade *x* ∈ *C*_*l*_, where *C*_*l*_ is the union of labels in the estimated and true profiles, *T*_*x*_ = 0 or *E*_*x*_ = 0, exclusively. This also means 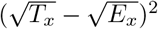 is either *T*_*x*_ or *E*_*x*_, depending on if clade *x* is defined in the estimated or the true profile.

While the true profile sums to 1 by definition (i.e., taxonomic labels are known for reads), only 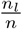 of the estimated profile counts toward the Hellinger distance computation at level *l*. Then, we can rewrite the Hellinger distance computation as:

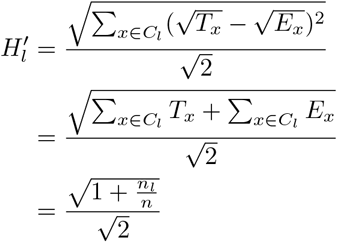

By Theorem S4.1, the Hellinger distance *H*_*l*_ of an estimated profile is bounded by 0 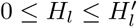 at a taxonomic level *l*, and 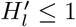. Only when all classified reads are assigned taxonomic labels at level *l* (*n*_*l*_ = *n*), the range is from 0 to 1. Therefore, when *H*_*l*_ is used to measure a method for abundance profiling accuracy, it would be biased if the method fails to classify all reads at level *l*.

A simple alternative is to measure profiling accuracy by 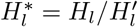, normalizing the measurement range to [0, 1] and being independent of the number of reads classified at a taxonomic level for each method (*n*_*l*_). We refer to this measurement as **normalized Hellinger distance**.

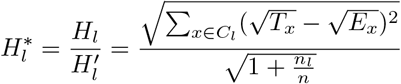

#### S4.2 Example for biases with Hellinger distance

Regular Hellinger distance is biased when the profile has a high fraction of unclassified reads. For example, let *n* be the total number of reads classified. At the species level, let a method *A* in total estimate 5 types of species [*s*_1_, *s*_2_, *s*_3_, *s*_4_, *s*_5_], where *s*_*i*_ = 0.1, *i* = 1, 2, 3, 4, 5. Then, the proportion of classified reads at taxonomic level *l* for method *A* is *n*_*l,A*_ = 0.5 and *n*_*l,A*_*/n* = 0.5, with the other 0.5 as “unclassified/unspecified”. In calculating Hellinger distance *H*_*l,A*_ between the profile of method *A* to the reference profile at level *l*, only the 0.5 that is classified/specified is used. Hence, the theoretically worst Hellinger distance 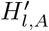 that method *A* can have is to have a disjoint profile to the reference, meaning that 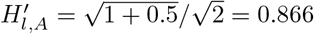. In other words, given a method’s estimated profile, 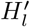 is the actual upper bound for *H*_*l*_.

In the most extreme case when a method *B* fails to produce *any classified groups* and has 1.0 as unclassified, then 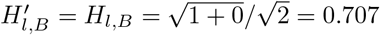. Let’s assume that the reference profile looks like [*s*_100_ = 0.7, *s*_150_ = 0.3]. Another method *C* may make some meaningful estimation, such as [*s*_1_ = 0.1, *s*_2_ = 0.1, *s*_3_ = 0.1, *s*_4_ = 0.1, *s*_100_ = 0.05], as it gets only one group overlapping with the reference. If we compute the Hellinger distance for method *C*, we will get:

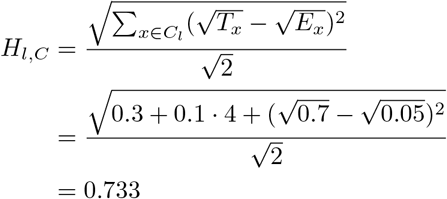

We can see that although method *B* does not give any estimation, it still gets a lower Hellinger distance than method *C*, which indicates a bias of Hellinger distance that favors making no estimation.

On the other hand, if we compare the normalized Hellinger distances of the two methods, we have:

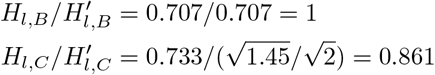

where we have method *C* having a lower error, which makes more sense.

### S5 Additional Results for Experiment 1: Designing TIPP3

#### S5.1 Adding reads to marker gene alignments

We obtained UPP and WITCH alignments of reads binned to marker genes RplO, RpsK, and RpsL. WITCH was run with different parameter settings, varying the number of hidden Markov models (HMMs) for aligning each read and the decomposition subset sizes (i.e., how many sequences to include in a subset to build an HMM on). Then, batch-SCAMPP (BSCAMPP) [11] was used to obtain placement and taxonomic identification of each read in each marker gene taxonomy.

Based on overall taxonomic identification accuracy, we selected WITCH with one hidden Markov model subset and subset size ranging from 10 to 1000 sequences as our query read alignment method. Full results for the alignment benchmarks can be found below.

**Fig S1.**
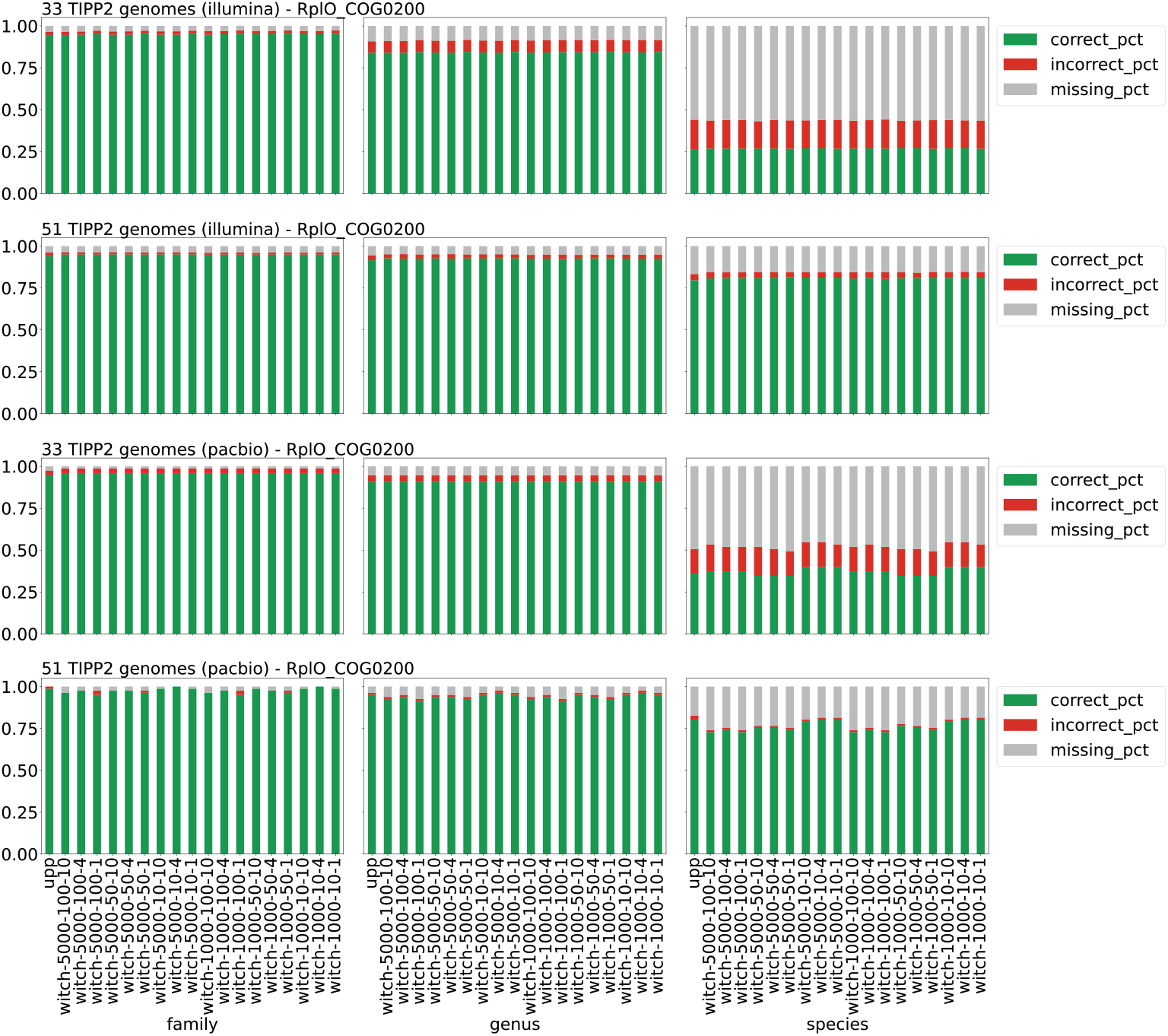
Taxonomic identification accuracy of UPP and different WITCH alignment variants on marker gene RplO_COG0200, for Illumina and PacBio reads from two TIPP2 datasets with 33 and 51 genomes (Family, Genus, and Species levels). The top two panels are for Illumina reads, and the bottom two are for PacBio reads. Taxonomic identification is done by performing query placements with Batch-SCAMPP with a subtree size of 1000 and a support value of 95%. WITCH variants are referred to as **witch-Z-A-k**, for which *Z* denotes the upper bound for subset size, *A* the lower bound, and *k* the number of subsets to align a query read. “correct_pct” denotes the fraction of correctly identified reads (at a taxonomic level), “incorrect_pct” denotes the fraction of incorrectly identified reads, and “missing_pct” denotes the fraction of not-identified reads. The three fractions sum to 1 for each bar.

**Fig S2.**
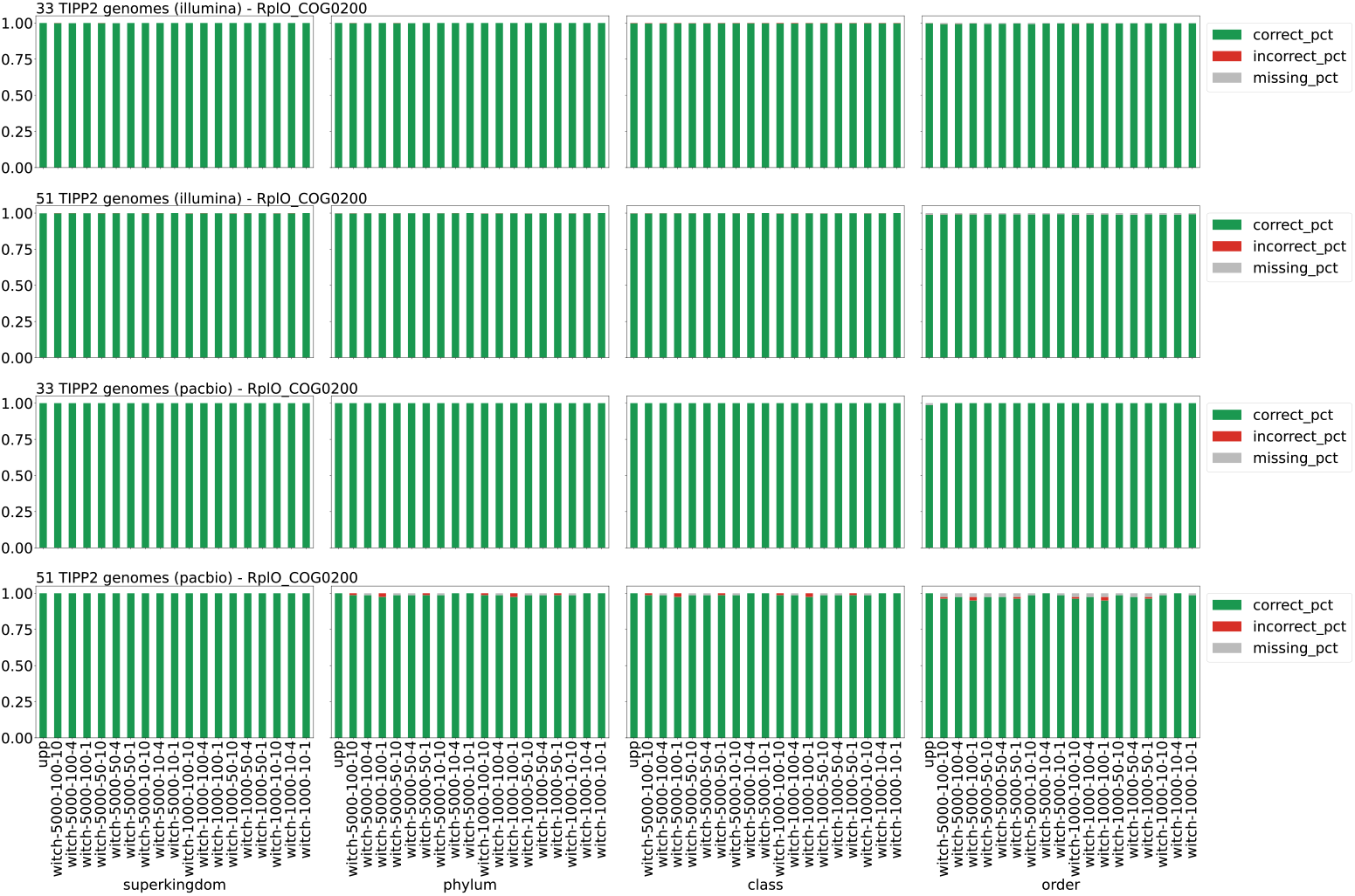
Taxonomic identification accuracy of UPP and different WITCH alignment variants on marker gene RplO_COG0200, for Illumina and PacBio reads from two TIPP2 datasets with 33 and 51 genomes (Superkingdom, Phylum, Class, and Order levels). The top two panels are for Illumina reads, and the bottom two are for PacBio reads. Taxonomic identification is done by performing query placements with Batch-SCAMPP with a subtree size of 1000 and a support value of 95%. WITCH variants are referred to as **witch-Z-A-k**, for which *Z* denotes the upper bound for subset size, *A* the lower bound, and *k* the number of subsets to align a query read. “correct_pct” denotes the fraction of correctly identified reads (at a taxonomic level), “incorrect_pct” denotes the fraction of incorrectly identified reads, and “missing_pct” denotes the fraction of not-identified reads. The three fractions sum to 1 for each bar.

**Fig S3.**
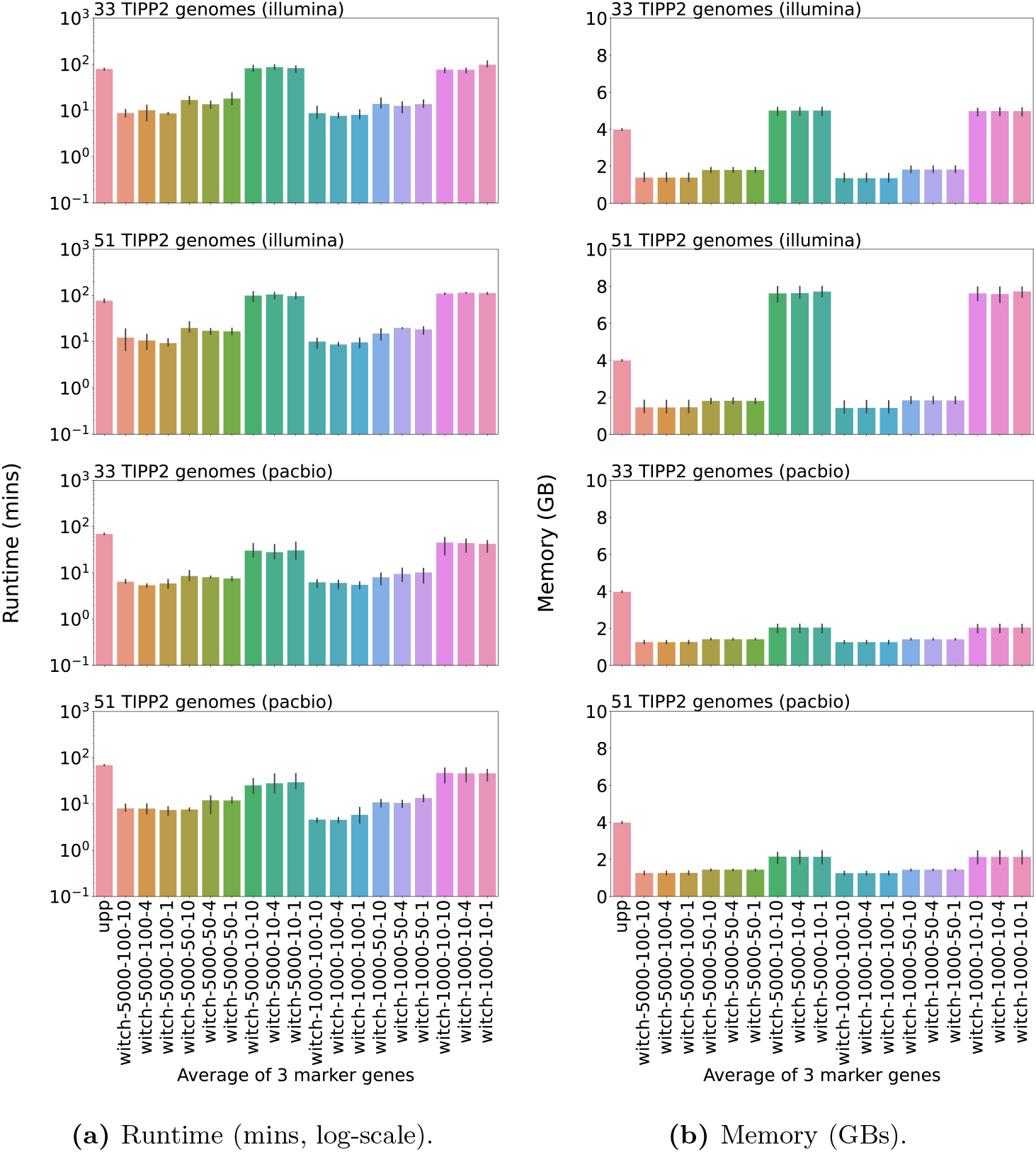
Runtime usage in minutes (left, log-scale) and memory usage in GBs (right) of identifying Illumina and PacBio reads from two TIPP3 datasets with 33 and 51 genomes, by UPP and different WITCH variants. Results are averaged over three marker genes (RplO, RpsK, and RpsL). Runtime for each variant includes read alignment and placement phases.

#### S5.2 Placing reads into marker gene taxonomies

After selecting the read alignment method, we explored different phylogenetic placement methods that place aligned reads into a target tree (i.e., the taxonomy of each marker gene), including SCAMPP [12], BSCAMPP [11], and pplacer with the `taxtastic` package (referred to as pplacer-taxtastic) [4, 13]. Both SCAMPP and BSCAMPP are divide-and-conquer methods that decompose the input tree into smaller subtrees and select suitable subtrees for query placements, using pplacer (SCAMPP) or EPA-ng [14] (BSCAMPP). We varied subtree sizes ranging from 1000 to 5000 leaves for SCAMPP and BSCAMPP, and support values ranging from 50% to 99% for all three methods (see main text for the definition of a support value). We then examined the taxonomic identification and abundance profiling accuracy of each method variant.

Overall, pplacer-taxtastic with a support value of 90% provides the best accuracy, while BSCAMPP with a subtree of size 1000 and a support value of 95% gives good accuracy and the fastest runtime. Hence, we selected pplacer with the `taxtastic` package (90% support value) as the read placement method for TIPP3. Full results for the placement benchmarks can be found below.

##### S5.2.1 Batch-SCAMPP

**Fig S4.**
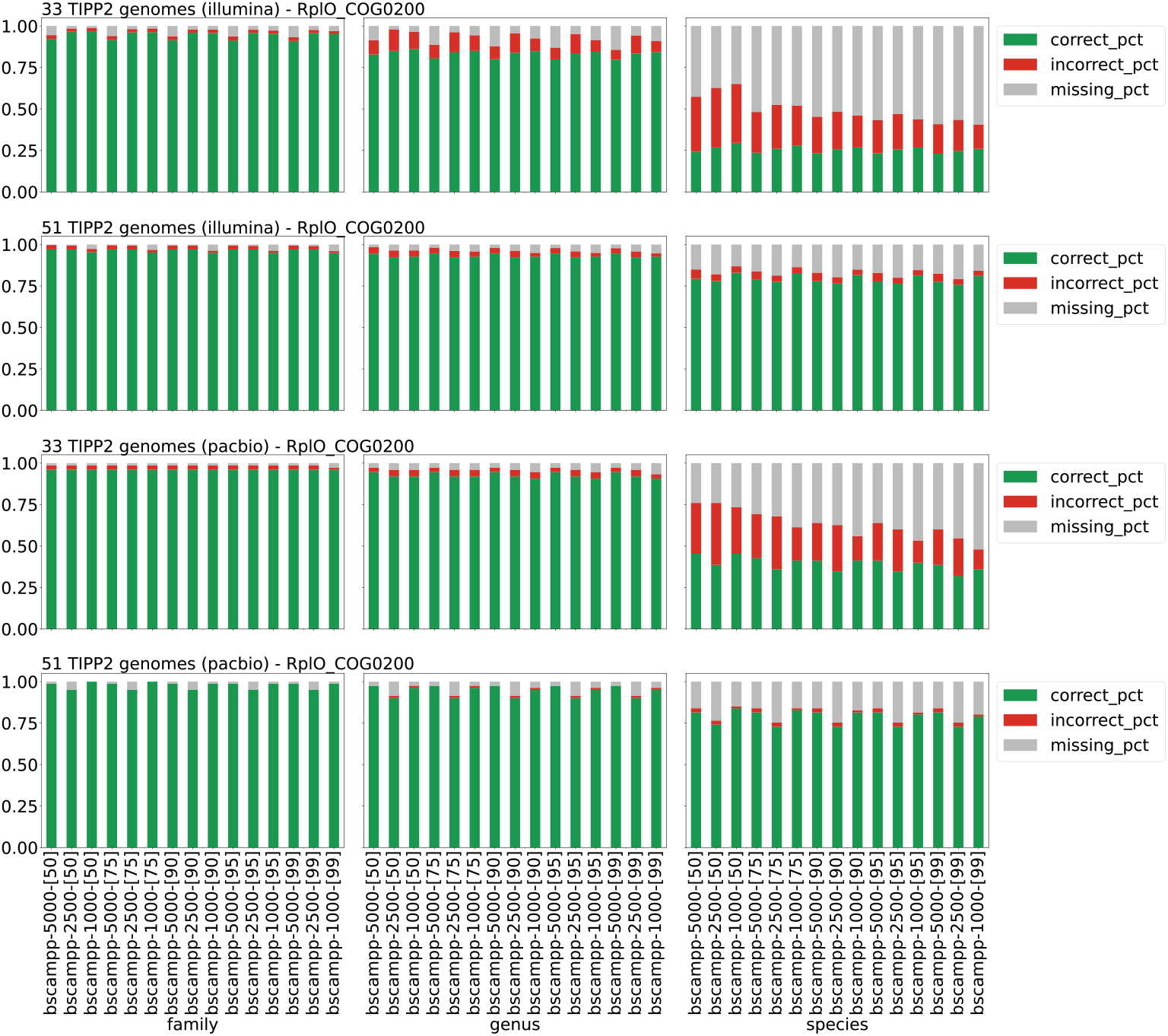
Taxonomic identification accuracy of different batch-SCAMPP variants for query placement in TIPP3 on marker gene RplO_COG0200, for Illumina and PacBio reads from two TIPP2 datasets with 33 and 51 genomes (Family, Genus, and Species levels). Query reads are aligned with WITCH. Batch-SCAMPP variants are named **bscampp-X-[Y]**, where *X* is the subtree size (*X* = {1000, 2500, 5000}) and *Y* is the support value (*Y* = {50, 75, 90, 95, 99}). “correct_pct” denotes the fraction of correctly identified reads (at a taxonomic level), “incorrect_pct” denotes the fraction of incorrectly identified reads, and “missing_pct” denotes the fraction of not-identified reads. The three fractions sum to 1 for each bar.

**Fig S5.**
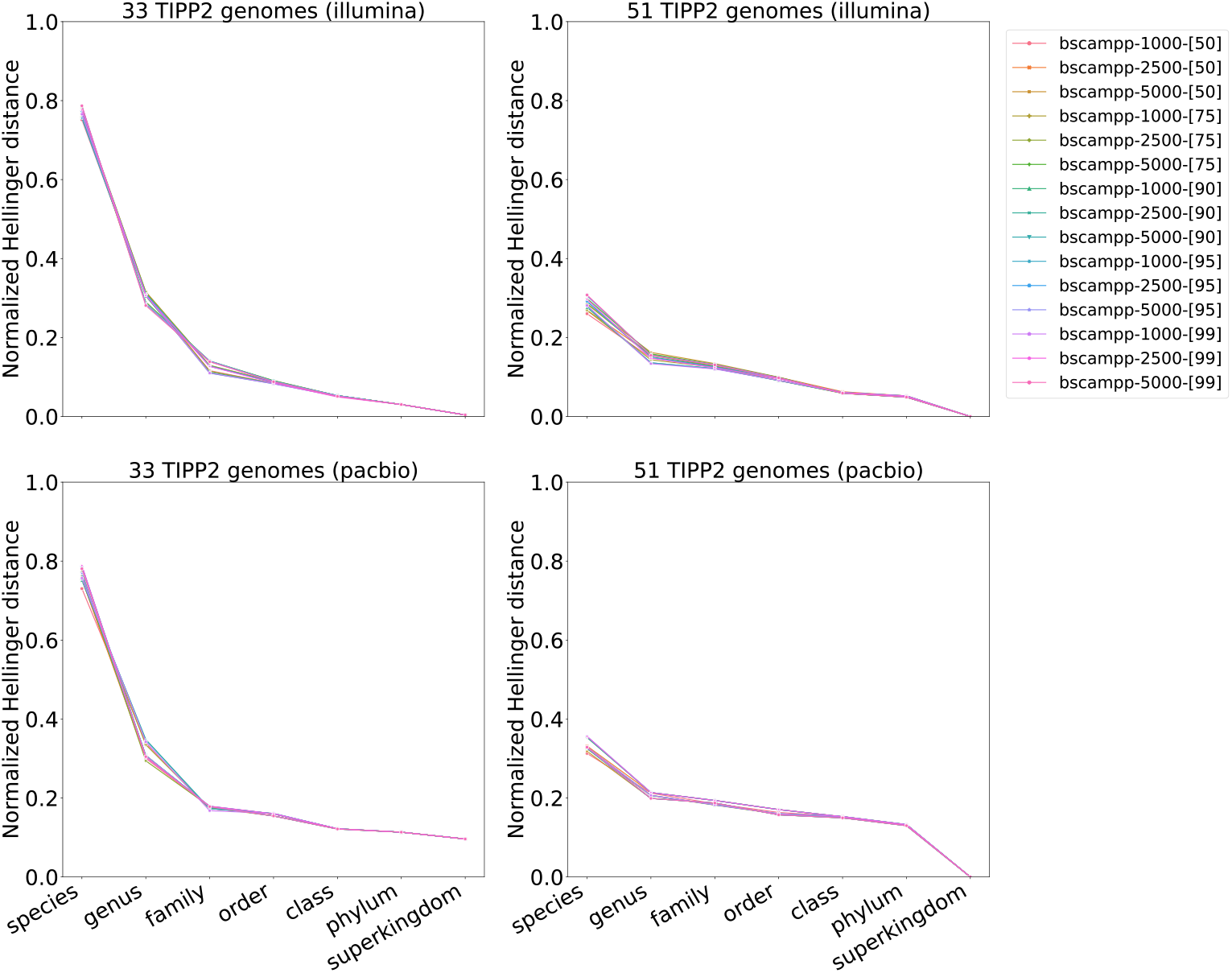
Abundance profile of different batch-SCAMPP variants for query placement in TIPP3, for Illumina and PacBio reads from two TIPP2 datasets with 33 and 51 genomes. Query reads are aligned with WITCH. The abundance profile is computed as the normalized Hellinger distance between the estimated and reference profiles using three marker genes, RplO, RpsK, and RpsL.

##### S5.2.2 SCAMPP

**Fig S6.**
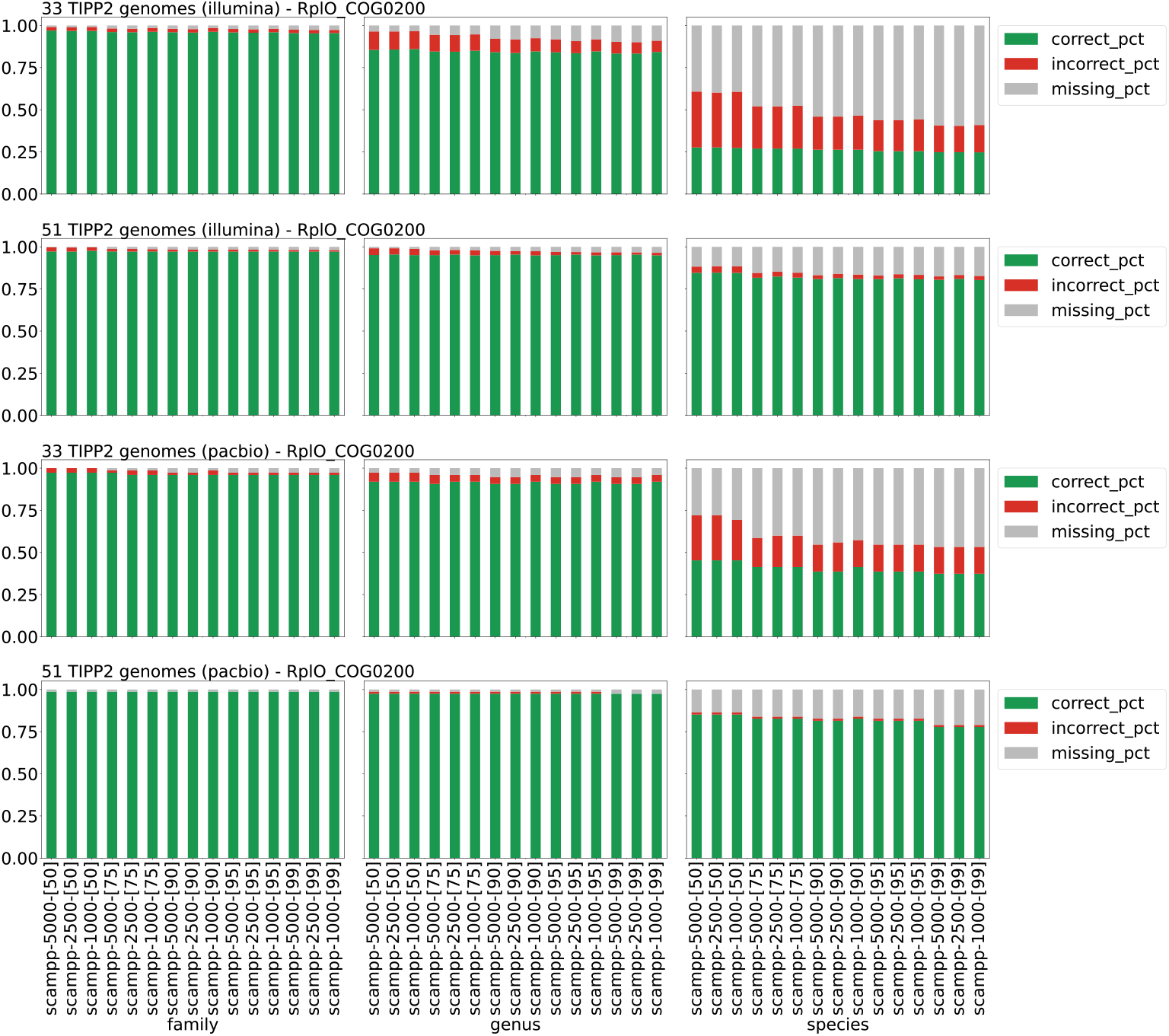
Taxonomic identification accuracy of different SCAMPP variants for query placement in TIPP3 on marker gene RplO_COG0200, for Illumina and PacBio reads from two TIPP2 datasets with 33 and 51 genomes (Family, Genus, and Species levels). Query reads are aligned with WITCH. SCAMPP variants are named **scampp-X-[Y]**, where *X* is the subtree size (*X* = {1000, 2500, 5000 }) and *Y* is the support value (*Y* = {50, 75, 90, 95, 99}). “correct_pct” denotes the fraction of correctly identified reads (at a taxonomic level), “incorrect_pct” denotes the fraction of incorrectly identified reads, and “missing_pct” denotes the fraction of not-identified reads. The three fractions sum to 1 for each bar.

**Fig S7.**
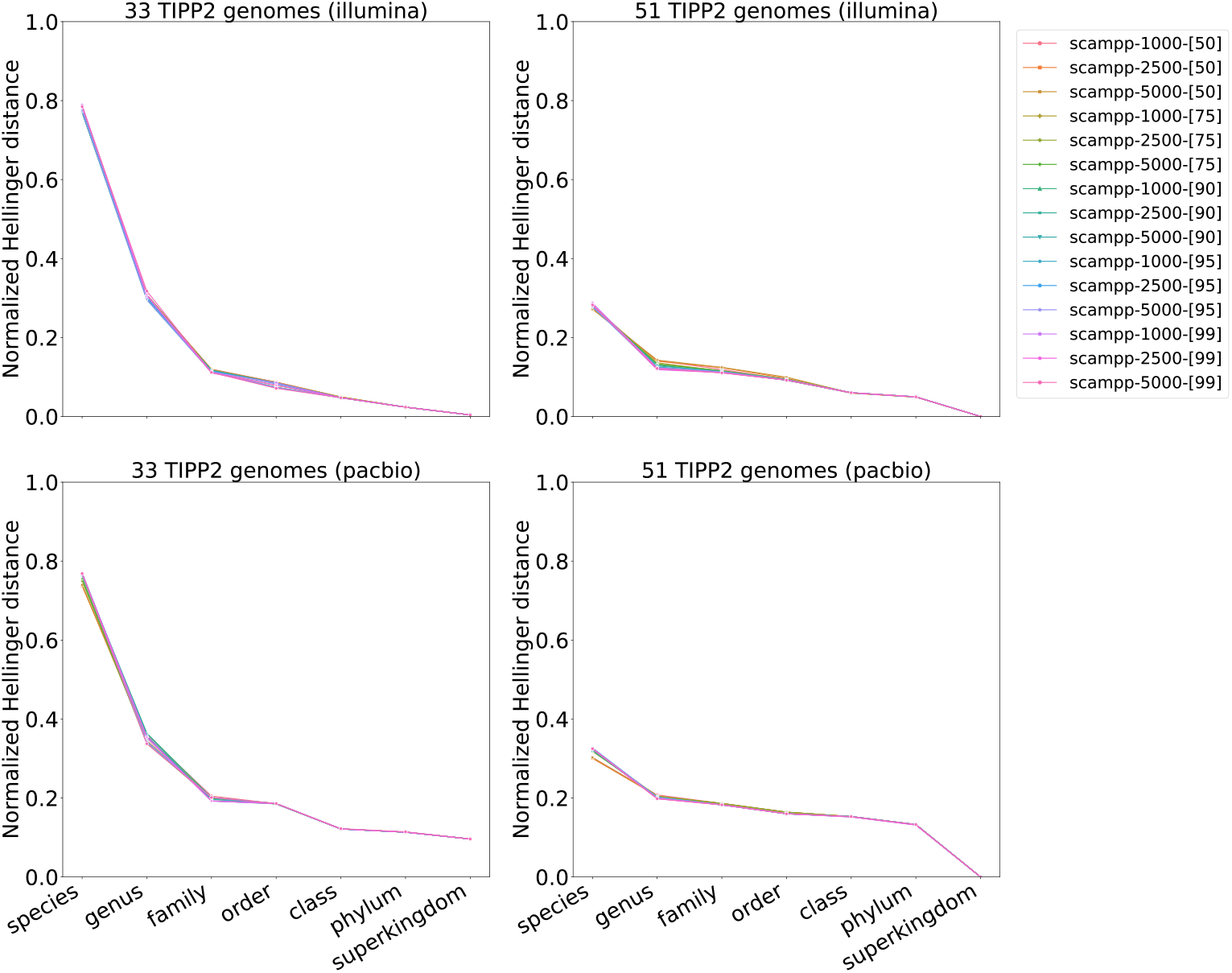
Abundance profile of different SCAMPP variants for query placement in TIPP3, for Illumina and PacBio reads from two TIPP2 datasets with 33 and 51 genomes. Query reads are aligned with WITCH. The abundance profile is computed as the normalized Hellinger distance between the estimated and reference profiles using three marker genes, RplO, RpsK, and RpsL.

##### S5.2.3 pplacer-taxtastic

**Fig S8.**
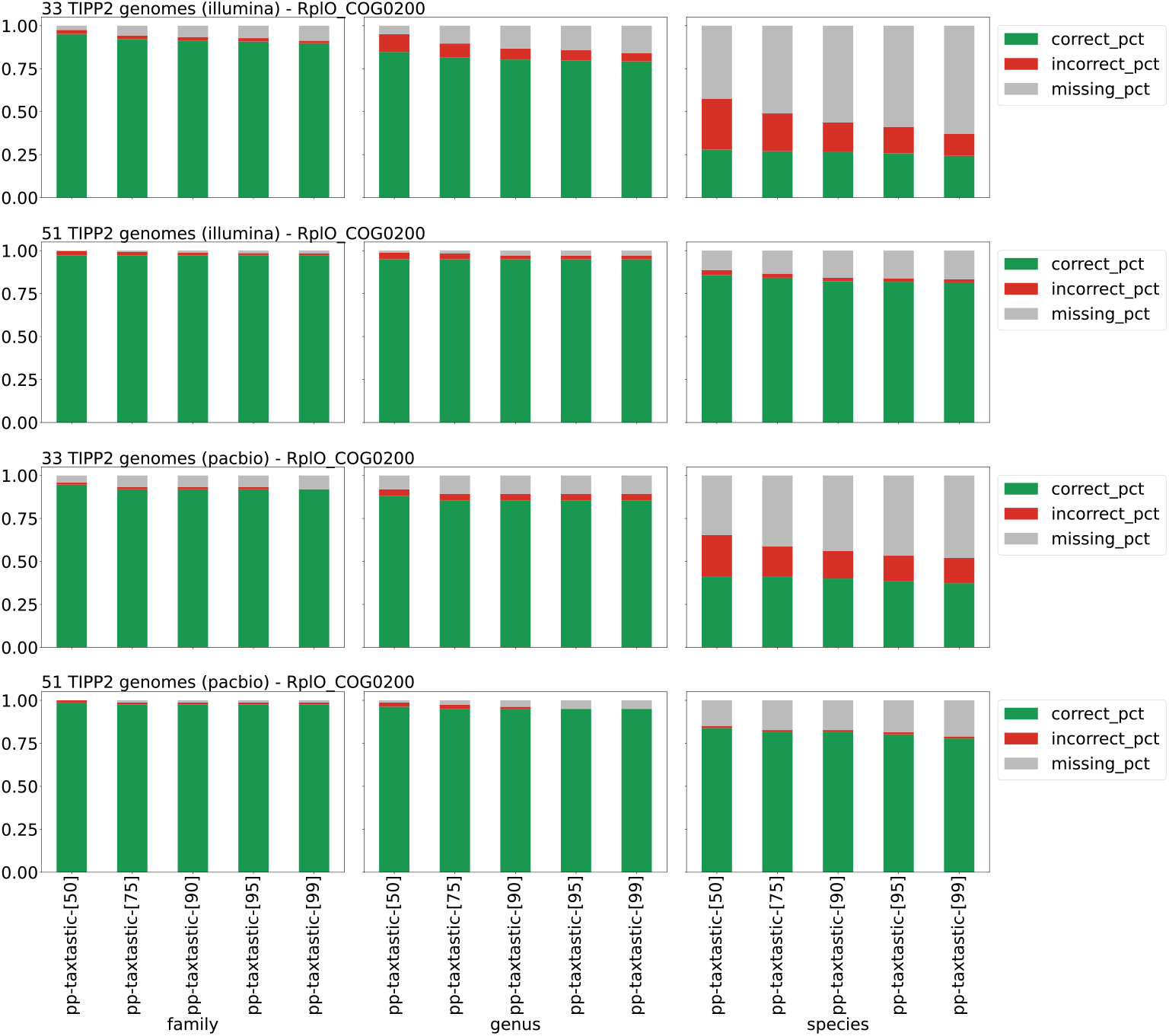
Taxonomic identification accuracy of different pplacer-taxtastic variants for query placement in TIPP3 on marker gene RplO_COG0200, for Illumina and PacBio reads from two TIPP2 datasets with 33 and 51 genomes (Family, Genus, and Species levels). Query reads are aligned with WITCH. pplacer-taxtastic variants are named **pp-taxtastic-[Y]**, where *Y* is the support value (*Y* = {50, 75, 90, 95, 99}). “correct_pct” denotes the fraction of correctly identified reads (at a taxonomic level), “incorrect_pct” denotes the fraction of incorrectly identified reads, and “missing_pct” denotes the fraction of not-identified reads. The three fractions sum to 1 for each bar.

**Fig S9.**
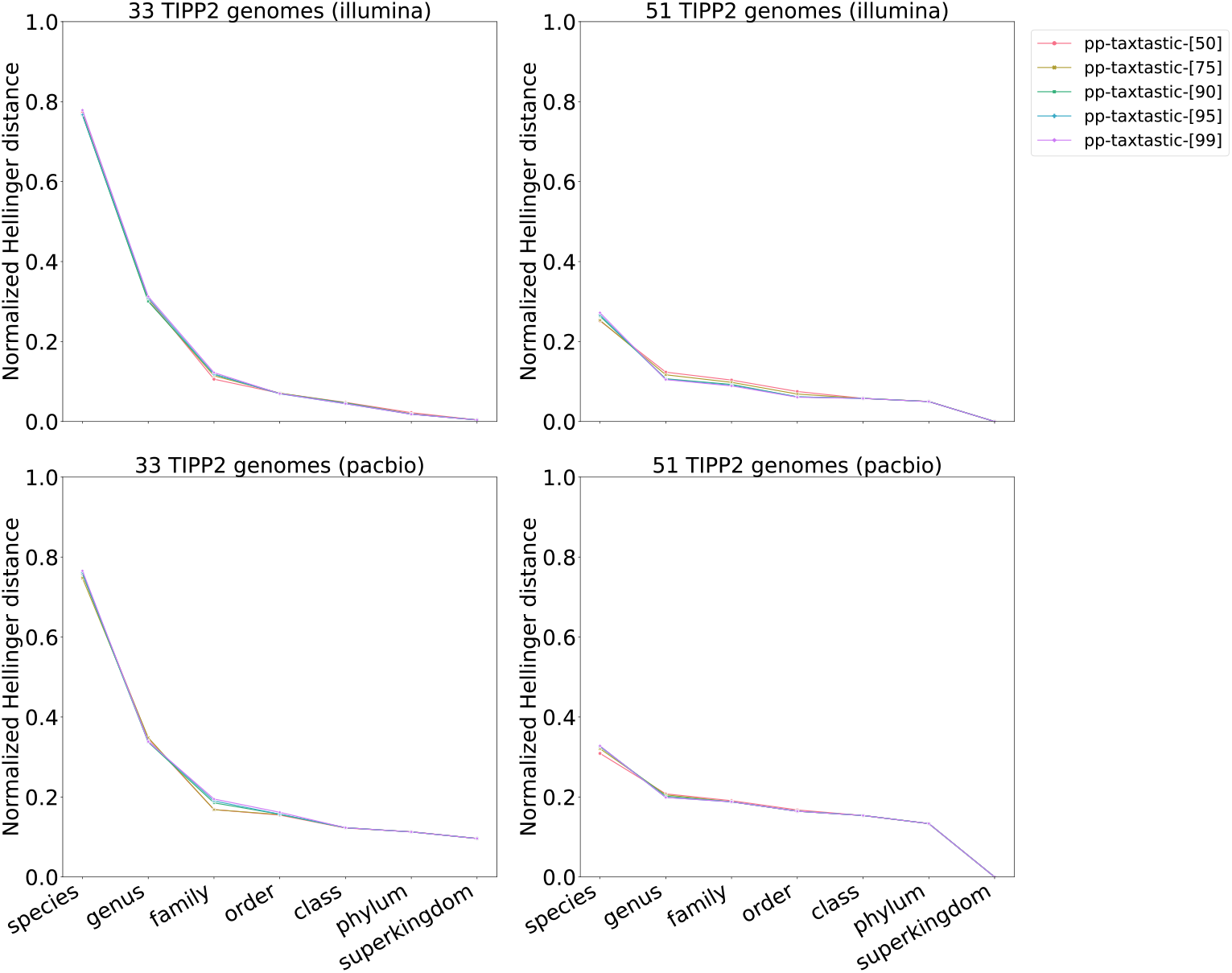
Abundance profile of different pplacer-taxtastic variants for query placement in TIPP3, for Illumina and PacBio reads from two TIPP2 datasets with 33 and 51 genomes. Query reads are aligned with WITCH. The abundance profile is computed as the normalized Hellinger distance between the estimated and reference profiles using three marker genes, RplO, RpsK, and RpsL.

##### S5.2.4 Comparison of the three placement methods

**Fig S10.**
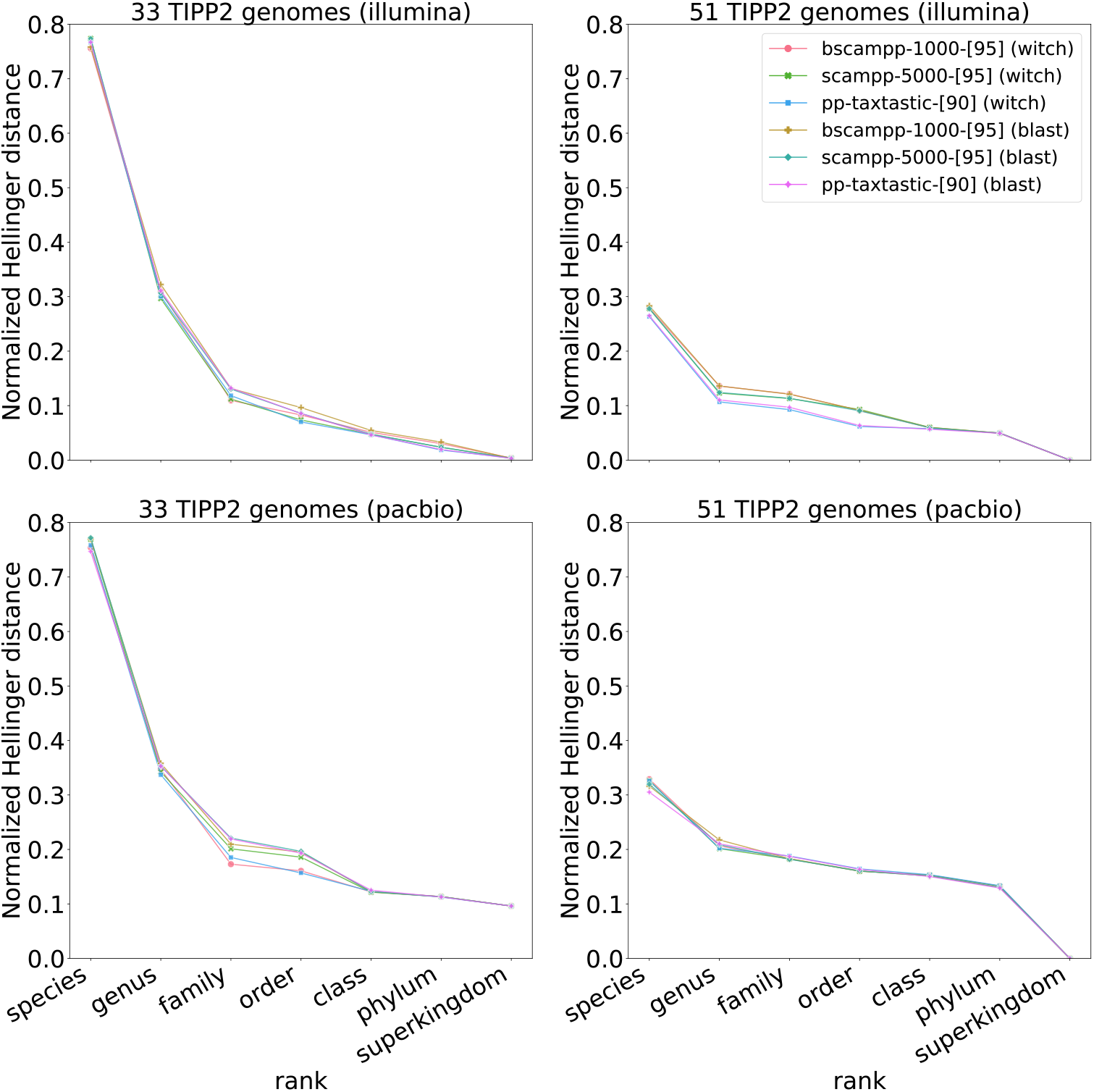
Abundance profile of the best variants of batch-SCAMPP, SCAMPP, and pplacer-taxtastic for Illumina and PacBio reads from two TIPP2 datasets with 33 and 51 genomes. Query reads are aligned with either WITCH or BLAST (denoted after the name of each method). The abundance profile is computed as the normalized Hellinger distance between the estimated and reference profiles using three marker genes, RplO, RpsK, and RpsL.

**Fig S11.**
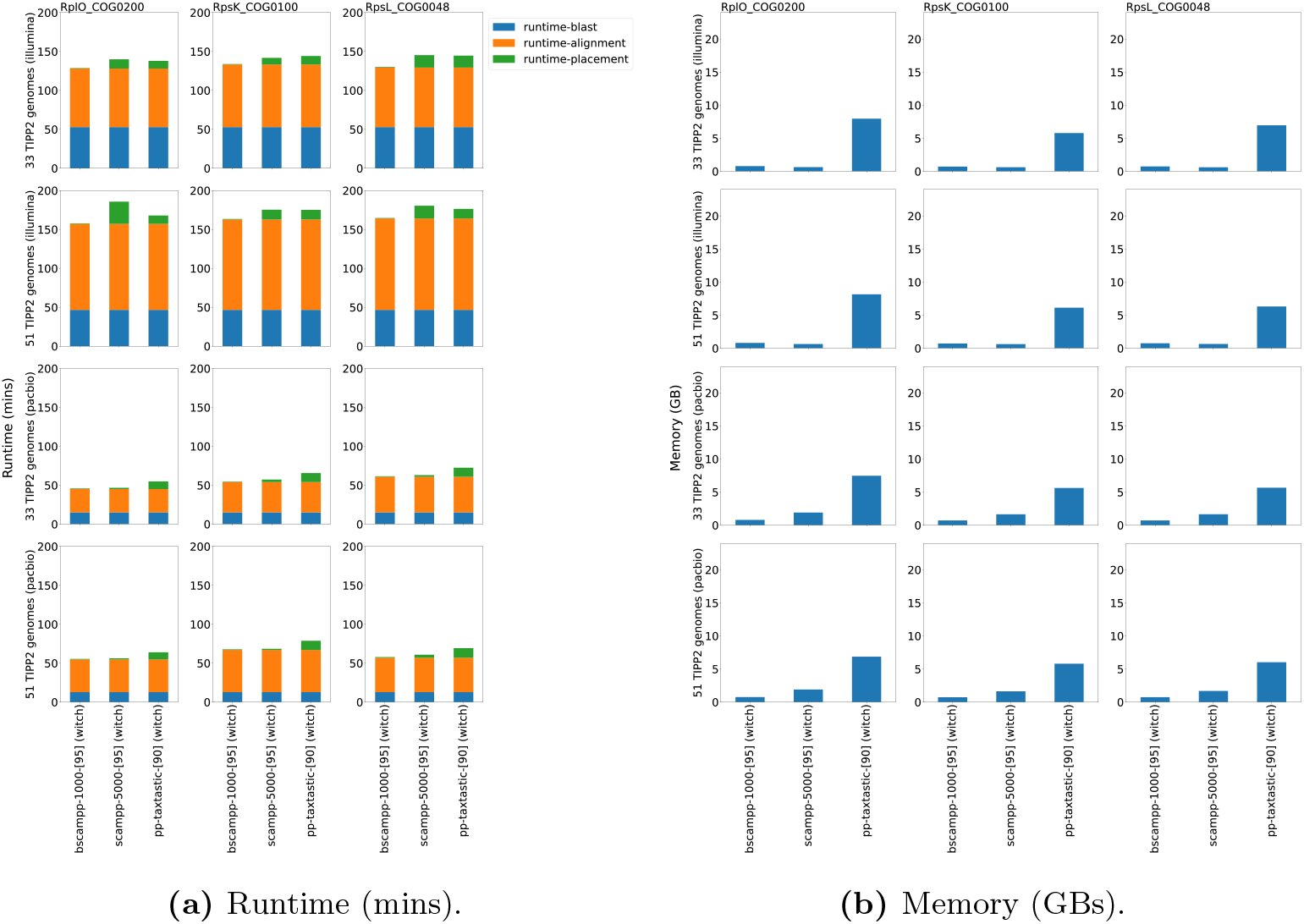
Runtime usage in minutes (left) and memory usage in GBs (right), using WITCH alignment, of the best variants of BSCAMPP, SCAMPP, and pplacer-taxtastic identifying Illumina and PacBio reads from two TIPP2 datasets (33 and 51 genomes) assigned to three marker genes, RplO, RpsK, and RpsL. Runtime for blast, alignment (by WITCH), and placement are shown. “runtime-alignment” denotes the time to obtain the read alignment by WITCH assigned to each marker gene. “runtime-placement” denotes the total time used to obtain placements of query reads assigned to each marker gene.

**Fig S12.**
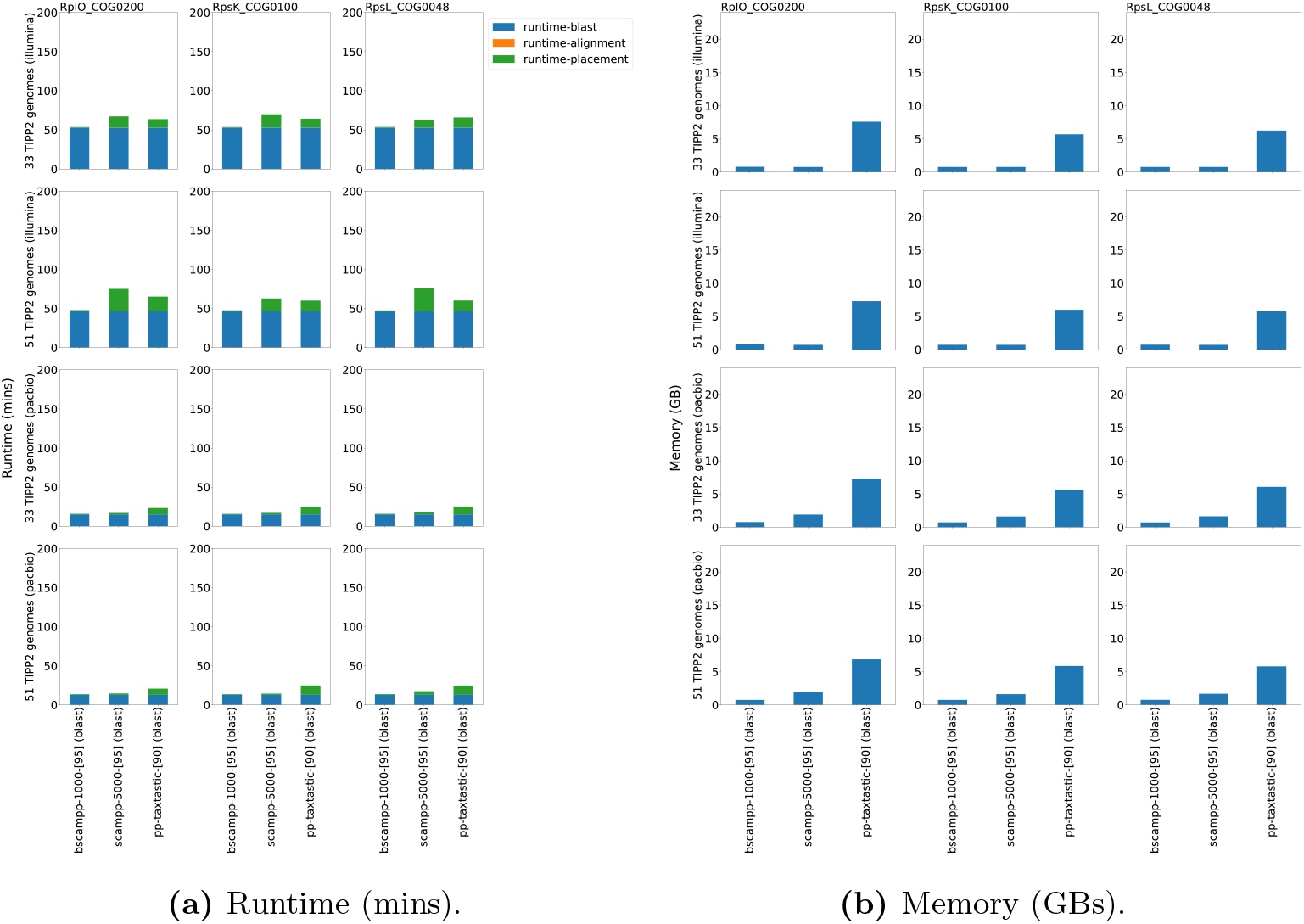
Runtime usage in minutes (left) and memory usage in GBs (right), using BLAST alignment, of the best variants of BSCAMPP, SCAMPP, and pplacer-taxtastic identifying Illumina and PacBio reads from two TIPP2 datasets (33 and 51 genomes) assigned to three marker genes, RplO, RpsK, and RpsL. Runtime for blast, alignment (by BLAST), and placement are shown. “runtime-alignment” denotes the time to post-process the BLAST output to obtain best-scored pairwise alignments (to reference genomes) of binned query reads.

#### S5.3 Marker gene selection

So far, we have only looked at three marker genes (RplO, RpsK, and RpsL) for abundance profiling. Here, we examined the accuracy of TIPP3 on each of the 40 marker genes when profiling Illumina reads from the TIPP2 dataset with 51 genomes. We then sorted the marker genes by their averaged normalized Hellinger distance across all taxonomic levels (Figure S13). Full results can be found in this sub-section.

We observed a wide gap between the best and the worst marker genes in terms of their profiling accuracy, particularly on the genus level and up. When using a subset of marker genes for aggregated abundance profiling, we found that excluding either the largest two marker genes (FtsY and RpoB, ranked by marker gene alignment sizes) or the bottom 20 marker genes generally resulted in higher profiling accuracy compared to using all 40 marker genes or just three marker genes (Figure S14). Hence, we removed FtsY and RpoB genes from the TIPP3 reference package and used the remaining 38 marker genes for the following experiments.

##### S5.3.1 Abundance profile on each marker gene

**Fig S13.**
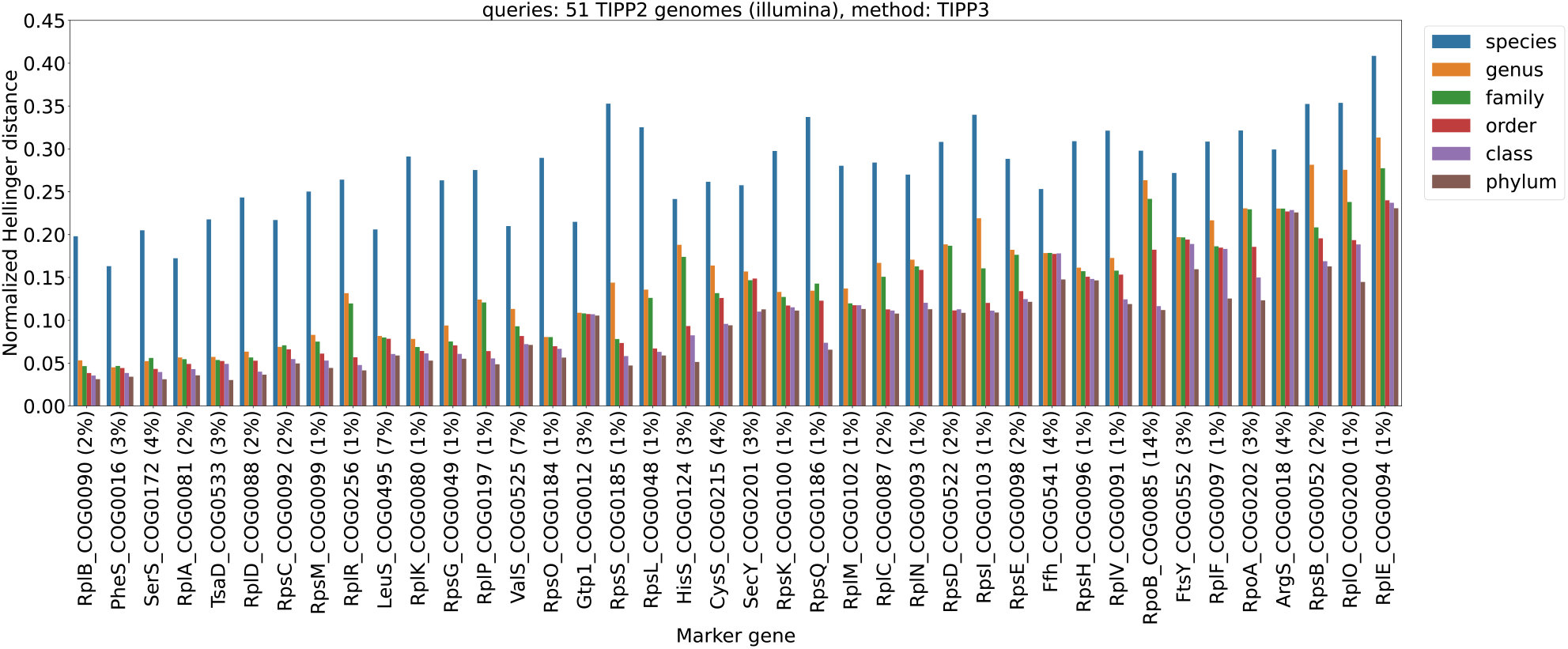
Individual marker gene abundance profile accuracy for TIPP3 on each taxonomic level. The query reads used are Illumina reads and generated from the TIPP2 dataset with 51 genomes. The x-axis denotes different marker genes and taxonomic levels (excluding superkingdom). The marker genes are sorted from left to right in order of average ranking of normalized Hellinger distance (i.e., the lower average normalized Hellinger distance a marker gene has, the closer it is to left on the x-axis). The “%” sign for each marker gene denotes the percentage of query reads assigned to that marker gene.

##### S5.3.2 Abundance profile using different sets of marker genes

**Fig S14.**
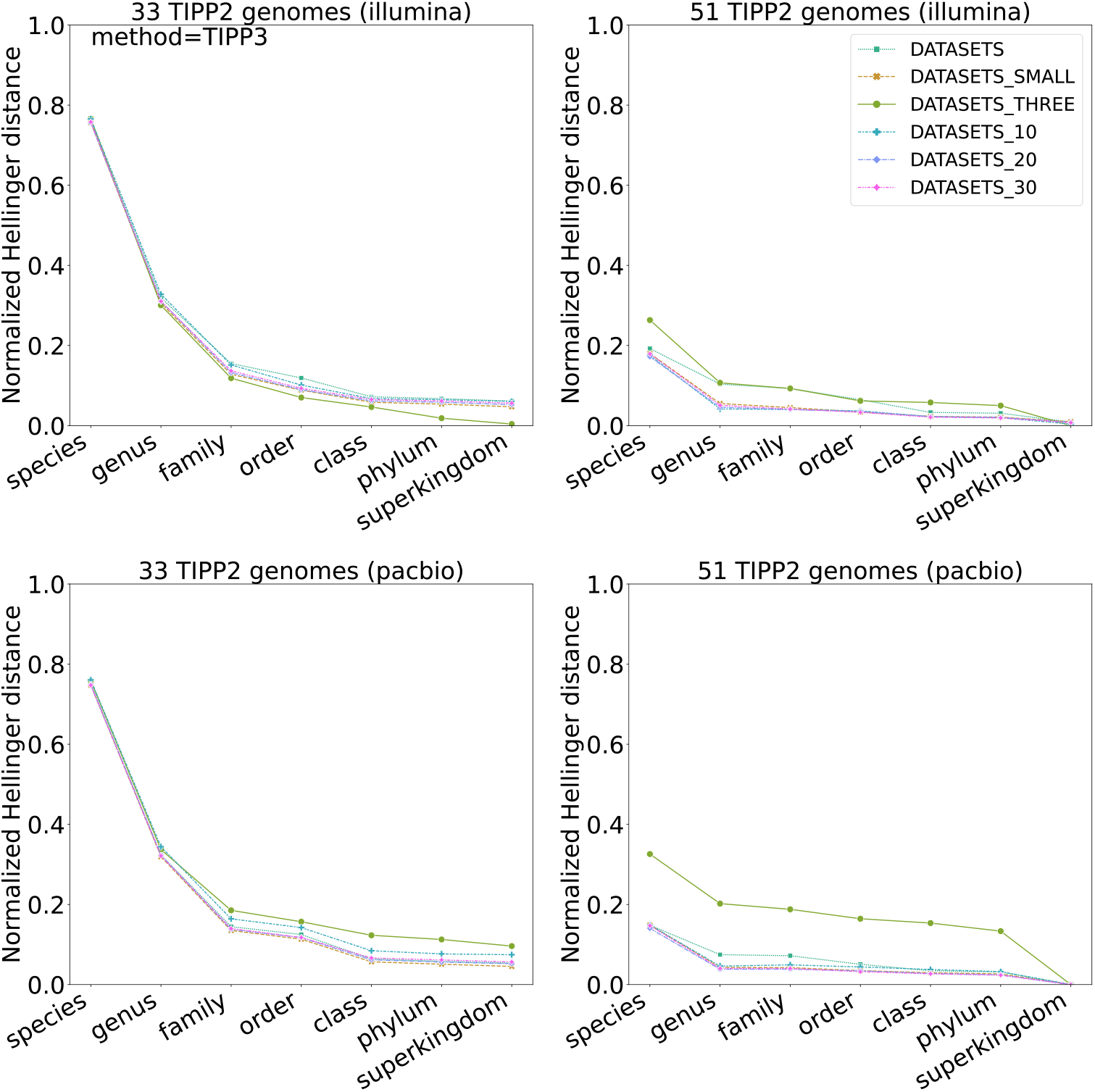
Abundance profiles of TIPP3 using different sets of marker genes. DATASETS denotes using all marker genes, DATASETS_SMALL for using all except RpoB and FtsY, DATASETS_THREE for using only RplO, RpsK, and RpsL, DATASETS_10 for using the top 10 marker genes denoted in Figure S13, DATASETS_20 for using the top 20, and DATASETS_30 for using the top 30 marker genes. Reads are either Illumina or PacBio generated from two TIPP2 datasets with 33 and 51 genomes.

### S6 Additional Results for Experiment 3: Evaluation of TIPP3 for abundance profiling

#### S6.1 Abundance profile comparison for all methods

**Fig S15.**
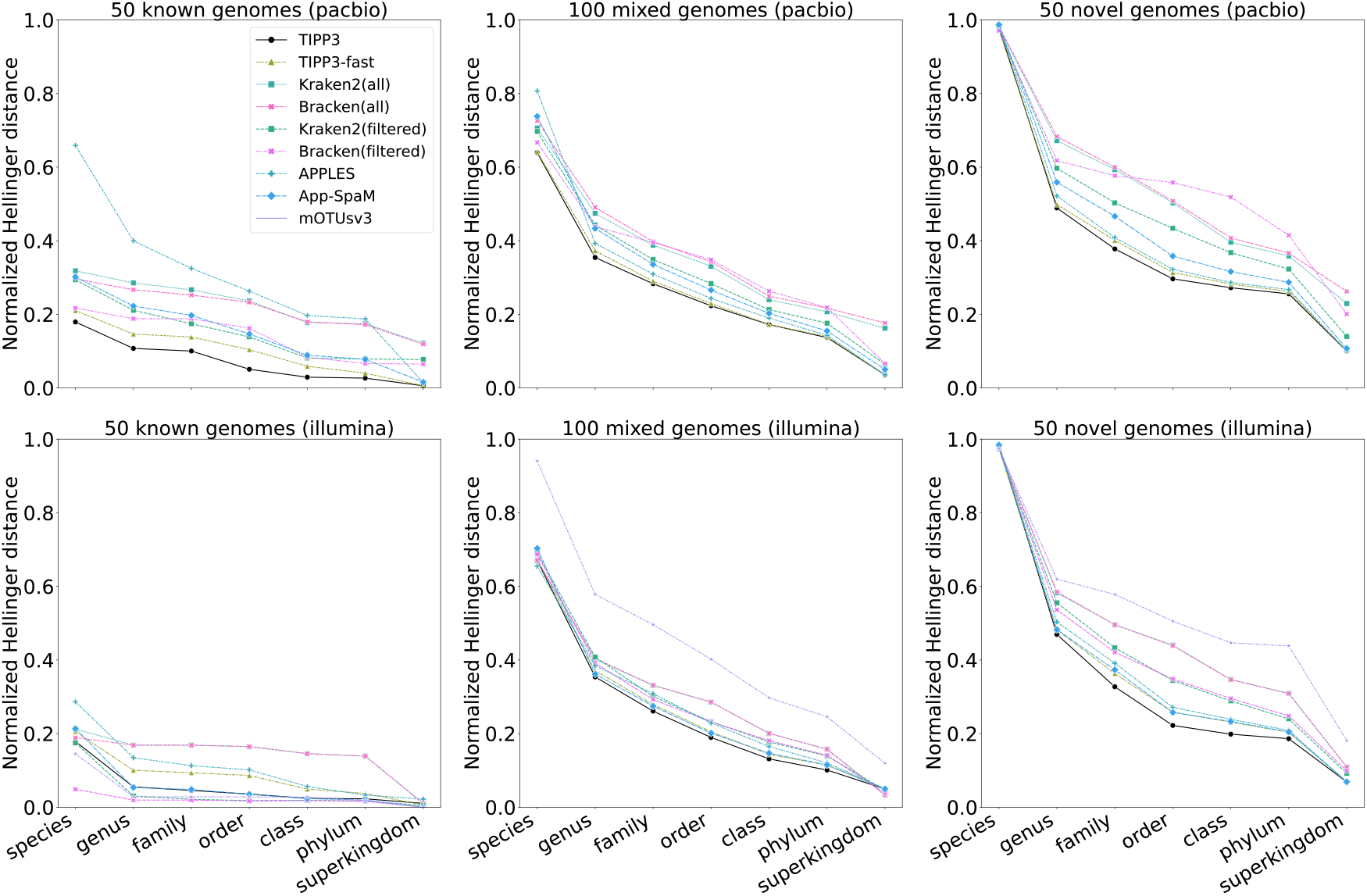
Abundance profiling accuracy by normalized Hellinger distance (lower means more accurate) of all methods/pipelines for PacBio and Illumina simulated reads from 50 known, 100 mixed, and 50 novel genomes. Abundance profiles are computed using 38 marker genes for TIPP3, TIPP3-fast, Bracken(filtered), Kraken2(filtered), APPLES, and App-SpaM. For PacBio read datasets, mOTUsv3 did not produce any classification and thus is not shown.

#### S6.2 Detailed evaluation on species and genus abundances

Genus- and Species-specific abundance estimation errors are shown. Species-specific errors are not shown for 50 novel genomes, since no methods should be able to produce any non-zero profile for the species by definition. The genus and species names and their corresponding reference abundances of the 50 known genomes are shown in Tables S1.

**Table S1.**
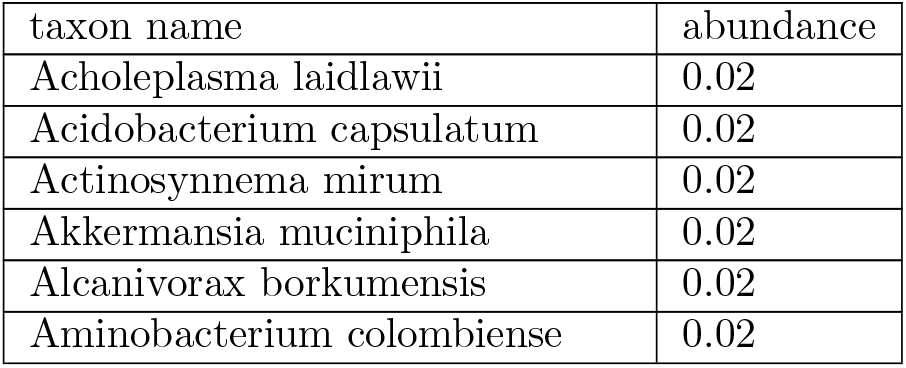
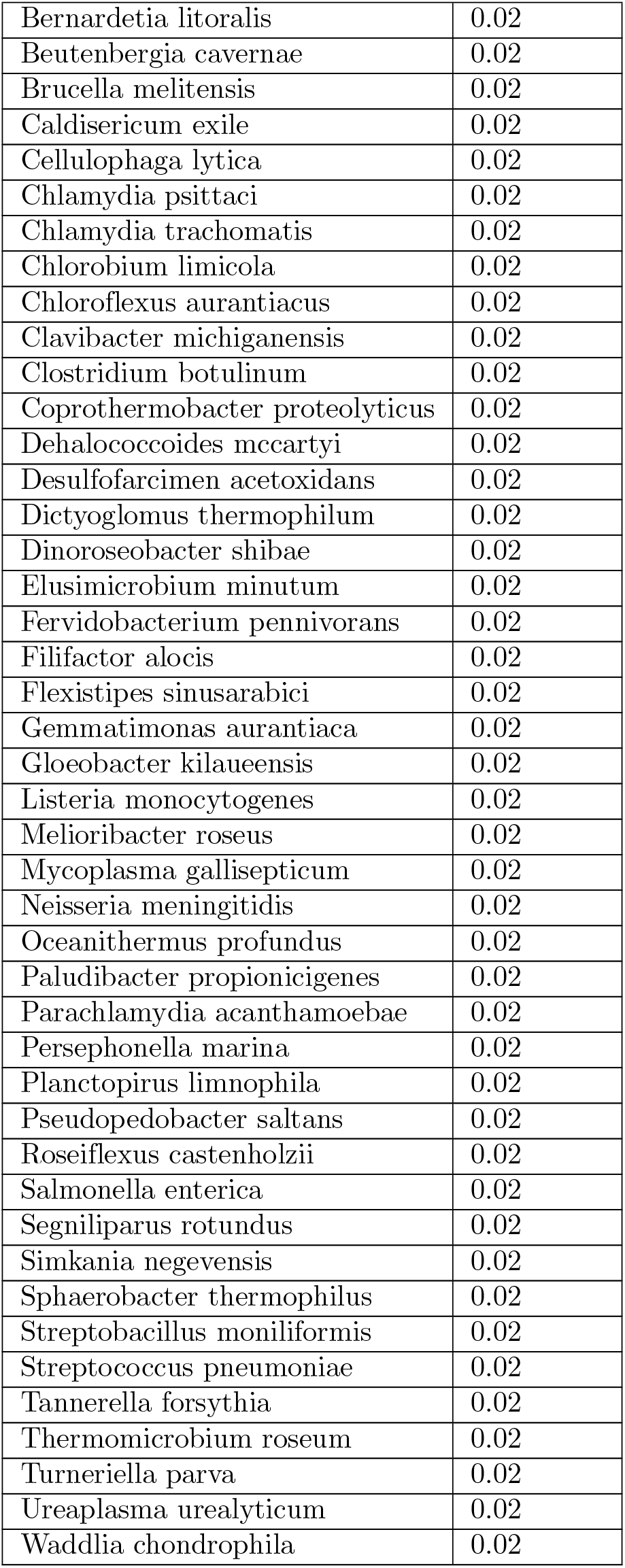
Taxon names and reference abundances of the 50 known genomes, sorted by alphabetical order.

**Fig S16.**
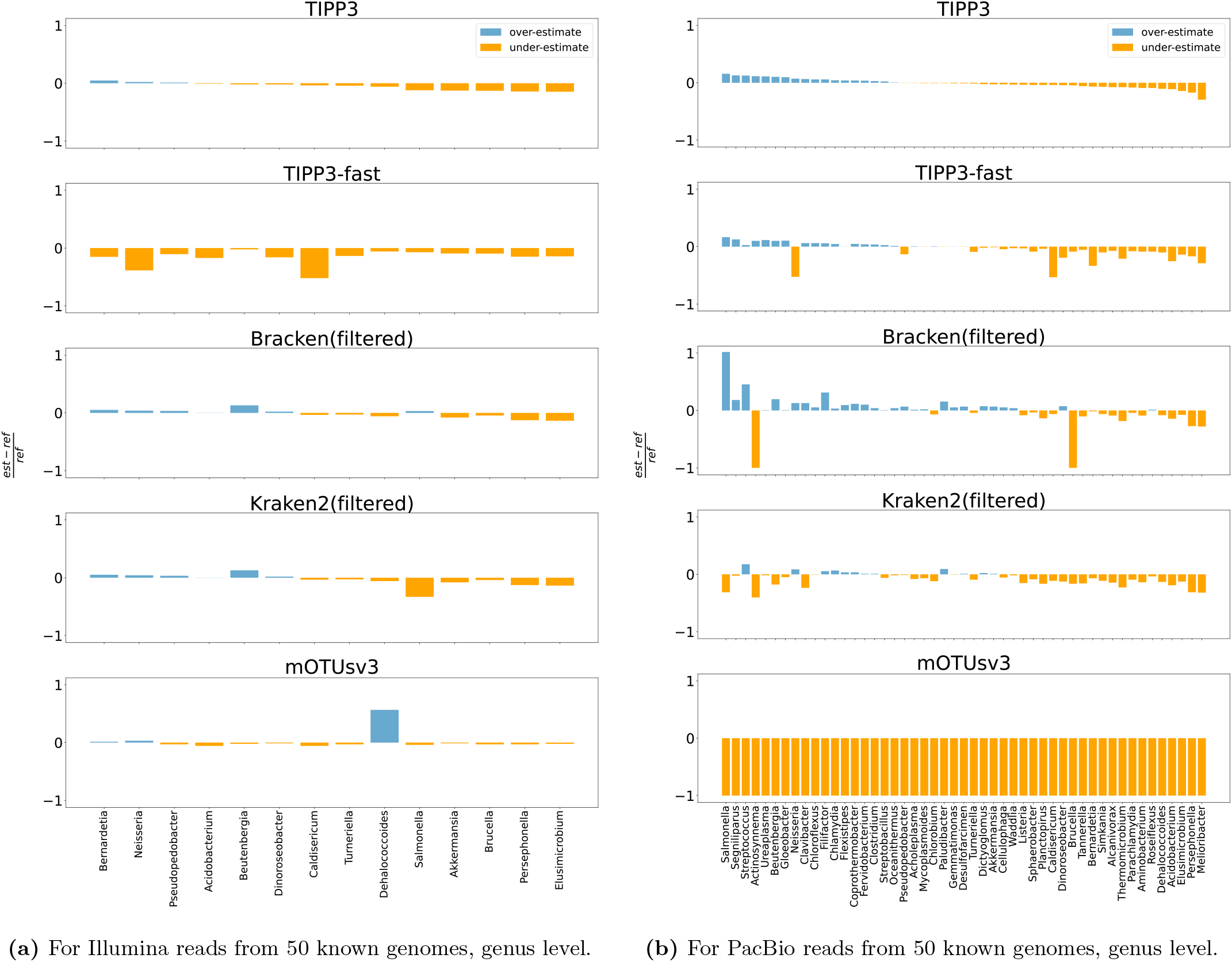
Genus-specific abundance estimation error for Illumina (left) and PacBio (right) reads of 50 known genomes of TIPP3, Bracken(filtered), Bracken(all), Kraken2(filtered), and Kraken2(all). The estimation error of a taxon is computed as the fractional difference between its estimated and reference compositions, shown on the Y-axis. Taxons are sorted by the estimation errors by TIPP3 in ascending order. A taxon group is shown if and only if it is present in the reference, and any of the methods has an estimation error *±*10%.

**Fig S17.**
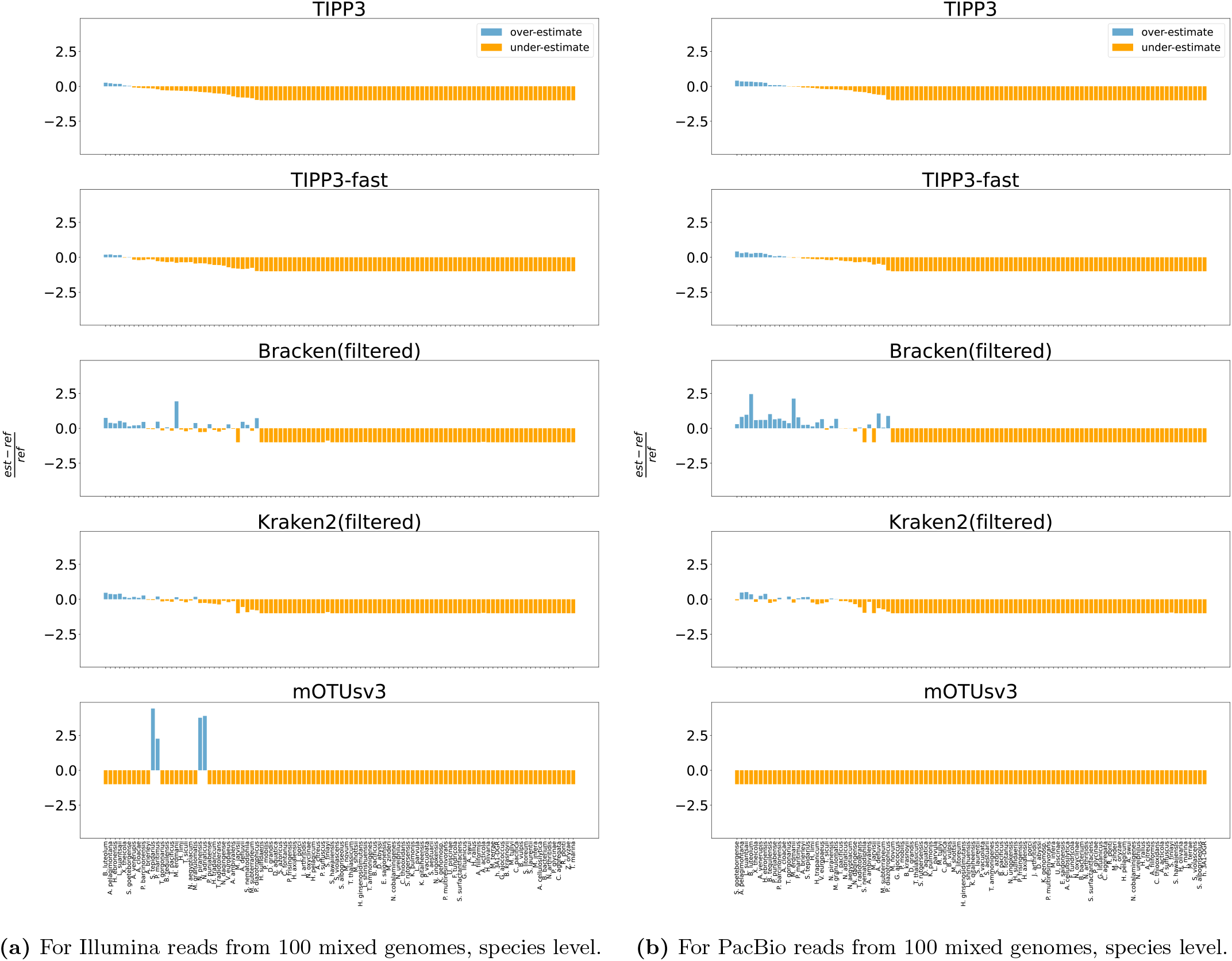
Species-specific abundance estimation error for Illumina (left) and PacBio (right) reads of 100 mixed genomes of TIPP3, Bracken(filtered), Bracken(all), Kraken2(filtered), and Kraken2(all). The estimation error of a taxon is computed as the fractional difference between its estimated and reference compositions, shown on the Y-axis. Taxons are sorted by the estimation errors by TIPP3 in ascending order. A taxon group is shown if and only if it is present in the reference, and any of the methods has an estimation error *±*10%.

**Fig S18.**
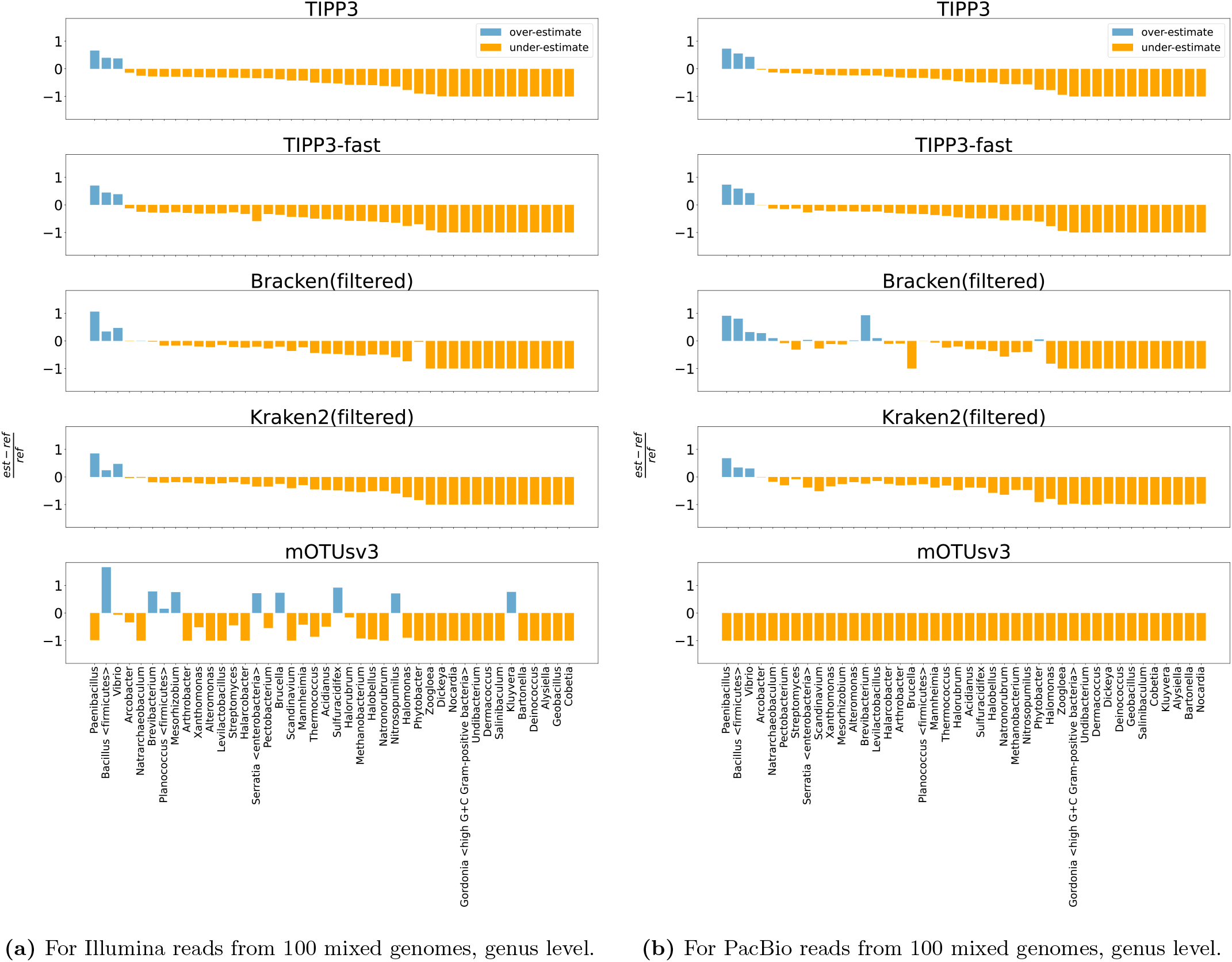
Genus-specific abundance estimation error for Illumina (left) and PacBio (right) reads of 100 mixed genomes of TIPP3, Bracken(filtered), Bracken(all), Kraken2(filtered), and Kraken2(all). The estimation error of a taxon is computed as the fractional difference between its estimated and reference compositions, shown on the Y-axis. Taxons are sorted by the estimation errors by TIPP3 in ascending order. A taxon group is shown if and only if it is present in the reference, and any of the methods has an estimation error *±*10%.

**Fig S19.**
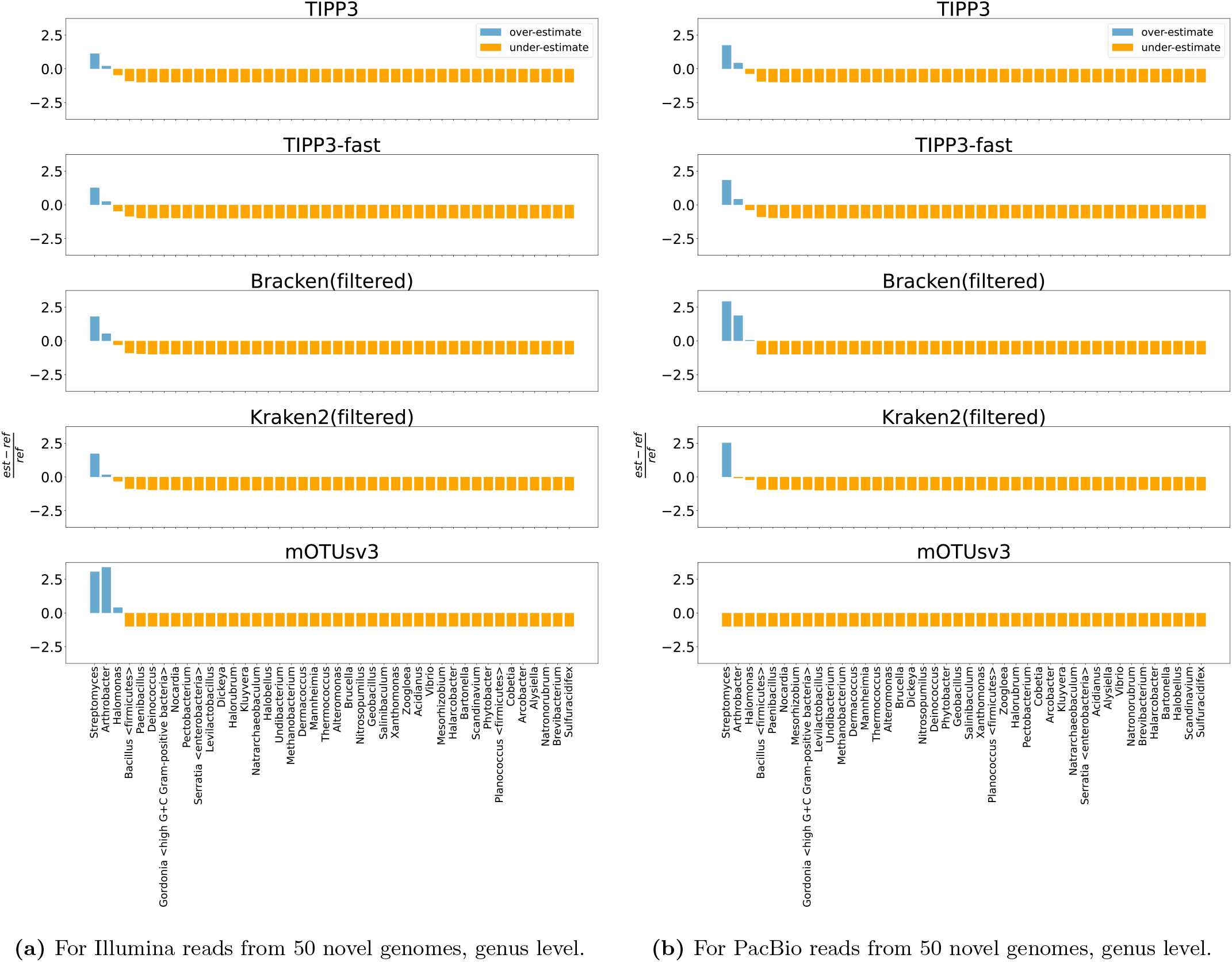
Genus-specific abundance estimation error for Illumina (left) and PacBio (right) reads of 50 novel genomes of TIPP3, Bracken(filtered), Bracken(all), Kraken2(filtered), and Kraken2(all). The estimation error of a taxon is computed as the fractional difference between its estimated and reference compositions, shown on the Y-axis. Taxons are sorted by the estimation errors by TIPP3 in ascending order. A taxon group is shown if and only if it is present in the reference, and any of the methods has an estimation error *±*10%.

#### S6.3 Runtime and memory usage

**Fig S20.**
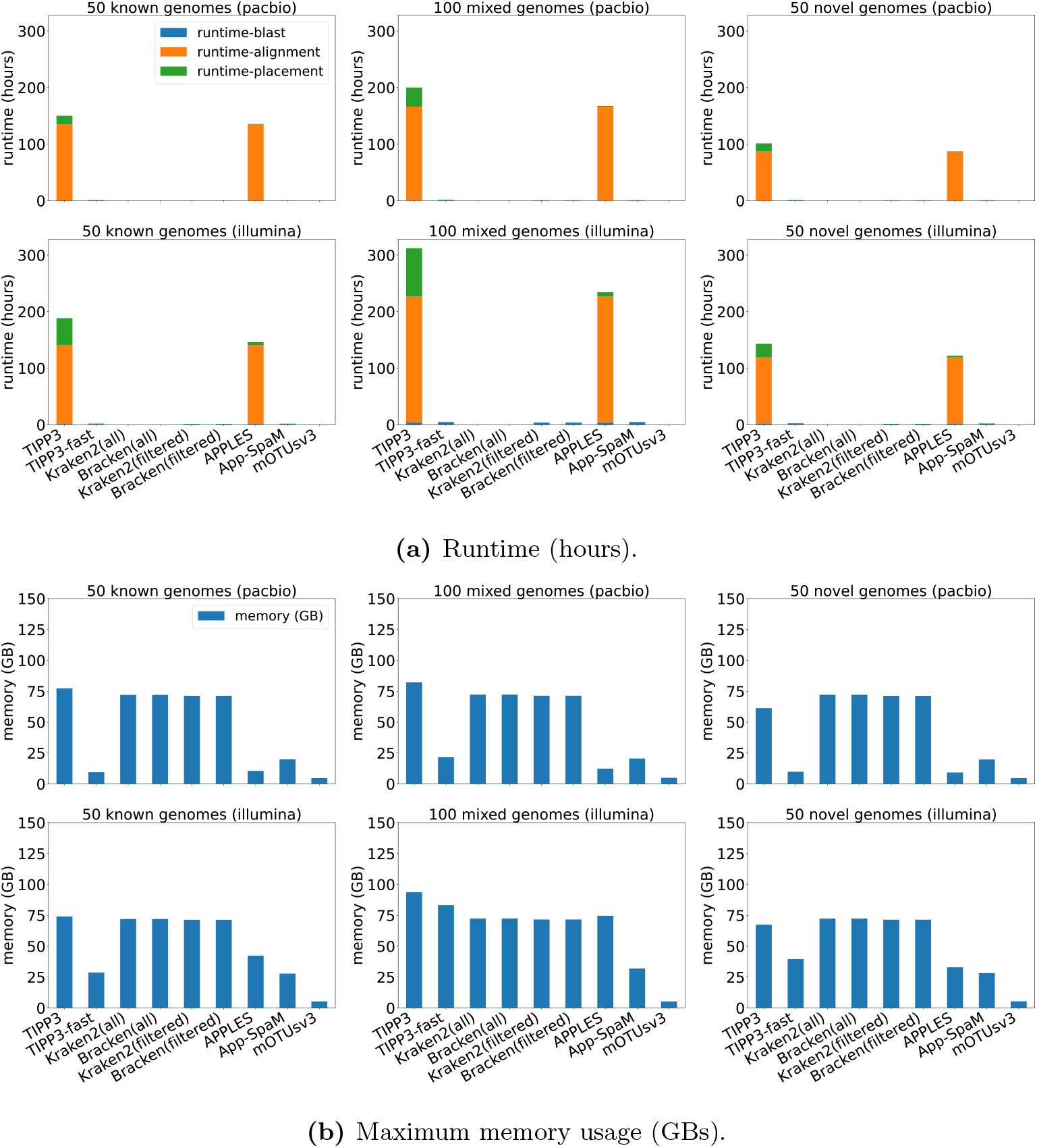
Runtime in hours (top) and maximum memory usage in GBs (bottom) of TIPP3, TIPP3-fast, Kraken2(filtered), Bracken(filtered), mOTUsv3, APPLES, and App-SpaM for PacBio and Illumina simulated reads from 50 known, 100 mixed, and 50 novel genomes.

#### S6.4 Improving TIPP3 runtime performance

A key observation on TIPP3 and other tested methods is that TIPP3 is among the slowest and uses a high amount of memory. To improve the runtime and memory usage of TIPP3, we explored different modifications to the TIPP3 pipeline by changing one or more steps. Each TIPP3 variant is denoted as “`<align method>+<placement method>`“ (using 38 marker genes) or “`<align method>+<placement method>+<Xmarker>`“ (using *X* marker genes). For example, default TIPP3 uses WITCH for query alignment and pplacer-taxtastic for query placement and is denoted as `witch+pplacer`. TIPP3-fast uses BLAST for query alignment and BSCAMPP for query placement and is denoted as `blast+bscampp`. We compare these variants of TIPP3 on their profiling accuracy, runtime, and memory usage on Illumina and PacBio reads of the testing datasets (50 known, 100 mixed, and 50 novel genomes).

We showed the best variants in terms of profiling accuracy in Figure S21 and the corresponding runtime/memory usage in Figure S22.

For reads from known genomes, all TIPP variants that use pplacer-taxtastic for read placement have similar profiling accuracy for Illumina or PacBio reads. Switching to BSCAMPP or SCAMPP for read placement leads to higher errors at all taxonomic levels observed on known reads. For the read alignment method, using WITCH results in slightly better accuracy than using BLAST, and the difference is more noticeable at lower taxonomic levels. One exception is at the species level for PacBio reads, where using BLAST is more accurate than using WITCH. For marker gene selection, using 20 instead of 38 marker genes does not impact profiling accuracy noticeably. 20 marker genes lead to slightly higher accuracy at the species level for both Illumina and PacBio reads, but we also observe slightly lower accuracy for family level and above of PacBio reads.

For reads from a mixture of known and novel genomes or entirely novel genomes, all TIPP3 variants have lower accuracy compared to known genomes. The relative performance resembled most observations on known genomes with a few differences. Switching to BSCAMPP or SCAMPP for read placement had similar accuracy to the other TIPP3 variants. Using 20 instead of 38 marker genes harmed profiling accuracy and was particularly noticeable on higher taxonomic levels (e.g., phylum and superkingdom).

**Fig S21.**
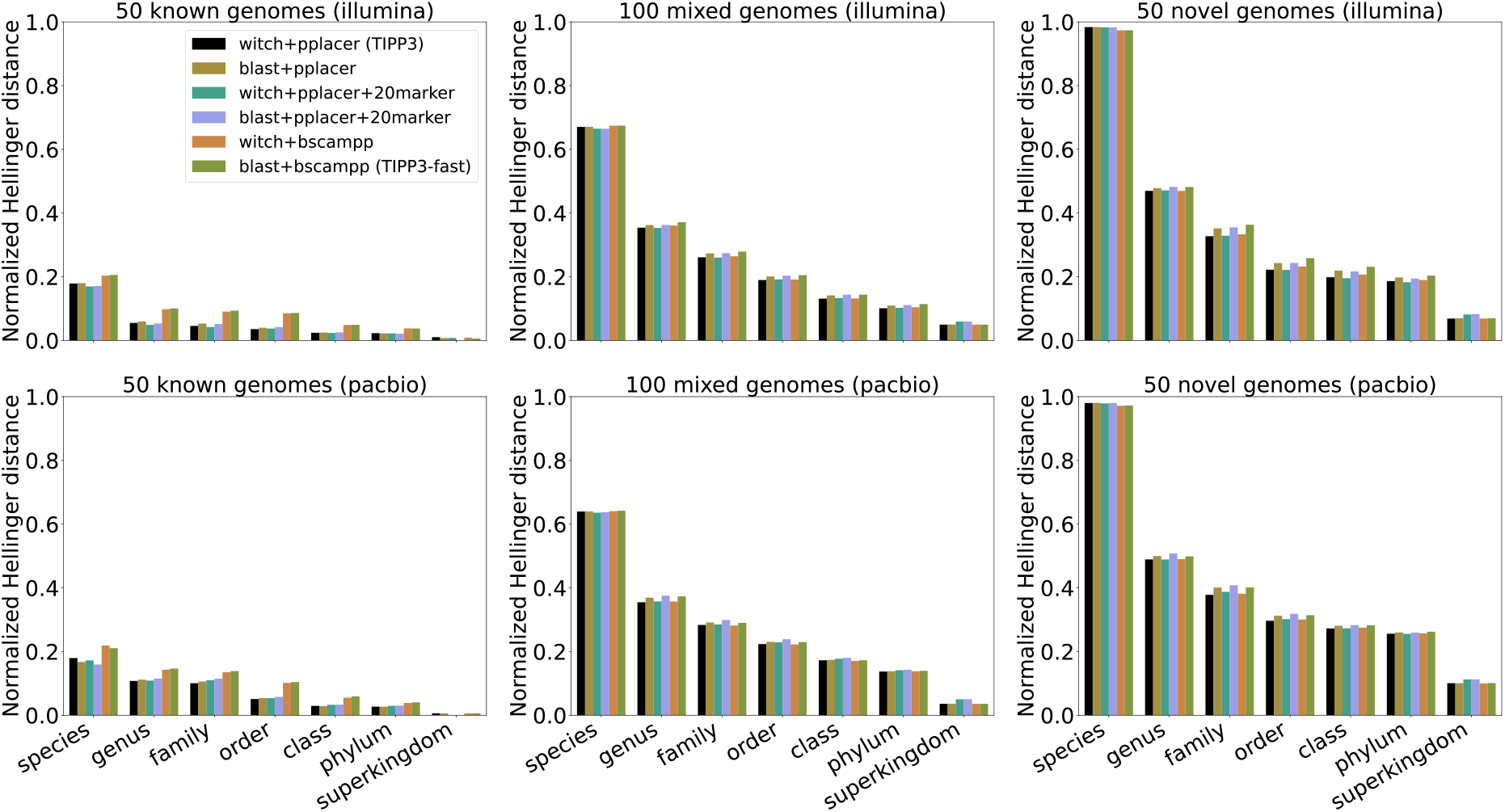
Abundance profile accuracy by normalized Hellinger distance of TIPP3 variants for Illumina and PacBio simulated reads from known, mixed, and novel genomes. Each TIPP3 variant is denoted as “`<align method>`+`<placement method>`“ (using 38 marker genes) or “`<align method>`+`<placement method>`+`<Xmarker>`“ (using *X* marker genes). For example, TIPP3 uses WITCH for query alignment and pplacer-taxtastic for query placement and is denoted as “`witch`+`pplacer`”.

**Fig S22.**
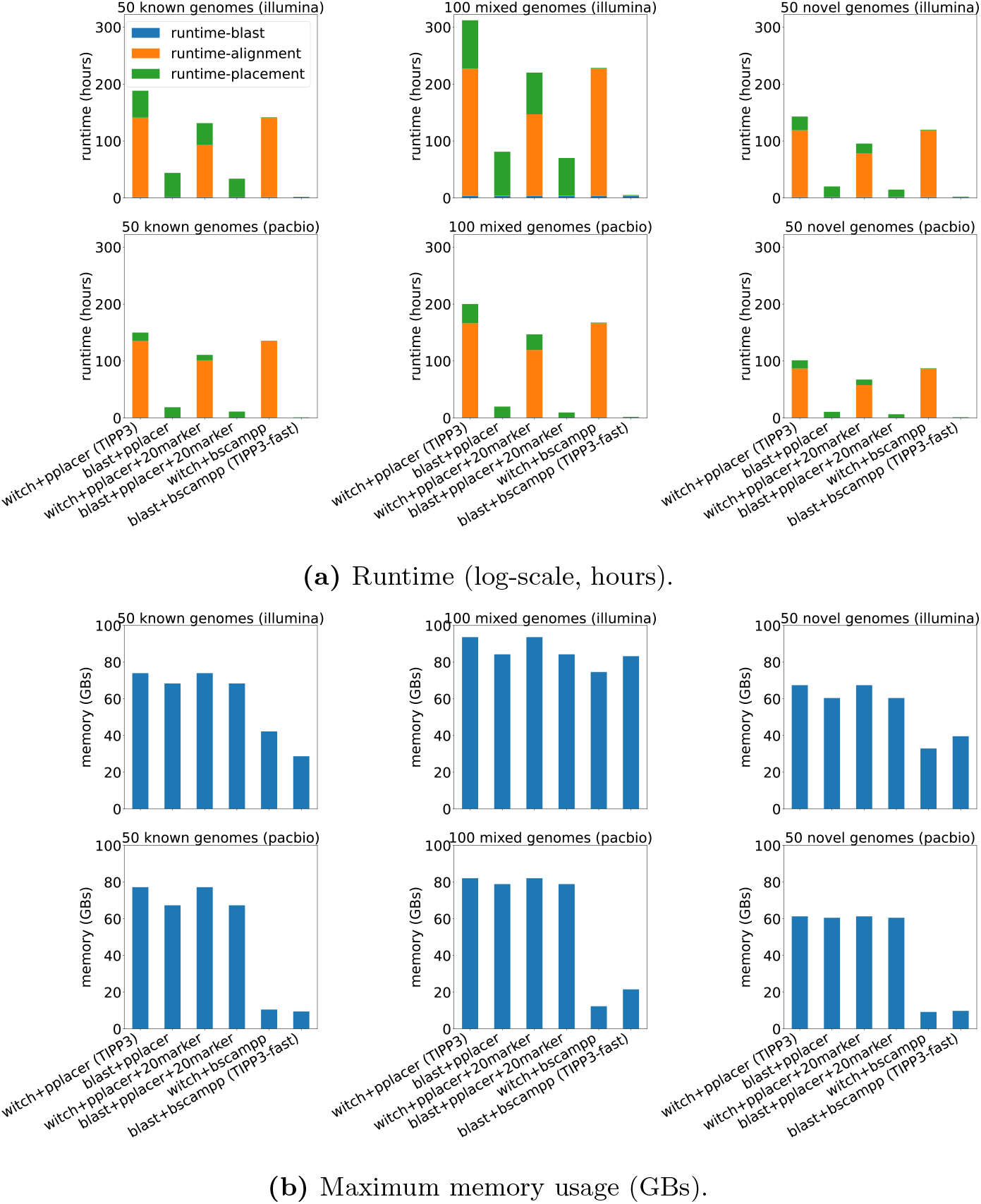
Runtime in hours (top) and peak memory usage in GBs (bottom) of all benchmarked methods/pipelines for Illumina and PacBio simulated reads from known, mixed, and novel genomes.

**Table S2.**
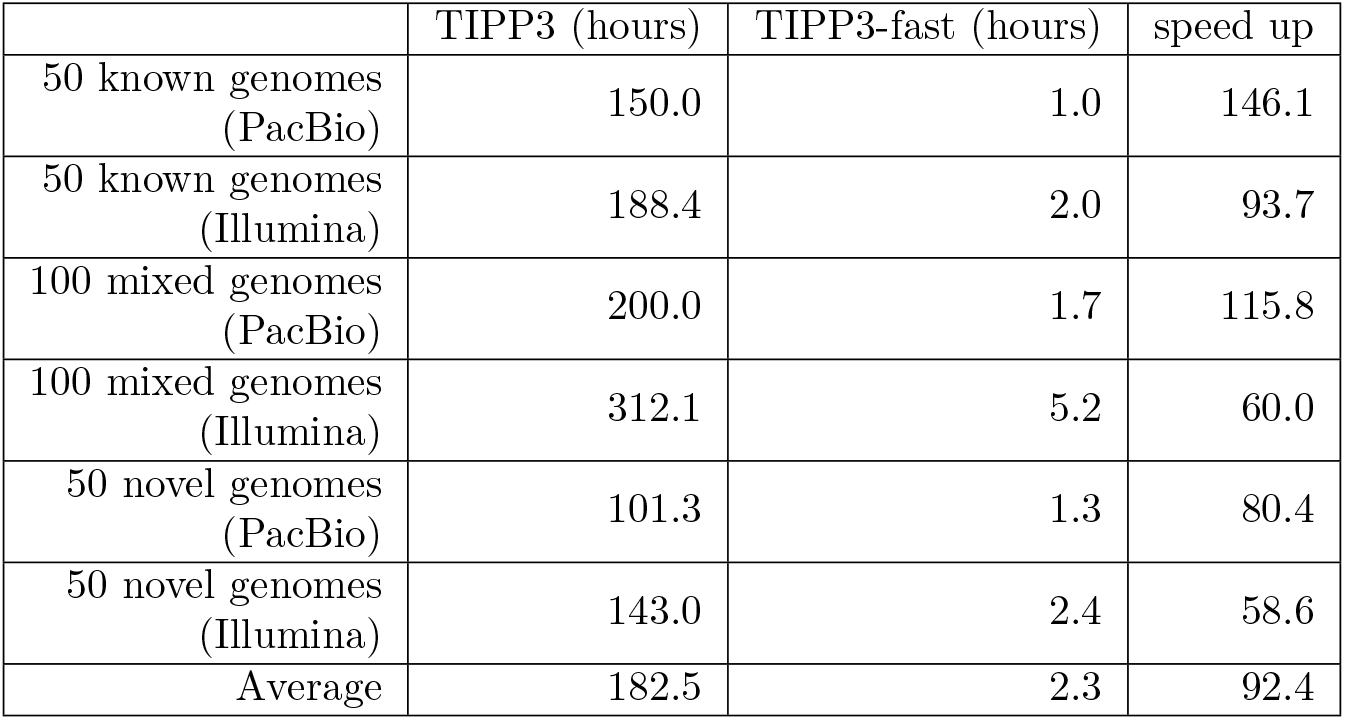
TIPP3 and TIPP3-fast runtime comparison in hours and the corresponding speedup by TIPP3-fast to TIPP3, on the six testing datasets; average results are shown in the last row.

