## Supplementary Materials for "TIPP3 and TIPP3-fast: Improved Abundance Profiling in Metagenomics"

#### Contents

**S1 Additional Information on TIPP3 Reference Packages 3**

**S2 Software Commands 4**

**S3 Additional Information for Read Generation 6**

**S4 Additional Information for Normalized Hellinger Distance 8**

### List of Tables

S1 Taxon names and abundances of the 50 known genomes . . . . . . . . . 24
S2 Testing: TIPP3-fast vs. TIPP3 runtime speedup . . . . . . . . . . . . . 33

### Supporting Materials

This document provides the supporting materials for TIPP3, including additional information for obtaining the reference package, software commands for benchmarking, information for input read generation and normalized Hellinger distance, and additional results.

#### S1 Additional Information on TIPP3 Reference Packages

We used the same NCBI RefSeq Bacteria and Archaea genomes as TIPP2 [1], downloaded in November 2019 from the NCBI RefSeq database. The TIPP3 reference package was then constructed using the following pipeline. We first extracted and cleaned the same set of 40 marker genes as TIPP2 from the RefSeq genomes. After filtering, we aligned each set of marker gene sequences with MAGUS [2] and built a taxonomy tree with RAxML [3]. For running pplacer [4] with the taxtastic package [5], we re-estimated tree branch lengths with FastTree-2 [6]. The latest TIPP3 reference package is available at `https://doi.org/10.13012/B2IDB-4931852_V1`. Scripts for data processing can be found at `https://github.com/shahnidhi/TIPP_reference_package`.

##### Pre-alignment data processing

1. Filter the set of Archaea and Bacteria genomes (a total of 173,240 genomes) for each marker gene by (1) removing sequences that do not have matched amino/nucleotide sequences, (2) removing sequences that are 3 standard deviations from the median length. (with `data/filterData.py`).
2. Update the taxids using the **NCBI taxonomy downloaded on July 5th, 2023** (with `taxit update_taxids`).
3. Create a new taxonomy table for each marker gene (with `taxit taxtable`).
4. Update species mapping (species name to taxid) (with `update_species_mapping.py`).
5. Build taxonomy for each marker gene (with `build_unrefined_tree.pl`).
6. The outputs, for each marker gene, are (1) a final filtered set of sequences ($\sim 55,000$ sequences), and (2) an unrefined taxonomy of the sequences.

##### Alignment

1. Perform a non-recursive MAGUS alignment [2] on the filtered sequences for each marker gene.
2. The exact command of non-recursive MAGUS:

```
$ magus.py -np 16 -i [filtered unaligned sequences] \
    --recurse false \
    -d [output directory] -o [output alignment path]
```

3. The output is an alignment for each set of marker gene sequences.

**Build taxonomy**

1. Back-translate the amino acid alignment to nucleotide alignment (with `backtranslate_refseq.py`).

2. Clean up the tree internal labels on the taxonomy before refining it (with `nw_topology -bI`).

3. Mask gappy sites (> 95% gaps) of the nucleotide alignment (with `ogcat mask -p 0.95`).

4. Resolve polytomies with RAxML [3] under the GTR+CAT model.

```
$ raxml-PTHREADS-AVX -s [masked alignment] -m GTRCAT \
    -n [output prefix] -g [cleaned taxonomy] \
    -T 24 -p 12345 -w [work directory]
```

5. Add branch length information to the RAxML refined tree.

```
$ raxml-PTHREADS-AVX -s [masked alignment] -m GTRCAT \
    -n [output prefix] -F -f e [RAxML refined tree] \
    -T 24 -p 54321 -w [work directory]
```

6. Re-estimate the branch lengths with RAxML-ng [7] under the GTR+GAMMA model.

```
$ raxml-ng --evaluate --msa [masked alignment] \
    --model GTR+G --tree [RAxML branch-length tree] \
    --brlen scaled --redo --force perf_threads \
    --prefix [output prefix] --threads 24
```

7. Add back the internal labels with taxonomic information (with `relabel-modified.py`).

**Re-estimating numeric parameters** For `pplacer` [4] running with the `taxtastic` Python package [5], we re-estimated the taxonomy's numeric parameters with FastTree-2 [6] using the following command:

```
$ FastTreeMP --nosupport -gtr -gamma -nt -mllen -nome \
    -log [FastTree-2 log] -intree [inferred taxonomy] \
    < [masked alignment] > [re-estimated taxonomy]
```

Then, a `taxtastic` package is created as:

```
$ taxit create -P [output package dir] -l [name] \
    --aln-fast [masked alignment] \
    --tree-file [re-estimated taxonomy] \
    --tree-stats [FastTree-2 log]
```

### S2 Software Commands

We ran all software with 16 CPU cores and up to 256 GB of memory, running each software with no time limit until completion.

1. We ran Kraken2 (v2.1.3) with the following command:

```
$ kraken2 --db [Kraken2 database] \
    --threads 16 --report [Kraken2 report] \
    [query reads file] > [Kraken2 output]
```

2. We ran Bracken (v2.9) with the following command (using Kraken2 output):

```
$ bracken -d [Bracken database] -i [Kraken2 report] \
    -w [Bracken report] -t 10 -r 150 \
    -o [Bracken output]
```

3. We ran mOTUsv3 (v3.1.0) with the following command:

```
$ motus profile -db [mOTUs database] \
    -q -u -p -c -s [query reads file] \
    -CC 16 -t 16 -o [mOTUs output]
```

4. We ran APPLES-2 (v2.0.11) with the following command (for each marker gene and its assigned reads):

```
$ run_apples.py -t [marker gene taxonomy] \
    -s [marker gene alignment] -q [query reads file] \
    -T 16 -o [placement results] -X -D
```

5. We ran App-SpaM (v1.03) with the following command (for each marker gene and its assigned reads):

```
$ appspam -t [marker gene taxonomy] \
    -s [marker gene sequences] -q [query reads file] \
    --threads 16 -o [placement results]
```

6. We ran SCAMPP (v2.0.1) with the following command (for each marker gene and its assigned reads):

```
$ pplacer-SCAMPP.py -b [subtree size] \
    -i [marker gene taxonomy RAxML info] \
    -t [marker gene taxonomy] -d [output dir] \
    -o [placement results] -a [query read alignment] \
    --threads 16
```

7. We ran BSCAMPP (v1.0.0) with the following command (for each marker gene and its assigned reads):

```
$ EPA-ng-BSCAMPP.py -b [subtree size] \
    -i [marker gene taxonomy RAxML info] \
    -t [marker gene taxonomy] -d [output dir] \
    -o [placement results] -a [query read alignment] \
    --threads 16
```

8. We ran pplacer (v1.1.alpha19-0-g807f6f3) with the `taxtastic` (v0.10.0) package with the following command (for each marker gene and its assigned reads):

```
// re-estimating marker gene taxonomy branch lengths with FastTree-2
$ FastTreeMP -nosupport -gtr -gamma -nt \
    -mllen -nome -log [FastTree-2 log file] \
    -intree [marker gene taxonomy]

// getting taxtastic reference package
$ taxit create -P [taxtastic reference package] \
    -l [marker gene name] \
    --aln-fast [marker gene alignment] \
    --tree-file [FastTree-2 re-estimated tree] \
    --tree-stats [FastTree-2 log file]

// running pplacer
$ pplacer -m GTR -c [taxtastic reference package] \
    -o [placement results] -j 16 \
    [query read alignment]
```

### S3 Additional Information for Read Generation

Reads are generated for each set of reference genomes using either the ART sequence simulator [8] or PBSIM [9]. For a given set of genome sequences, query reads are simulated in two ways: Illumina (150bp) or PacBio ($\sim$ 3000bp). The commands used are shown below.

**Illumina reads generation** Illumina reads are simulated with `art_illumina` v2.5.8 with the following command. The sequence length is 150 bp and the coverage is 20x.

```
$ art_illumina -ss HS25 -sam \
     -i [genome path] -p -l 150 -f 20 -m 200 \
     -s 10 -o train_dataset2
```

**PacBio reads generation** PacBio reads are simulated with `pbsim` with the following command. The model used is CLR, and the average sequence length is set to 3000bp with a minimum length of 400bp. The average accuracy is set to 0.78 with a standard deviation of 0.07. According to the authors [9], the simulated data generated by `pbsim` with the CLR model and a 0.78 accuracy has 3.23% substitution rate, 10.53% insertion rate, and 3.98% deletion rate (obtained by aligning simulated reads to reference using LAST [10]). The coverage is also set to 20x.

```
$ pbsim [genome path] --prefix [output prefix] \
         --depth 20 --length-min 400 --length-mean 3000 \
         --accuracy-mean 0.78 --accuracy-sd 0.07 \
         --seed 522170 --model_qc [CLR model]
```

**Genome accession numbers for reads generation**

1. **TIPP2 dataset 1 (33 genomes)**:

### S4 Additional Information for Normalized Hellinger Distance

#### S4.1 Normalized Hellinger distance

Remember that regular Hellinger distance is given by:

$$H_l = \frac{\sqrt{\sum_{x \in C_l} (\sqrt{T_x} - \sqrt{E_x})^2}}{\sqrt{2}},$$

where $T_x$ is the true abundance and $E_x$ the estimated abundance of a clade $x$, for each $x$ in the set of clades $C_l$ on a taxonomic level $l$. Reads that are unclassified at a certain level are not counted for the Hellinger distance calculation.

**Theorem S4.1** (Theoretically Worst Hellinger Distance)**.** *If an abundance profiling method classifies $n$ reads in total, and $n_l$ reads are assigned with taxonomic labels at the taxonomic level $l$, then the theoretically worst Hellinger distance that the method can have at taxonomic level $l$ is $H'_l = \frac{\sqrt{1+\frac{n_l}{n}}}{\sqrt{2}}$.*

*Proof.* In the worst case, an estimated profile is disjoint from the true profile at taxonomic level $l$. More formally this means that for any clade $x \in C_l$, where $C_l$ is the union of labels in the estimated and true profiles, $T_x = 0$ or $E_x = 0$, exclusively. This also means $(\sqrt{T_x} - \sqrt{E_x})^2$ is either $T_x$ or $E_x$, depending on if clade $x$ is defined in the estimated or the true profile.

While the true profile sums to 1 by definition (i.e., taxonomic labels are known for reads), only $\frac{n_l}{n}$ of the estimated profile counts toward the Hellinger distance computation at level $l$. Then, we can rewrite the Hellinger distance computation as:

$$\begin{aligned} H'_l &= \frac{\sqrt{\sum_{x \in C_l} (\sqrt{T_x} - \sqrt{E_x})^2}}{\sqrt{2}} \\ &= \frac{\sqrt{\sum_{x \in C_l} T_x + \sum_{x \in C_l} E_x}}{\sqrt{2}} \\ &= \frac{\sqrt{1 + \frac{n_l}{n}}}{\sqrt{2}} \end{aligned}$$

□

By Theorem S4.1, the Hellinger distance $H_l$ of an estimated profile is bounded by $0 \leq H_l \leq H'_l$ at a taxonomic level $l$, and $H'_l \leq 1$. Only when all classified reads are assigned taxonomic labels at level $l$ ($n_l = n$), the range is from 0 to 1. Therefore, when $H_l$ is used to measure a method for abundance profiling accuracy, it would be biased if the method fails to classify all reads at level $l$.

A simple alternative is to measure profiling accuracy by $H^*_l = H_l / H'_l$, normalizing the measurement range to $[0, 1]$ and being independent of the number of reads classified at a taxonomic level for each method ($n_l$). We refer to this measurement as **normalized Hellinger distance**.

$$H_l^* = \frac{H_l}{H_l'} = \frac{\sqrt{\sum_{x\in C_l}(\sqrt{T_x}-\sqrt{E_x})^2}}{\sqrt{1+\frac{n_l}{n}}}$$

##### S4.2 Example for biases with Hellinger distance

Regular Hellinger distance is biased when the profile has a high fraction of unclassified reads. For example, let $n$ be the total number of reads classified. At the species level, let a method $A$ in total estimate 5 types of species $[s_1, s_2, s_3, s_4, s_5]$, where $s_i = 0.1, i = 1, 2, 3, 4, 5$. Then, the proportion of classified reads at taxonomic level $l$ for method $A$ is $n_{l,A} = 0.5$ and $n_{l,A}/n = 0.5$, with the other 0.5 as "unclassified/unspecified". In calculating Hellinger distance $H_{l,A}$ between the profile of method $A$ to the reference profile at level $l$, only the 0.5 that is classified/specified is used. Hence, the theoretically worst Hellinger distance $H'_{l,A}$ that method $A$ can have is to have a disjoint profile to the reference, meaning that $H'_{l,A} = \sqrt{1+0.5}/\sqrt{2} = 0.866$. In other words, given a method's estimated profile, $H'_l$ is the actual upper bound for $H_l$.

In the most extreme case when a method $B$ fails to produce *any classified groups* and has 1.0 as unclassified, then $H'_{l,B} = H_{l,B} = \sqrt{1+0}/\sqrt{2} = 0.707$. Let's assume that the reference profile looks like $[s_{100} = 0.7, s_{150} = 0.3]$. Another method $C$ may make some meaningful estimation, such as $[s_1 = 0.1, s_2 = 0.1, s_3 = 0.1, s_4 = 0.1, s_{100} = 0.05]$, as it gets only one group overlapping with the reference. If we compute the Hellinger distance for method $C$, we will get:

$$\begin{aligned} H_{l,C} &= \frac{\sqrt{\sum_{x\in C_l}(\sqrt{T_x}-\sqrt{E_x})^2}}{\sqrt{2}} \\ &= \frac{\sqrt{0.3 + 0.1\cdot 4 + (\sqrt{0.7}-\sqrt{0.05})^2}}{\sqrt{2}} \\ &= 0.733 \end{aligned}$$

We can see that although method $B$ does not give any estimation, it still gets a lower Hellinger distance than method $C$, which indicates a bias of Hellinger distance that favors making no estimation.

On the other hand, if we compare the normalized Hellinger distances of the two methods, we have:

$$\begin{aligned} H_{l,B}/H'_{l,B} &= 0.707/0.707 = 1 \\ H_{l,C}/H'_{l,C} &= 0.733/(\sqrt{1.45}/\sqrt{2}) = 0.861 \end{aligned}$$

where we have method $C$ having a lower error, which makes more sense.

#### S5 Additional Results for Experiment 1: Designing TIPP3

##### S5.1 Adding reads to marker gene alignments

We obtained UPP and WITCH alignments of reads binned to marker genes RplO, RpsK, and RpsL. WITCH was run with different parameter settings, varying the number of hidden Markov models (HMMs) for aligning each read and the decomposition subset sizes (i.e., how many sequences to include in a subset to build an HMM on). Then, batch-SCAMPP (BSCAMPP) [11] was used to obtain placement and taxonomic identification of each read in each marker gene taxonomy.

Based on overall taxonomic identification accuracy, we selected WITCH with one hidden Markov model subset and subset size ranging from 10 to 1000 sequences as our query read alignment method. Full results for the alignment benchmarks can be found below.

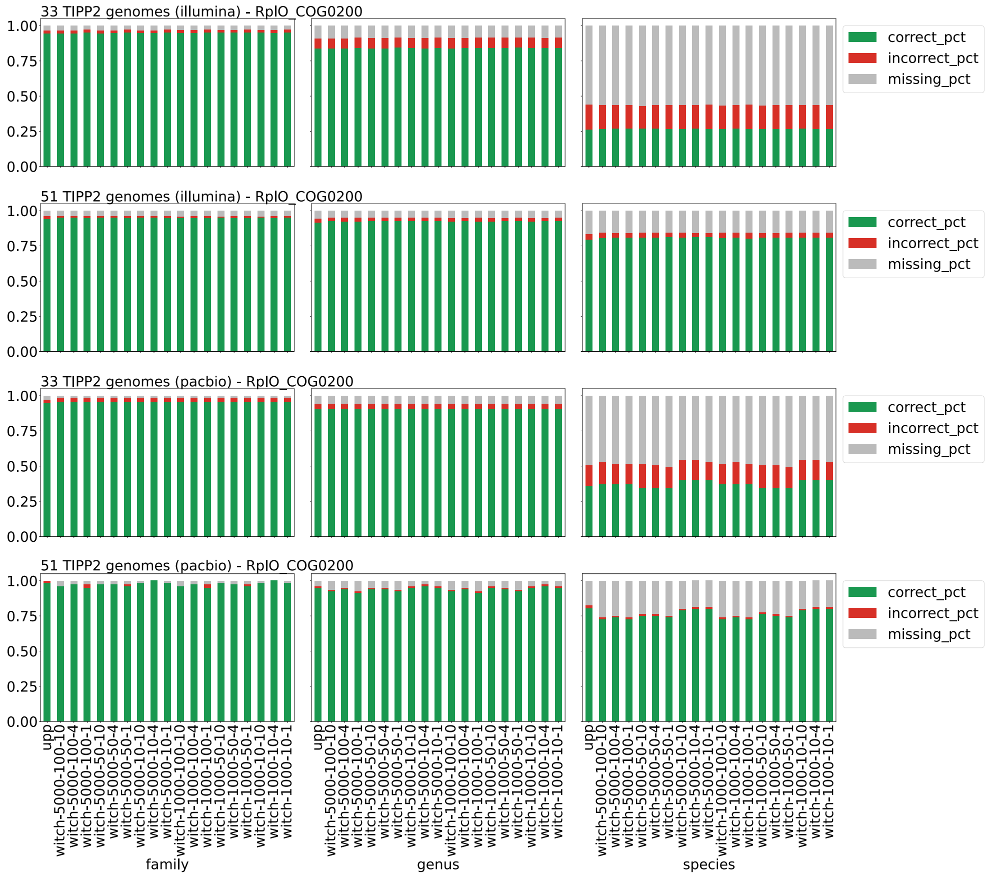

**Fig S1.** Taxonomic identification accuracy of UPP and different WITCH alignment variants on marker gene RplO_COG0200, for Illumina and PacBio reads from two TIPP2 datasets with 33 and 51 genomes (Family, Genus, and Species levels). The top two panels are for Illumina reads, and the bottom two are for PacBio reads. Taxonomic identification is done by performing query placements with Batch-SCAMPP with a subtree size of 1000 and a support value of 95%. WITCH variants are referred to as **witch-Z-A-k**, for which $Z$ denotes the upper bound for subset size, $A$ the lower bound, and $k$ the number of subsets to align a query read. "correct_pct" denotes the fraction of correctly identified reads (at a taxonomic level), "incorrect_pct" denotes the fraction of incorrectly identified reads, and "missing_pct" denotes the fraction of not-identified reads. The three fractions sum to 1 for each bar.

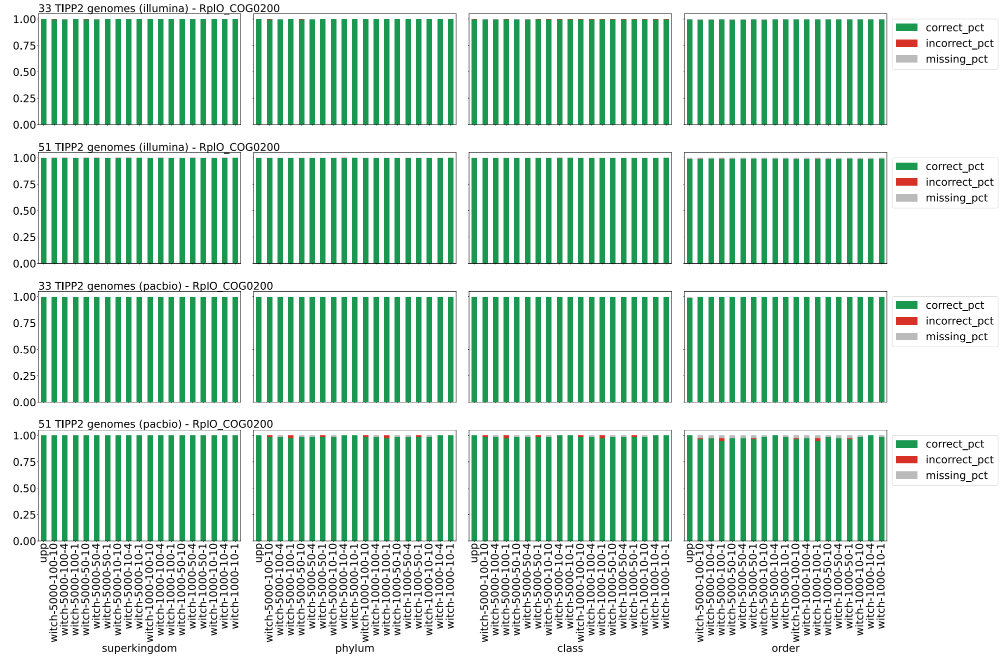

**Fig S2.** Taxonomic identification accuracy of UPP and different WITCH alignment variants on marker gene RplO_COG0200, for Illumina and PacBio reads from two TIPP2 datasets with 33 and 51 genomes (Superkingdom, Phylum, Class, and Order levels). The top two panels are for Illumina reads, and the bottom two are for PacBio reads. Taxonomic identification is done by performing query placements with Batch-SCAMPP with a subtree size of 1000 and a support value of 95%. WITCH variants are referred to as **witch-Z-A-k**, for which $Z$ denotes the upper bound for subset size, $A$ the lower bound, and $k$ the number of subsets to align a query read. "correct_pct" denotes the fraction of correctly identified reads (at a taxonomic level), "incorrect_pct" denotes the fraction of incorrectly identified reads, and "missing_pct" denotes the fraction of not-identified reads. The three fractions sum to 1 for each bar.

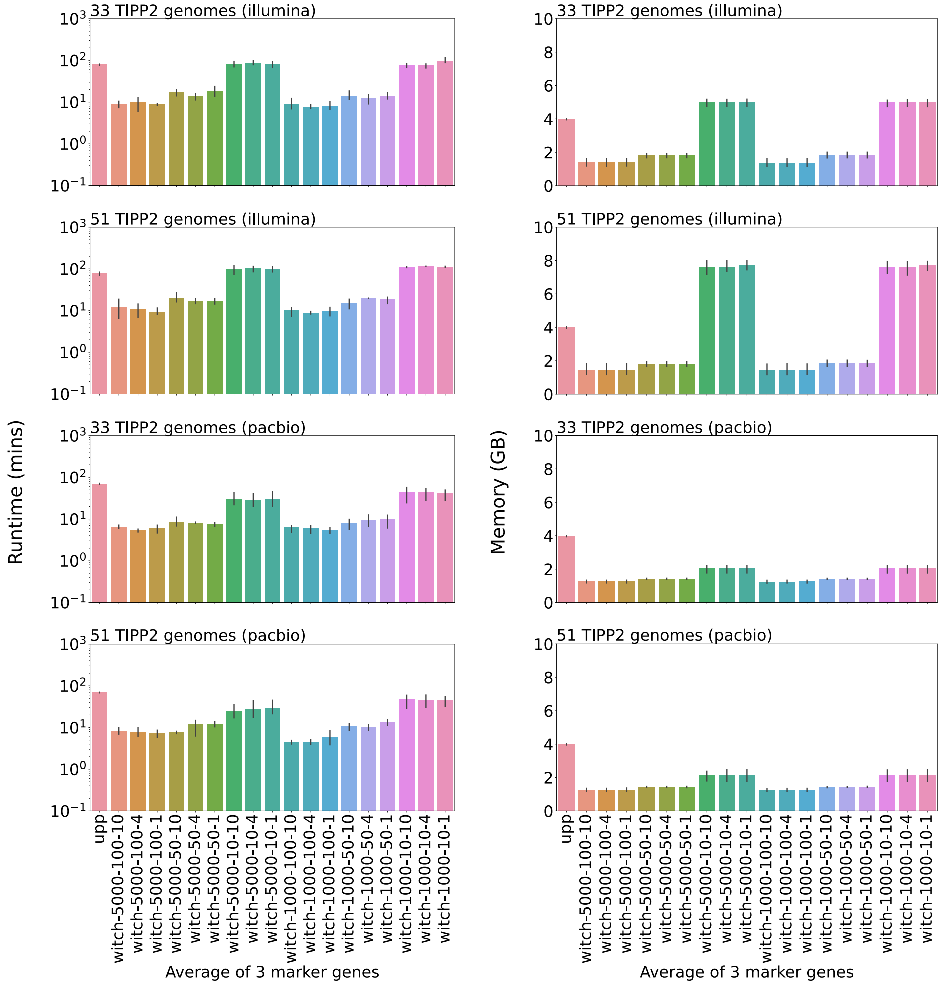

**(a)** Runtime (mins, log-scale). **(b)** Memory (GBs).

**Fig S3.** Runtime usage in minutes (left, log-scale) and memory usage in GBs (right) of identifying Illumina and PacBio reads from two TIPP3 datasets with 33 and 51 genomes, by UPP and different WITCH variants. Results are averaged over three marker genes (RplO, RpsK, and RpsL). Runtime for each variant includes read alignment and placement phases.

#### S5.2 Placing reads into marker gene taxonomies

After selecting the read alignment method, we explored different phylogenetic placement methods that place aligned reads into a target tree (i.e., the taxonomy of each marker gene), including SCAMPP [12], BSCAMPP [11], and pplacer with the `taxtastic` package (referred to as pplacer-taxtastic) [4,13]. Both SCAMPP and BSCAMPP are divide-and-conquer methods that decompose the input tree into smaller subtrees and select suitable subtrees for query placements, using pplacer (SCAMPP) or EPA-ng [14] (BSCAMPP). We varied subtree sizes ranging from 1000 to 5000 leaves for SCAMPP and BSCAMPP, and support values ranging from 50% to 99% for all three methods (see main text for the definition of a support value). We then examined the taxonomic identification and abundance profiling accuracy of each method variant.

Overall, pplacer-taxtastic with a support value of 90% provides the best accuracy, while BSCAMPP with a subtree of size 1000 and a support value of 95% gives good accuracy and the fastest runtime. Hence, we selected pplacer with the `taxtastic` package (90% support value) as the read placement method for TIPP3. Full results for the placement benchmarks can be found below.

##### S5.2.1 Batch-SCAMPP

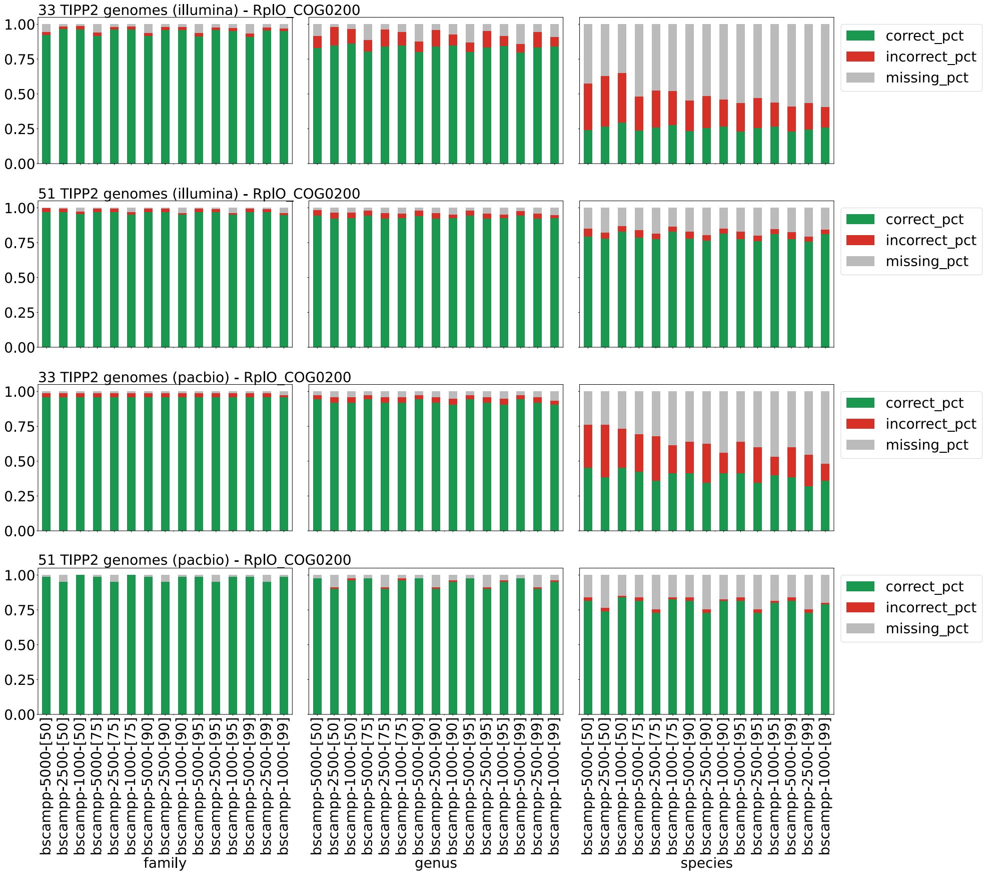

**Fig S4.** Taxonomic identification accuracy of different batch-SCAMPP variants for query placement in TIPP3 on marker gene RplO_COG0200, for Illumina and PacBio reads from two TIPP2 datasets with 33 and 51 genomes (Family, Genus, and Species levels). Query reads are aligned with WITCH. Batch-SCAMPP variants are named **bscampp-X-[Y]**, where $X$ is the subtree size ($X = \{1000, 2500, 5000\}$) and $Y$ is the support value ($Y = \{50, 75, 90, 95, 99\}$). "correct_pct" denotes the fraction of correctly identified reads (at a taxonomic level), "incorrect_pct" denotes the fraction of incorrectly identified reads, and "missing_pct" denotes the fraction of not-identified reads. The three fractions sum to 1 for each bar.

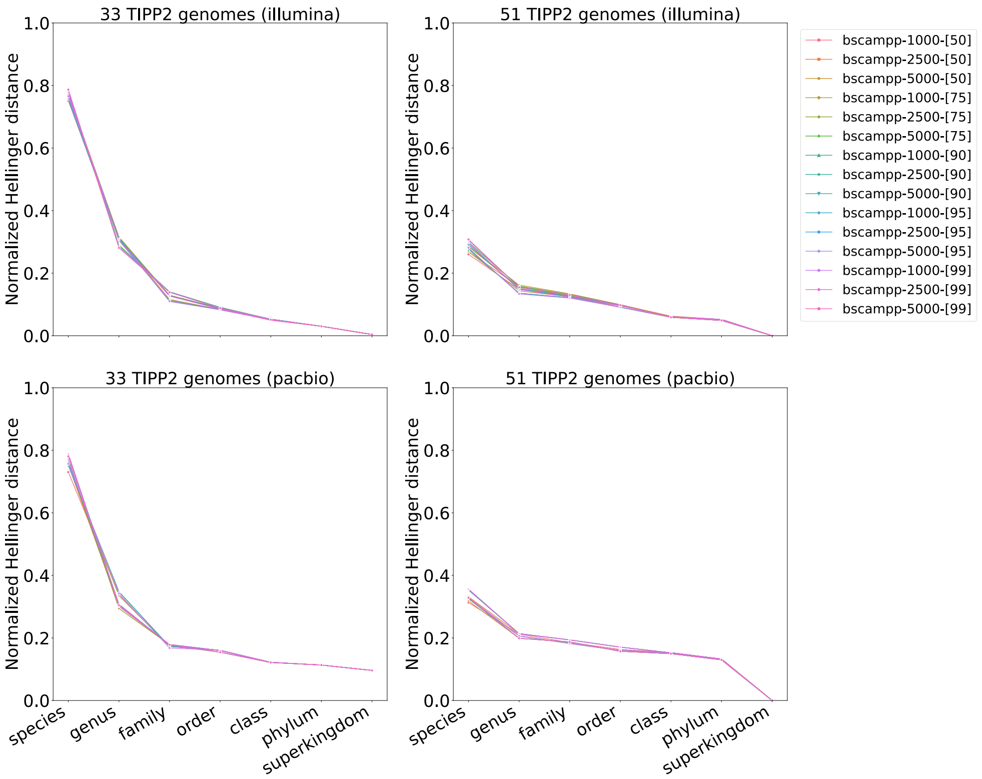

**Fig S5.** Abundance profile of different batch-SCAMPP variants for query placement in TIPP3, for Illumina and PacBio reads from two TIPP2 datasets with 33 and 51 genomes. Query reads are aligned with WITCH. The abundance profile is computed as the normalized Hellinger distance between the estimated and reference profiles using three marker genes, RplO, RpsK, and RpsL.

##### S5.2.2 SCAMPP

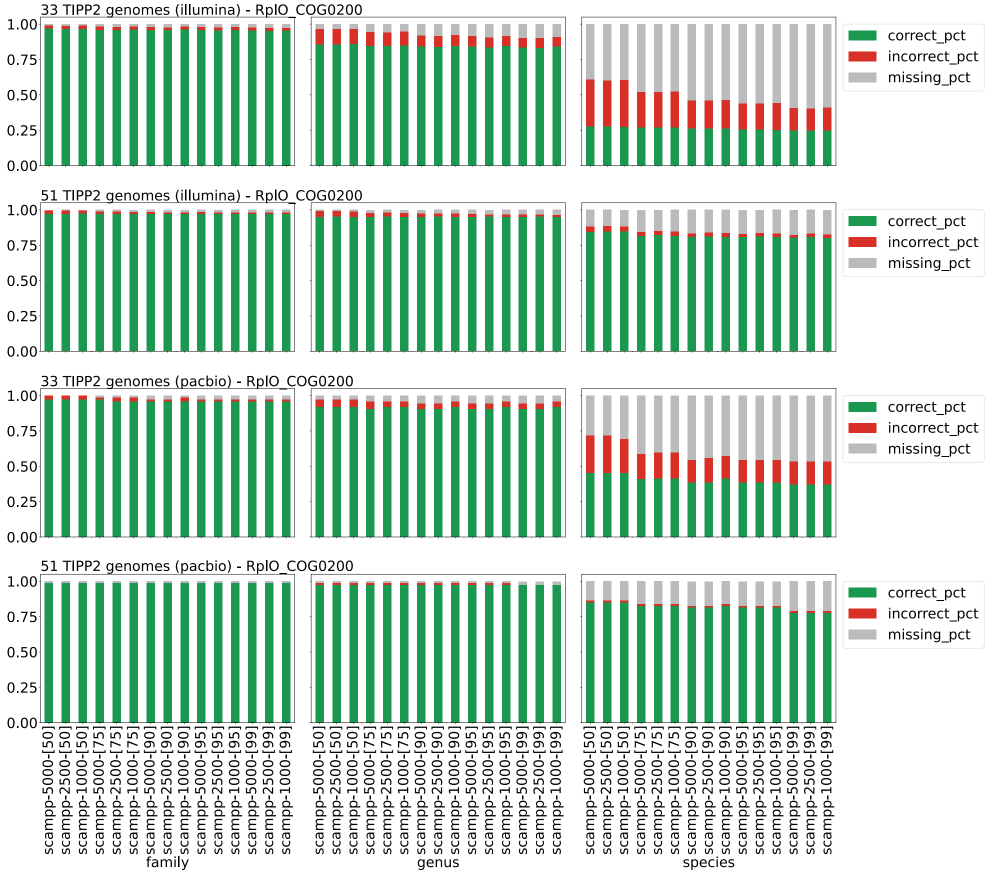

**Fig S6.** Taxonomic identification accuracy of different SCAMPP variants for query placement in TIPP3 on marker gene RplO_COG0200, for Illumina and PacBio reads from two TIPP2 datasets with 33 and 51 genomes (Family, Genus, and Species levels). Query reads are aligned with WITCH. SCAMPP variants are named **scampp-X-[Y]**, where $X$ is the subtree size ($X = \{1000, 2500, 5000\}$) and $Y$ is the support value ($Y = \{50, 75, 90, 95, 99\}$). "correct_pct" denotes the fraction of correctly identified reads (at a taxonomic level), "incorrect_pct" denotes the fraction of incorrectly identified reads, and "missing_pct" denotes the fraction of not-identified reads. The three fractions sum to 1 for each bar.

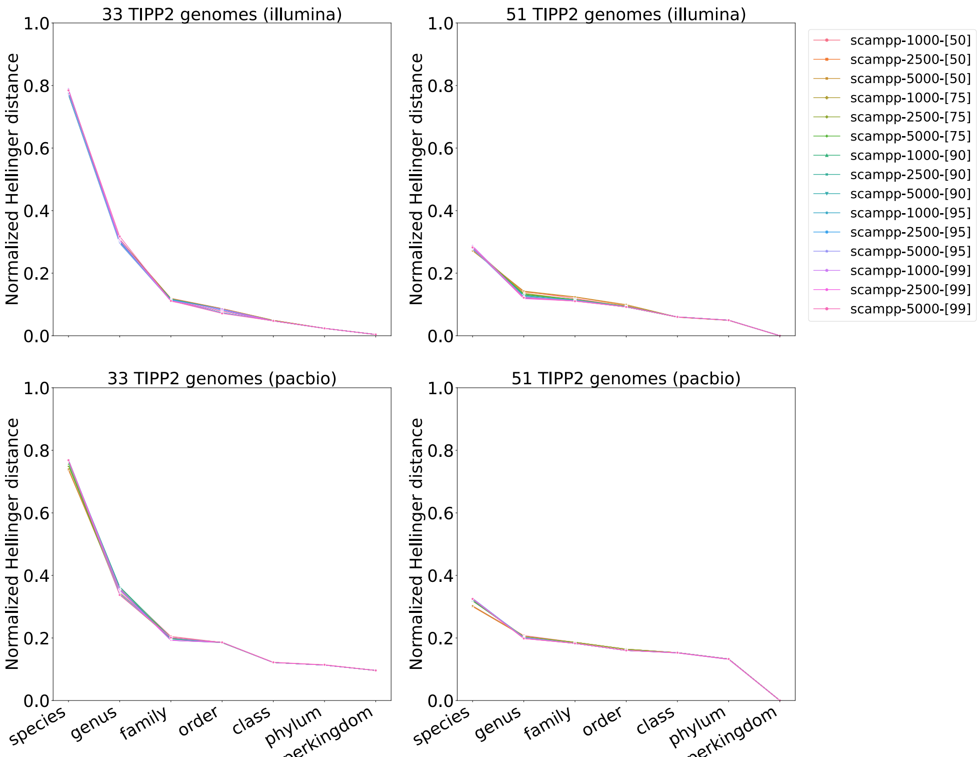

**Fig S7.** Abundance profile of different SCAMPP variants for query placement in TIPP3, for Illumina and PacBio reads from two TIPP2 datasets with 33 and 51 genomes. Query reads are aligned with WITCH. The abundance profile is computed as the normalized Hellinger distance between the estimated and reference profiles using three marker genes, RplO, RpsK, and RpsL.

##### S5.2.3 pplacer-taxtastic

297

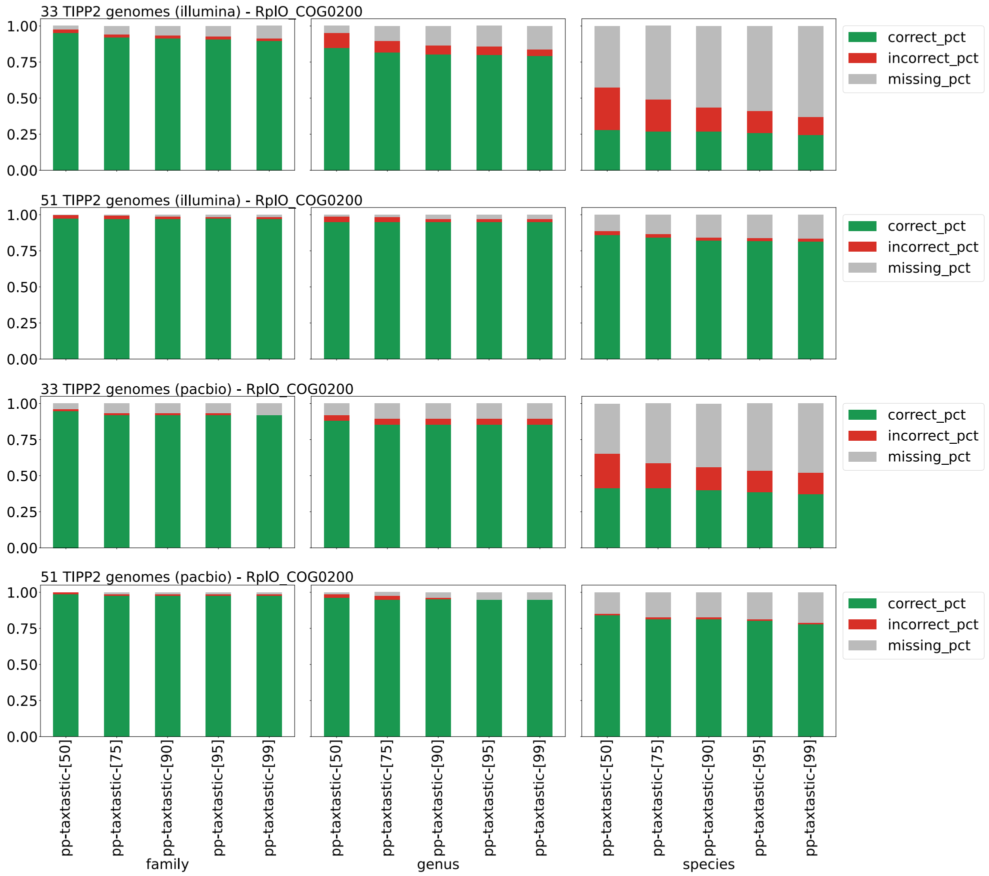

**Fig S8.** Taxonomic identification accuracy of different pplacer-taxtastic variants for query placement in TIPP3 on marker gene RplO_COG0200, for Illumina and PacBio reads from two TIPP2 datasets with 33 and 51 genomes (Family, Genus, and Species levels). Query reads are aligned with WITCH. pplacer-taxtastic variants are named **pp-taxtastic-[Y]**, where $Y$ is the support value ($Y = \{50, 75, 90, 95, 99\}$). "correct_pct" denotes the fraction of correctly identified reads (at a taxonomic level), "incorrect_pct" denotes the fraction of incorrectly identified reads, and "missing_pct" denotes the fraction of not-identified reads. The three fractions sum to 1 for each bar.

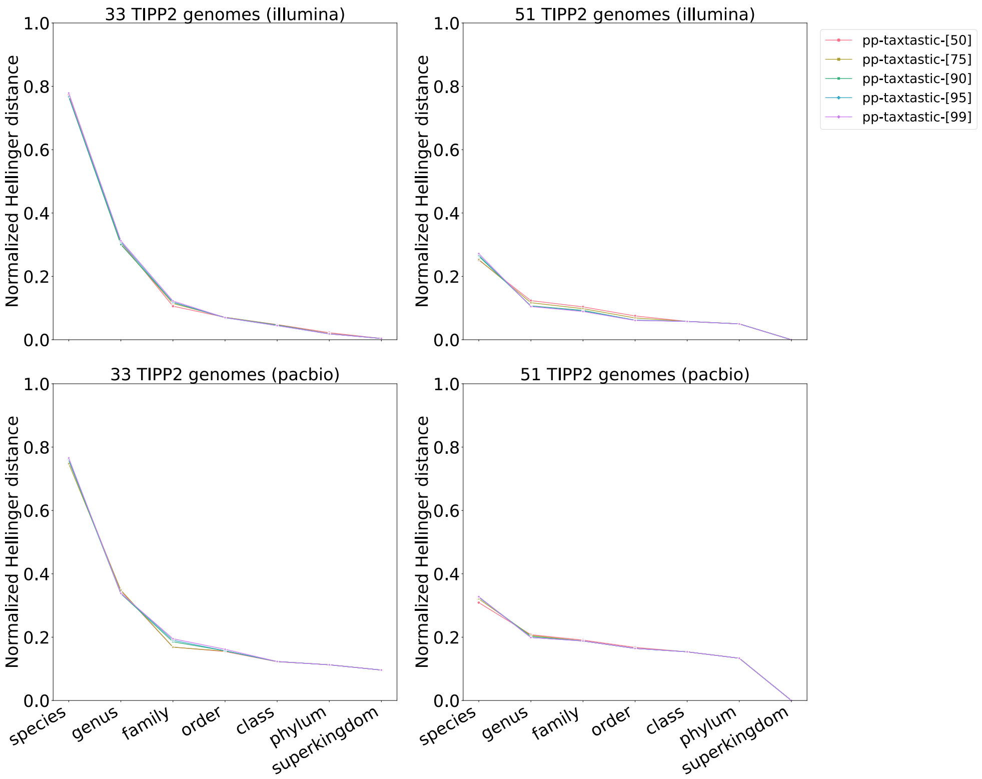

**Fig S9.** Abundance profile of different pplacer-taxtastic variants for query placement in TIPP3, for Illumina and PacBio reads from two TIPP2 datasets with 33 and 51 genomes. Query reads are aligned with WITCH. The abundance profile is computed as the normalized Hellinger distance between the estimated and reference profiles using three marker genes, RplO, RpsK, and RpsL.

##### S5.2.4 Comparison of the three placement methods

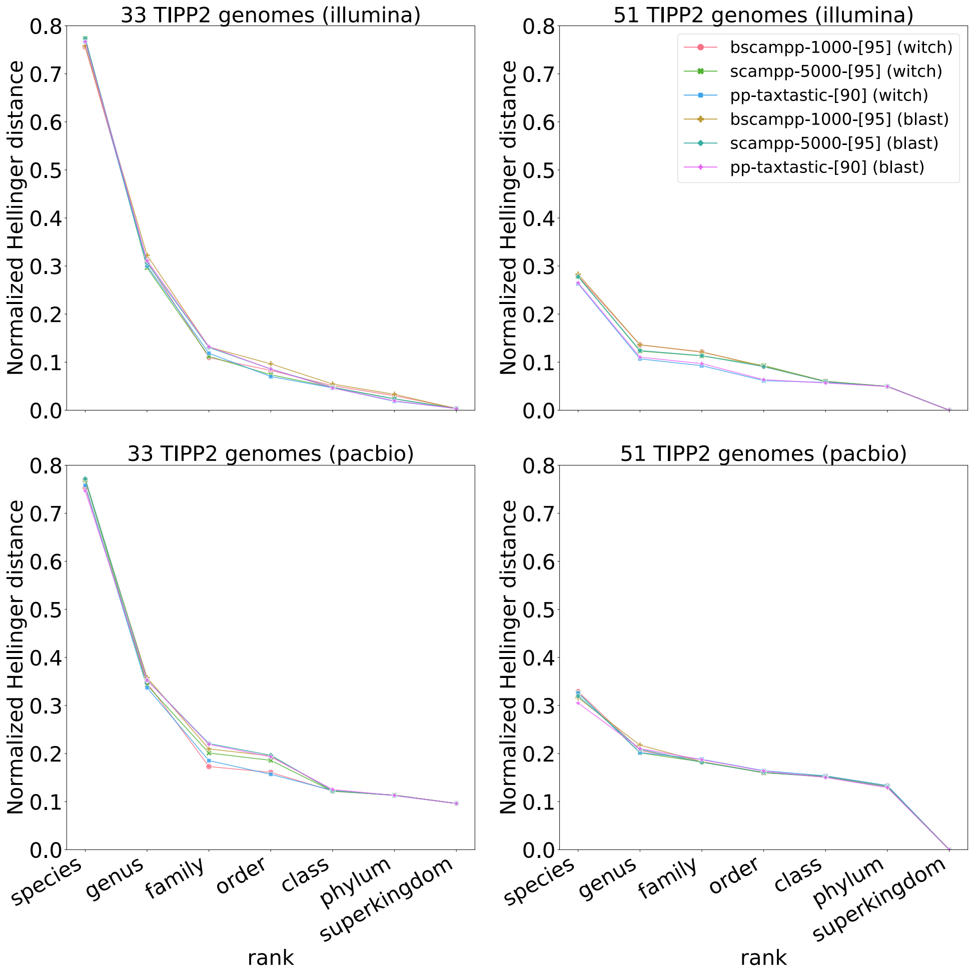

**Fig S10.** Abundance profile of the best variants of batch-SCAMPP, SCAMPP, and pplacer-taxtastic for Illumina and PacBio reads from two TIPP2 datasets with 33 and 51 genomes. Query reads are aligned with either WITCH or BLAST (denoted after the name of each method). The abundance profile is computed as the normalized Hellinger distance between the estimated and reference profiles using three marker genes, RplO, RpsK, and RpsL.

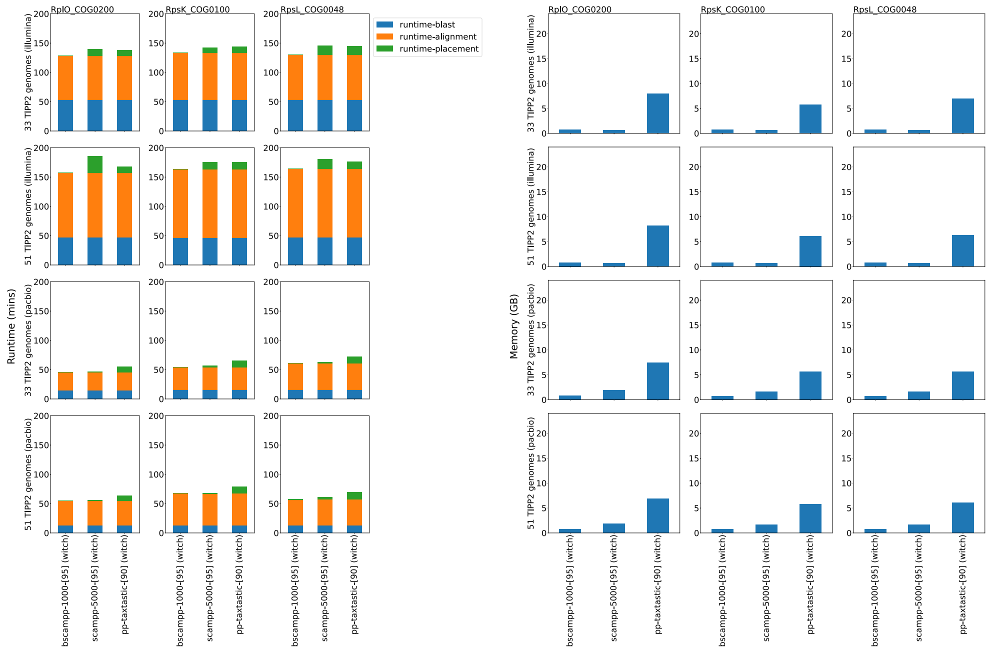

**(a)** Runtime (mins).

**(b)** Memory (GBs).

**Fig S11.** Runtime usage in minutes (left) and memory usage in GBs (right), using WITCH alignment, of the best variants of BSCAMPP, SCAMPP, and pplacer-taxtastic identifying Illumina and PacBio reads from two TIPP2 datasets (33 and 51 genomes) assigned to three marker genes, RplO, RpsK, and RpsL. Runtime for blast, alignment (by WITCH), and placement are shown. “runtime-alignment” denotes the time to obtain the read alignment by WITCH assigned to each marker gene. “runtime-placement” denotes the total time used to obtain placements of query reads assigned to each marker gene.

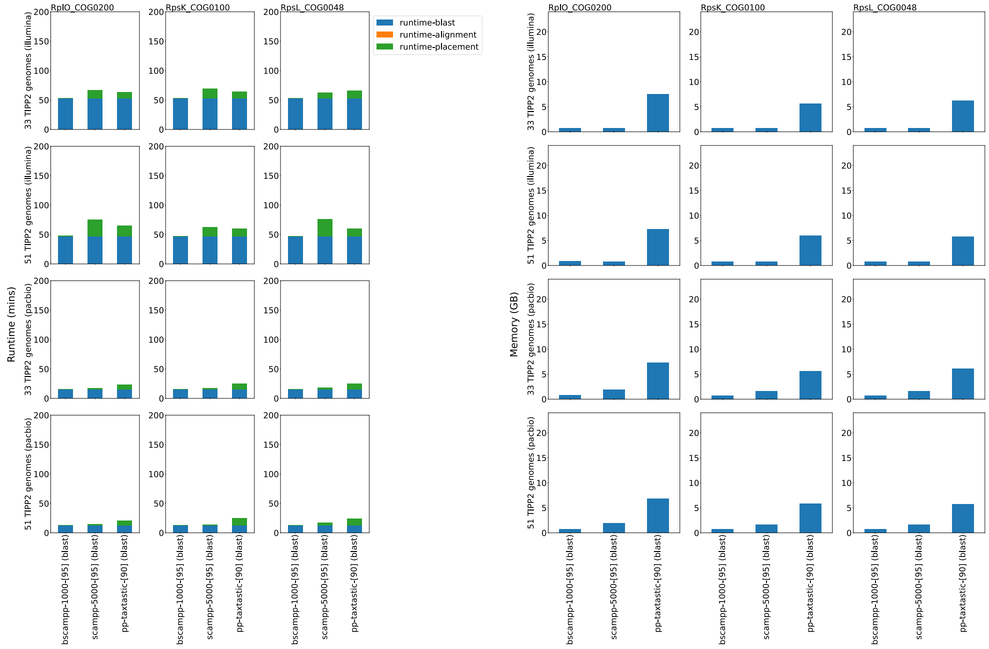

**(a)** Runtime (mins).   **(b)** Memory (GBs).

**Fig S12.** Runtime usage in minutes (left) and memory usage in GBs (right), using BLAST alignment, of the best variants of BSCAMPP, SCAMPP, and pplacer-taxtastic identifying Illumina and PacBio reads from two TIPP2 datasets (33 and 51 genomes) assigned to three marker genes, RplO, RpsK, and RpsL. Runtime for blast, alignment (by BLAST), and placement are shown. "runtime-alignment" denotes the time to post-process the BLAST output to obtain best-scored pairwise alignments (to reference genomes) of binned query reads. "runtime-placement" denotes the total time used to obtain placements of query reads assigned to each marker gene.

#### S5.3 Marker gene selection

So far, we have only looked at three marker genes (RplO, RpsK, and RpsL) for abundance profiling. Here, we examined the accuracy of TIPP3 on each of the 40 marker genes when profiling Illumina reads from the TIPP2 dataset with 51 genomes. We then sorted the marker genes by their averaged normalized Hellinger distance across all taxonomic levels (Figure S13). Full results can be found in this sub-section.

We observed a wide gap between the best and the worst marker genes in terms of their profiling accuracy, particularly on the genus level and up. When using a subset of marker genes for aggregated abundance profiling, we found that excluding either the largest two marker genes (FtsY and RpoB, ranked by marker gene alignment sizes) or the bottom 20 marker genes generally resulted in higher profiling accuracy compared to using all 40 marker genes or just three marker genes (Figure S14). Hence, we removed FtsY and RpoB genes from the TIPP3 reference package and used the remaining 38 marker genes for the following experiments.

##### S5.3.1 Abundance profile on each marker gene

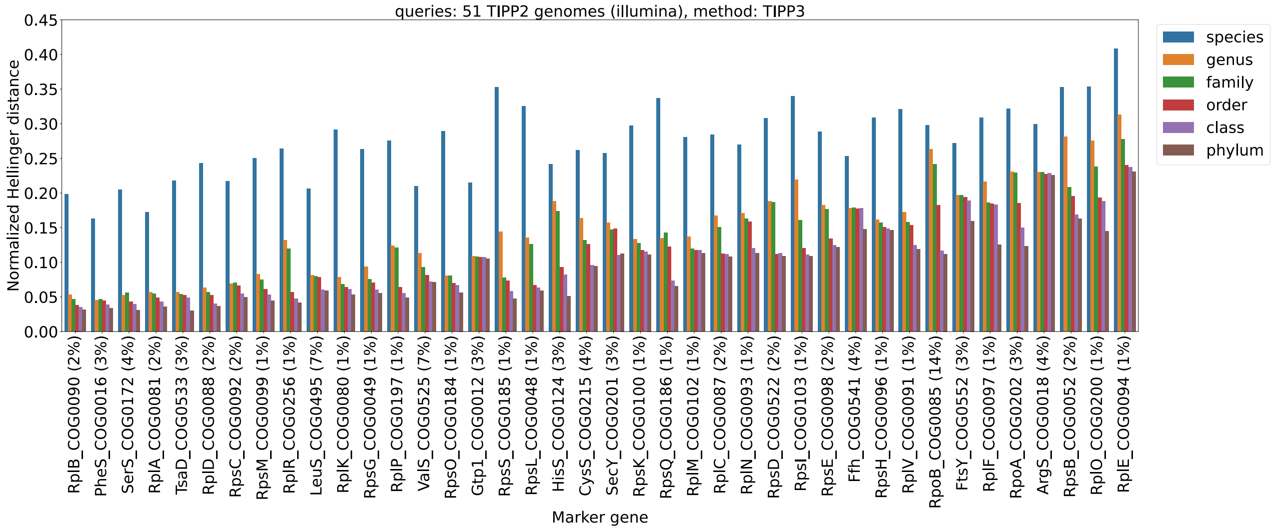

**Fig S13.** Individual marker gene abundance profile accuracy for TIPP3 on each taxonomic level. The query reads used are Illumina reads and generated from the TIPP2 dataset with 51 genomes. The x-axis denotes different marker genes and taxonomic levels (excluding superkingdom). The marker genes are sorted from left to right in order of average ranking of normalized Hellinger distance (i.e., the lower average normalized Hellinger distance a marker gene has, the closer it is to left on the x-axis). The "%" sign for each marker gene denotes the percentage of query reads assigned to that marker gene.

##### S5.3.2 Abundance profile using different sets of marker genes

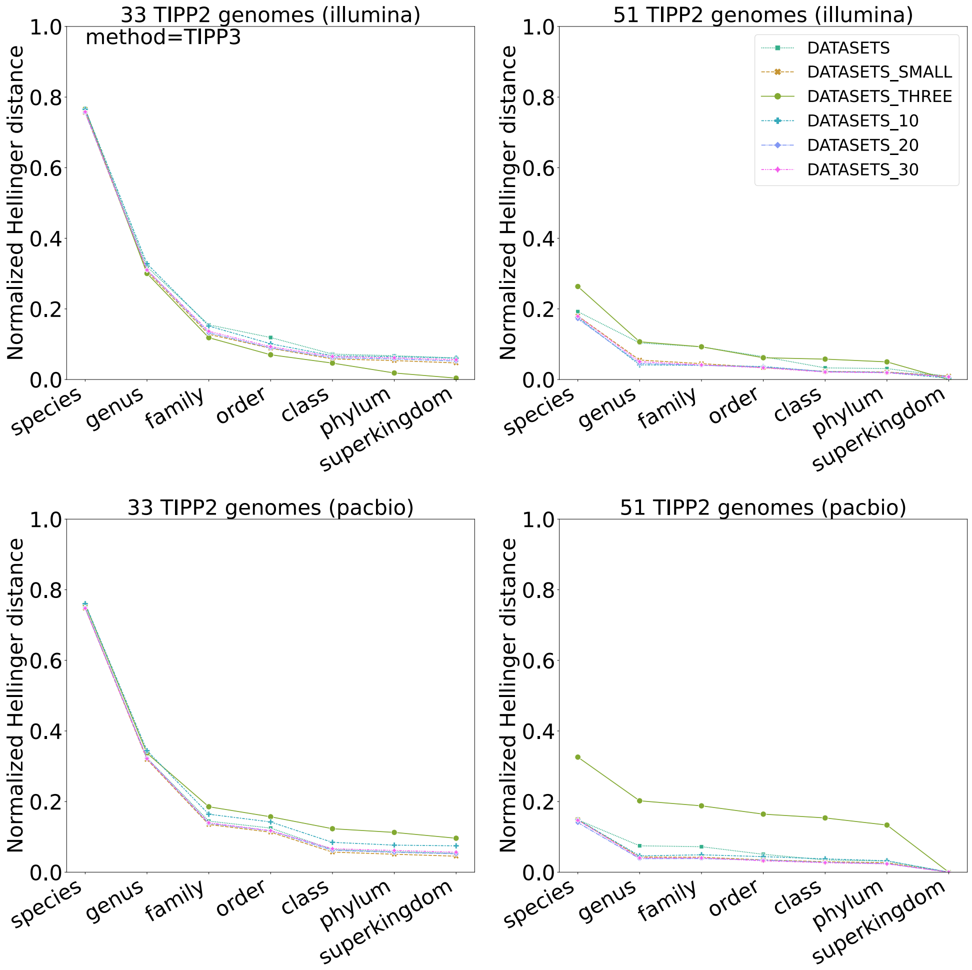

**Fig S14.** Abundance profiles of TIPP3 using different sets of marker genes. DATASETS denotes using all marker genes, DATASETS_SMALL for using all except RpoB and FtsY, DATASETS_THREE for using only RplO, RpsK, and RpsL, DATASETS_10 for using the top 10 marker genes denoted in Figure S13, DATASETS_20 for using the top 20, and DATASETS_30 for using the top 30 marker genes. Reads are either Illumina or PacBio generated from two TIPP2 datasets with 33 and 51 genomes.

### S6 Additional Results for Experiment 3: Evaluation of TIPP3 for abundance profiling

#### S6.1 Abundance profile comparison for all methods

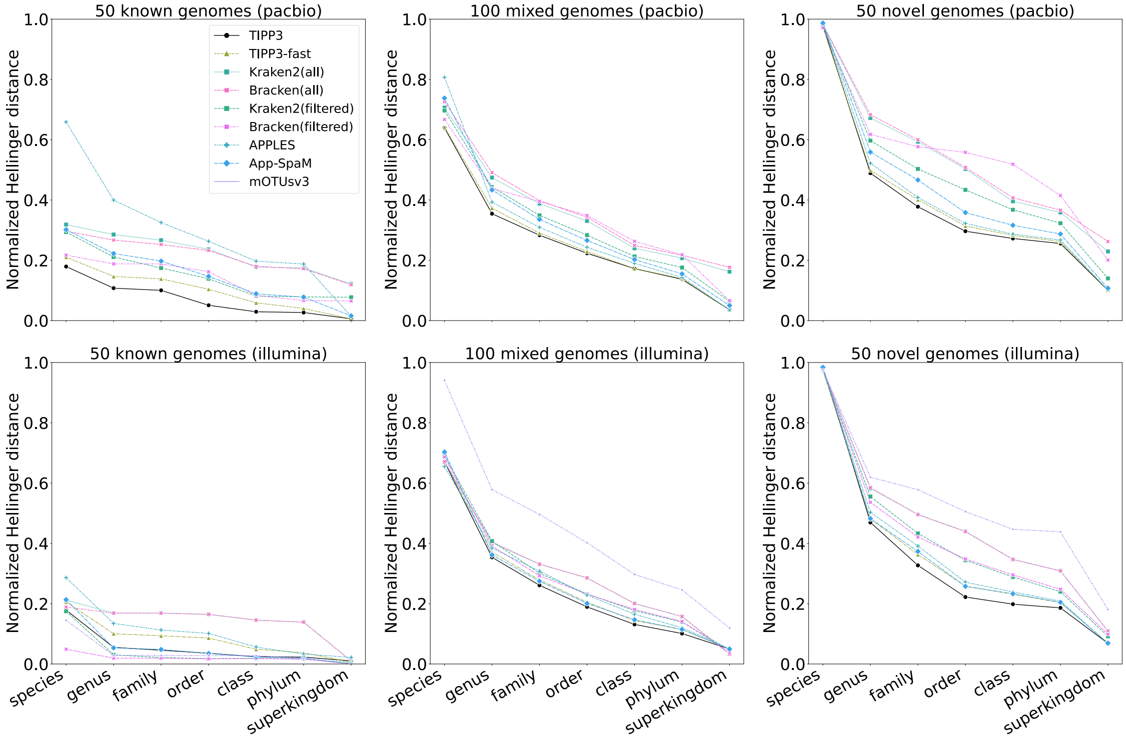

**Fig S15.** Abundance profiling accuracy by normalized Hellinger distance (lower means more accurate) of all methods/pipelines for PacBio and Illumina simulated reads from 50 known, 100 mixed, and 50 novel genomes. Abundance profiles are computed using 38 marker genes for TIPP3, TIPP3-fast, Bracken(filtered), Kraken2(filtered), APPLES, and App-SpaM. For PacBio read datasets, mOTUsv3 did not produce any classification and thus is not shown.

#### S6.2 Detailed evaluation on species and genus abundances

Genus- and Species-specific abundance estimation errors are shown. Species-specific errors are not shown for 50 novel genomes, since no methods should be able to produce any non-zero profile for the species by definition. The genus and species names and their corresponding reference abundances of the 50 known genomes are shown in Tables S1.

**Table S1.** Taxon names and reference abundances of the 50 known genomes, sorted by alphabetical order.

| taxon name | abundance |
|---|---|
| Acholeplasma laidlawii | 0.02 |
| Acidobacterium capsulatum | 0.02 |
| Actinosynnema mirum | 0.02 |
| Akkermansia muciniphila | 0.02 |
| Alcanivorax borkumensis | 0.02 |
| Aminobacterium colombiense | 0.02 |

| | |
|---|---|
| Bernardetia litoralis | 0.02 |
| Beutenbergia cavernae | 0.02 |
| Brucella melitensis | 0.02 |
| Caldisericum exile | 0.02 |
| Cellulophaga lytica | 0.02 |
| Chlamydia psittaci | 0.02 |
| Chlamydia trachomatis | 0.02 |
| Chlorobium limicola | 0.02 |
| Chloroflexus aurantiacus | 0.02 |
| Clavibacter michiganensis | 0.02 |
| Clostridium botulinum | 0.02 |
| Coprothermobacter proteolyticus | 0.02 |
| Dehalococcoides mccartyi | 0.02 |
| Desulfofarcimen acetoxidans | 0.02 |
| Dictyoglomus thermophilum | 0.02 |
| Dinoroseobacter shibae | 0.02 |
| Elusimicrobium minutum | 0.02 |
| Fervidobacterium pennivorans | 0.02 |
| Filifactor alocis | 0.02 |
| Flexistipes sinusarabici | 0.02 |
| Gemmatimonas aurantiaca | 0.02 |
| Gloeobacter kilaueensis | 0.02 |
| Listeria monocytogenes | 0.02 |
| Melioribacter roseus | 0.02 |
| Mycoplasma gallisepticum | 0.02 |
| Neisseria meningitidis | 0.02 |
| Oceanithermus profundus | 0.02 |
| Paludibacter propionicigenes | 0.02 |
| Parachlamydia acanthamoebae | 0.02 |
| Persephonella marina | 0.02 |
| Planctopirus limnophila | 0.02 |
| Pseudopedobacter saltans | 0.02 |
| Roseiflexus castenholzii | 0.02 |
| Salmonella enterica | 0.02 |
| Segniliparus rotundus | 0.02 |
| Simkania negevensis | 0.02 |
| Sphaerobacter thermophilus | 0.02 |
| Streptobacillus moniliformis | 0.02 |
| Streptococcus pneumoniae | 0.02 |
| Tannerella forsythia | 0.02 |
| Thermomicrobium roseum | 0.02 |
| Turneriella parva | 0.02 |
| Ureaplasma urealyticum | 0.02 |
| Waddlia chondrophila | 0.02 |

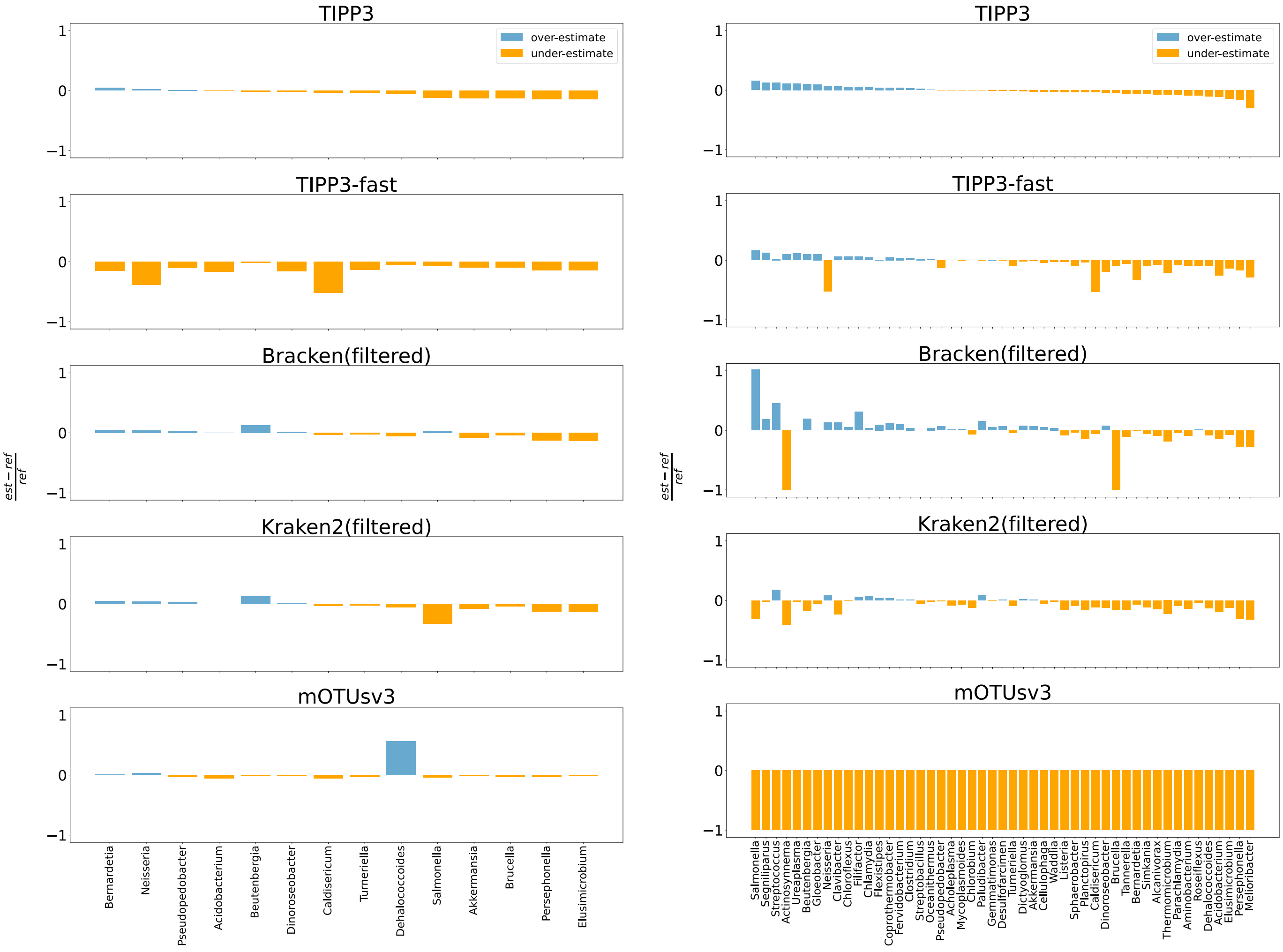

**(a)** For Illumina reads from 50 known genomes, genus level. **(b)** For PacBio reads from 50 known genomes, genus level.

**Fig S16.** Genus-specific abundance estimation error for Illumina (left) and PacBio (right) reads of 50 known genomes of TIPP3, Bracken(filtered), Bracken(all), Kraken2(filtered), and Kraken2(all). The estimation error of a taxon is computed as the fractional difference between its estimated and reference compositions, shown on the Y-axis. Taxons are sorted by the estimation errors by TIPP3 in ascending order. A taxon group is shown if and only if it is present in the reference, and any of the methods has an estimation error ±10%.

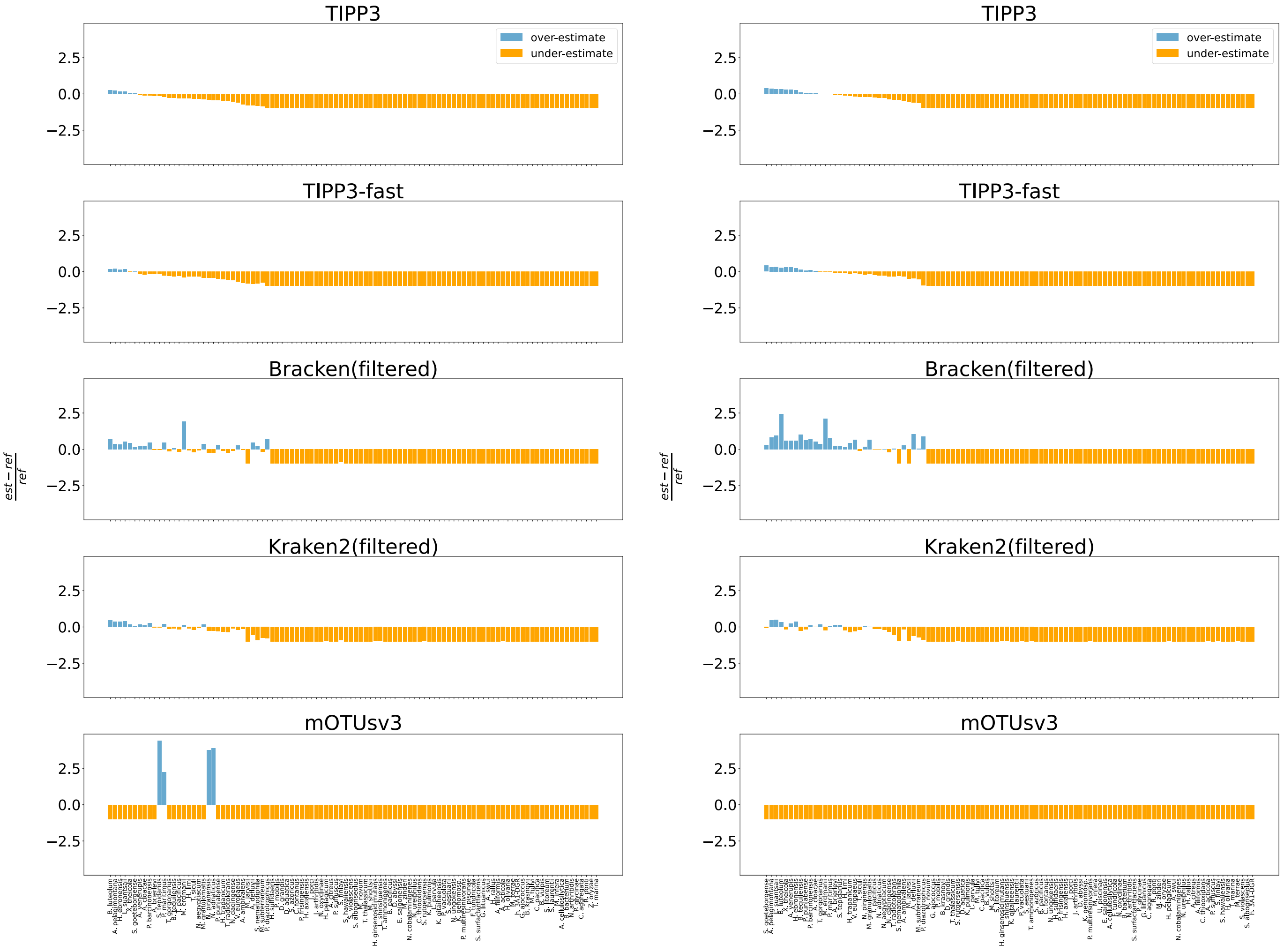

**(a)** For Illumina reads from 100 mixed genomes, species level. **(b)** For PacBio reads from 100 mixed genomes, species level.

**Fig S17.** Species-specific abundance estimation error for Illumina (left) and PacBio (right) reads of 100 mixed genomes of TIPP3, Bracken(filtered), Bracken(all), Kraken2(filtered), and Kraken2(all). The estimation error of a taxon is computed as the fractional difference between its estimated and reference compositions, shown on the Y-axis. Taxons are sorted by the estimation errors by TIPP3 in ascending order. A taxon group is shown if and only if it is present in the reference, and any of the methods has an estimation error ±10%.

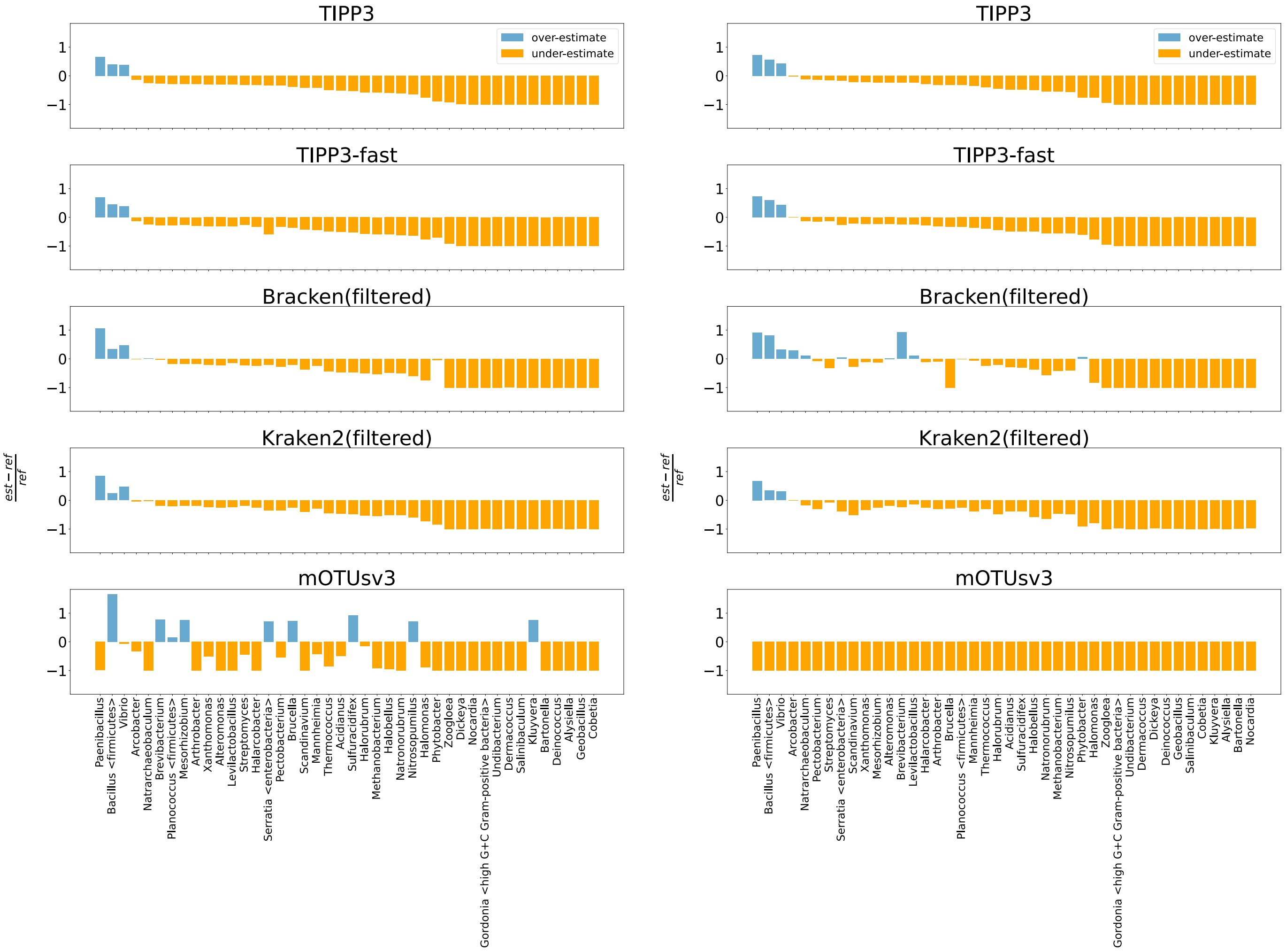

**(a)** For Illumina reads from 100 mixed genomes, genus level. **(b)** For PacBio reads from 100 mixed genomes, genus level.

**Fig S18.** Genus-specific abundance estimation error for Illumina (left) and PacBio (right) reads of 100 mixed genomes of TIPP3, Bracken(filtered), Bracken(all), Kraken2(filtered), and Kraken2(all). The estimation error of a taxon is computed as the fractional difference between its estimated and reference compositions, shown on the Y-axis. Taxons are sorted by the estimation errors by TIPP3 in ascending order. A taxon group is shown if and only if it is present in the reference, and any of the methods has an estimation error ±10%.

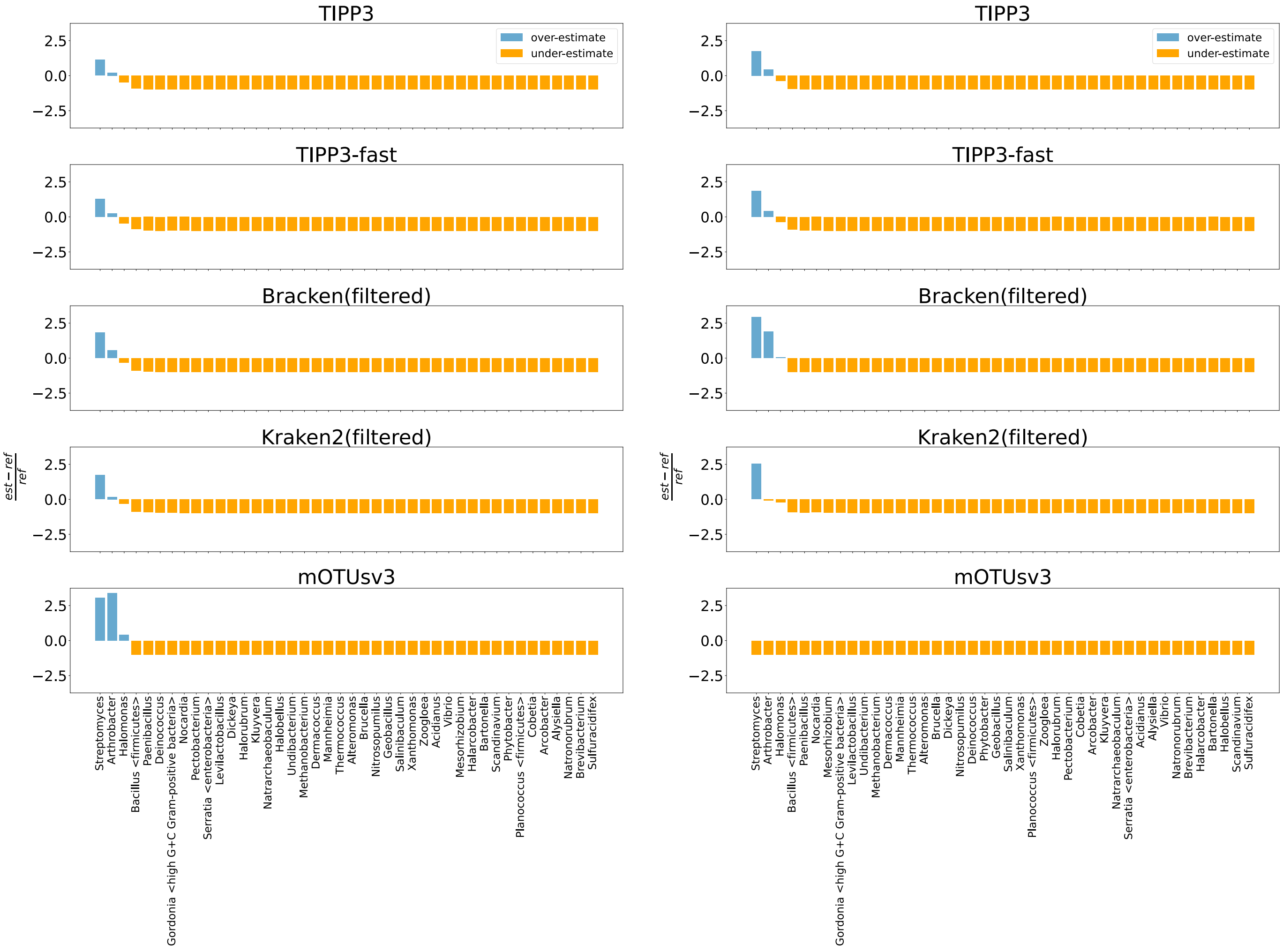

**(a)** For Illumina reads from 50 novel genomes, genus level. **(b)** For PacBio reads from 50 novel genomes, genus level.

**Fig S19.** Genus-specific abundance estimation error for Illumina (left) and PacBio (right) reads of 50 novel genomes of TIPP3, Bracken(filtered), Bracken(all), Kraken2(filtered), and Kraken2(all). The estimation error of a taxon is computed as the fractional difference between its estimated and reference compositions, shown on the Y-axis. Taxons are sorted by the estimation errors by TIPP3 in ascending order. A taxon group is shown if and only if it is present in the reference, and any of the methods has an estimation error ±10%.

#### S6.3 Runtime and memory usage

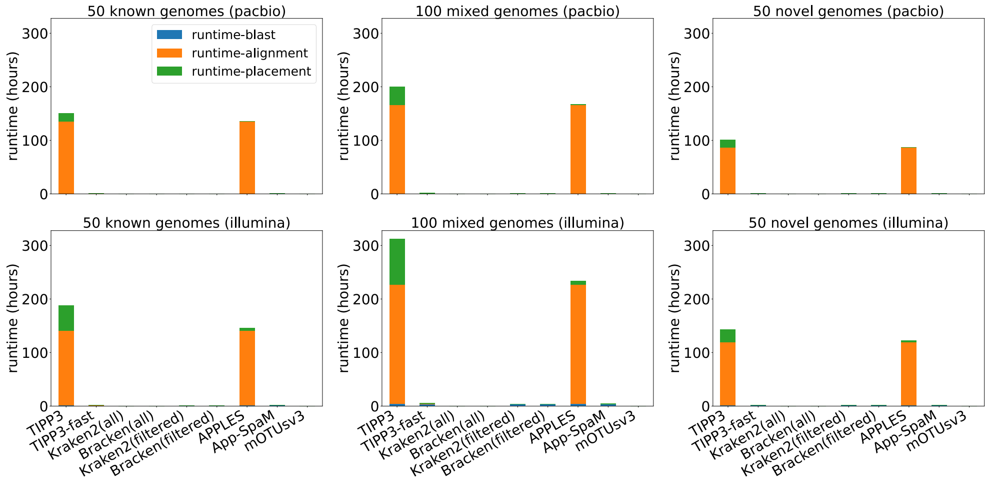

**(a)** Runtime (hours).

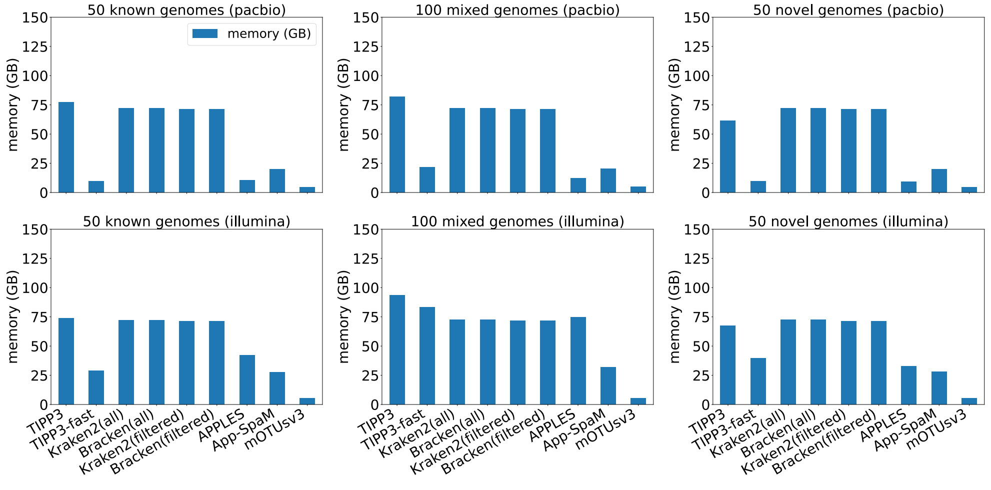

**(b)** Maximum memory usage (GBs).

**Fig S20.** Runtime in hours (top) and maximum memory usage in GBs (bottom) of TIPP3, TIPP3-fast, Kraken2(filtered), Bracken(filtered), mOTUsv3, APPLES, and App-SpaM for PacBio and Illumina simulated reads from 50 known, 100 mixed, and 50 novel genomes.

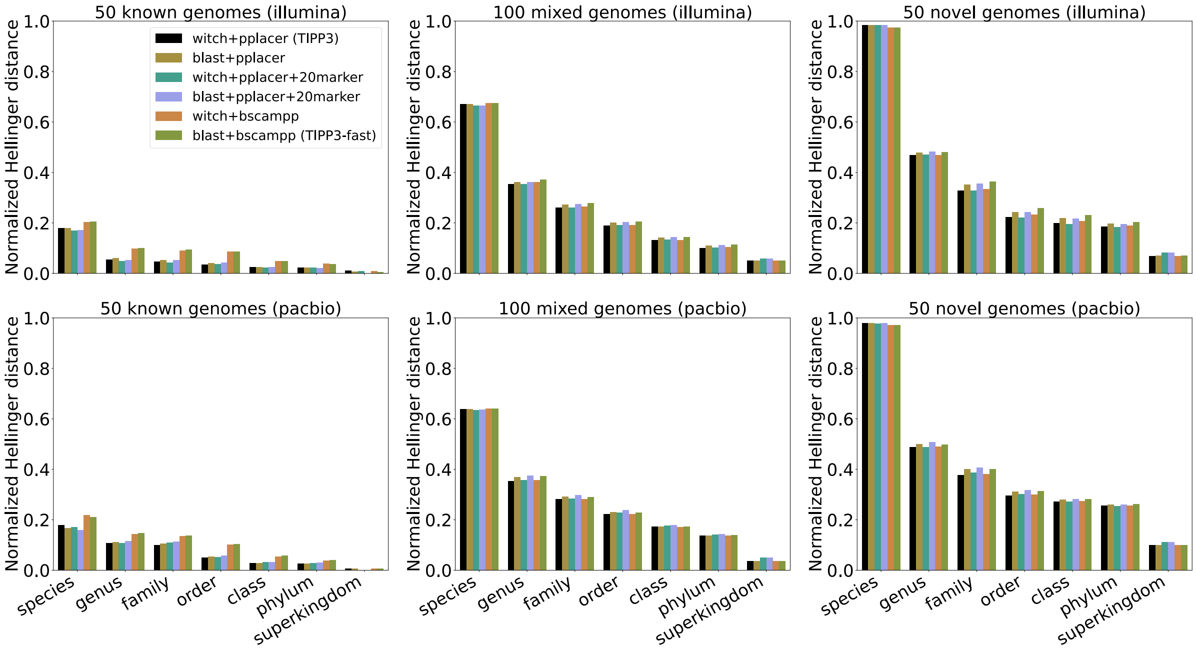

**Fig S21.** Abundance profile accuracy by normalized Hellinger distance of TIPP3 variants for Illumina and PacBio simulated reads from known, mixed, and novel genomes. Each TIPP3 variant is denoted as "`<align method>+<placement method>`" (using 38 marker genes) or "`<align method>+<placement method>+<Xmarker>`" (using $X$ marker genes). For example, TIPP3 uses WITCH for query alignment and pplacer-taxtastic for query placement and is denoted as "`witch+pplacer`".

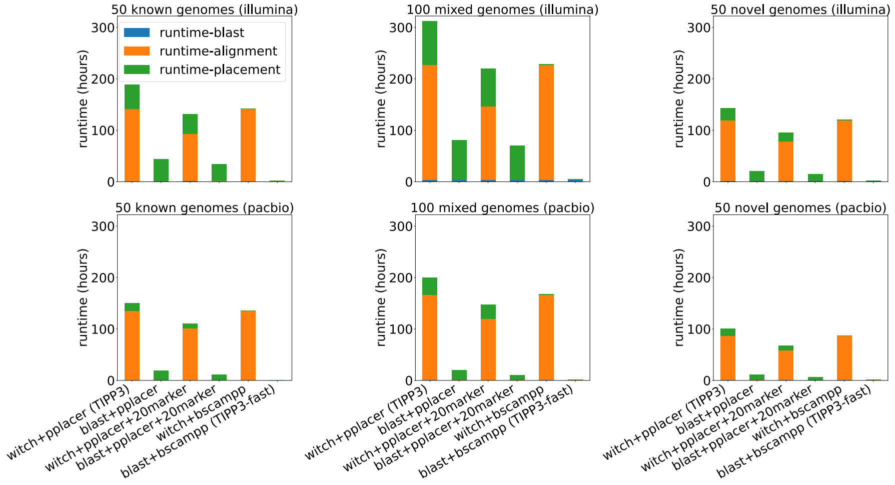

**(a)** Runtime (log-scale, hours).

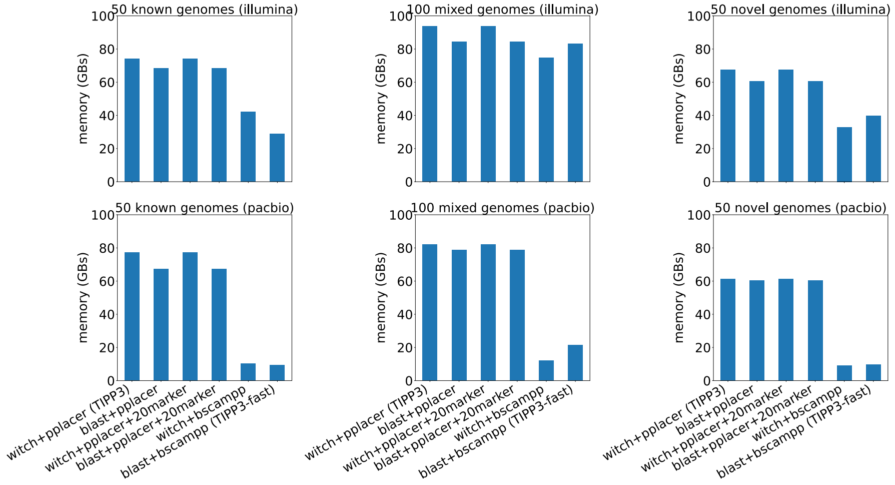

**(b)** Maximum memory usage (GBs).

**Fig S22.** Runtime in hours (top) and peak memory usage in GBs (bottom) of all benchmarked methods/pipelines for Illumina and PacBio simulated reads from known, mixed, and novel genomes.

**Table S2.** TIPP3 and TIPP3-fast runtime comparison in hours and the corresponding speedup by TIPP3-fast to TIPP3, on the six testing datasets; average results are shown in the last row.

| | TIPP3 (hours) | TIPP3-fast (hours) | speed up |
|---|---|---|---|
| 50 known genomes (PacBio) | 150.0 | 1.0 | 146.1 |
| 50 known genomes (Illumina) | 188.4 | 2.0 | 93.7 |
| 100 mixed genomes (PacBio) | 200.0 | 1.7 | 115.8 |
| 100 mixed genomes (Illumina) | 312.1 | 5.2 | 60.0 |
| 50 novel genomes (PacBio) | 101.3 | 1.3 | 80.4 |
| 50 novel genomes (Illumina) | 143.0 | 2.4 | 58.6 |
| Average | 182.5 | 2.3 | 92.4 |
